# Near-atomic Structure of the Cytoplasmic Ring of the *Xenopus laevis* Nuclear Pore Complex

**DOI:** 10.1101/2022.02.14.480321

**Authors:** Xuechen Zhu, Gaoxingyu Huang, Chao Zeng, Xiechao Zhan, Ke Liang, Yanyu Zhao, Pan Wang, Qifan Wang, Qiang Zhou, Qinghua Tao, Minhao Liu, Jianlin Lei, Chuangye Yan, Yigong Shi

## Abstract

The nuclear pore complex (NPC) mediates nucleocytoplasmic shuttling. Here we present single-particle cryo-EM structure of the cytoplasmic ring (CR) from the *Xenopus laevis* NPC at 4.1-4.7 Å resolutions. The structure of an N-terminal domain of Nup358 was resolved at 3.0 Å, facilitating identification of five Nup358 molecules in each CR subunit. Aside from unveiling the assembly details of the two Y-shaped multicomponent complexes (Y complexes) in each CR subunit, the improved resolutions reveal the C-terminal fragment of Nup160 to be an organizing center at the vertex of each Y complex. Our structures show that the scaffold of a CR subunit comprises five Nup358, two Nup205 and two Nup93 molecules in addition to the previously characterized Y complexes.

**One-Sentence Summary:** Improved resolutions of the cytoplasmic ring (CR) of the *Xenopus laevis* nuclear pore complex reveal that five Nup358 molecules, together with two copies of interweaved Nup205, Nup93 and Y complexes, constitute the scaffold of each CR subunit.

---

NPC resides on the nuclear envelope (NE) and mediates nucleocytoplasmic cargo transport (*1-3*). As one of the largest cellular machineries (*4-6*), a vertebrate NPC has a molecular mass of over 100 MDa and consists of multiple cytoplasmic filaments (CF), a cytoplasmic ring (CR), an inner ring (IR), a nuclear ring (NR), a nuclear basket (NB), and a luminal ring (LR) (*5-9*) (Fig. 1A). Each NPC has eight repeating subunits, known as the spokes (*7*). Our present study concerns the structure of the CR constituent in each spoke, referred to as the CR subunit.

**Fig. 1.**
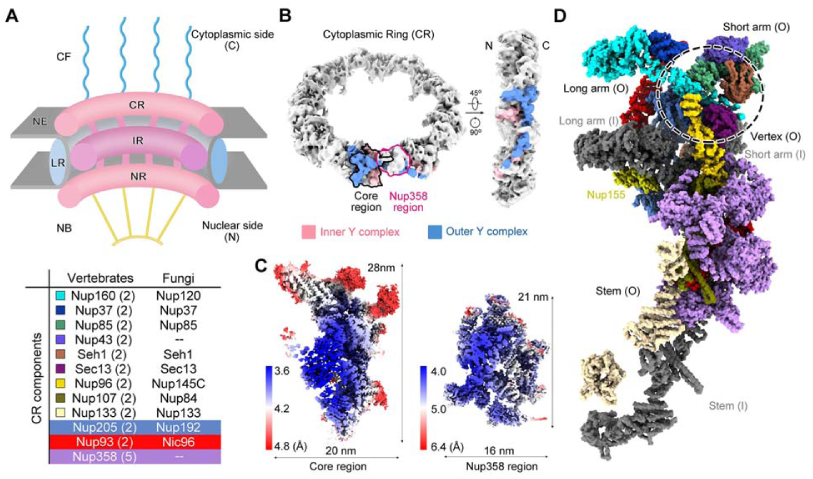
Single-particle cryo-EM analysis of the CR subunit of the *X. laevis* NPC. (*A*) A schematic overview of the architecture of a vertebrate NPC. Shown here is a cut-open side view of a cartoon depicting an NPC embedded in the nuclear envelop (NE). Each NPC, along its cytonuclear axis, comprises the cytoplasmic filaments (CF), cytoplasmic ring (CR), inner ring (IR), and nuclear basket (NB). A luminal ring (LR) that is localized inside the lumen of the NE encompasses the IR. The CR components from vertebrates and fungi are listed in the table below. The copy numbers of CR components in a vertebrate NPC are indicated in brackets. (*B*) EM map of the CR at 22.2 Å resolution. The core region and the Nup358 region in one CR subunit, which are separately calculated as single particles, are indicated by black and magenta contours in the left panel. The densities corresponding to the inner and outer Y complexes in one CR subunit are colored pink and blue, respectively. N and C denote the nuclear side and the cytoplasmic side of the CR ring shown on the right. (*C*) Heat maps for the local resolutions of the core region and the Nup358 region that were separately reconstructed to average resolutions of 4.1 Å and 4.7 Å, respectively. The local resolutions were calculated in Relion (*34*) and presented in ChimeraX (*35*). (*D*) Overall structure of a CR subunit from the *X. laevis* NPC. The structure is presented in approximately the same view as the two Y complexes in the right panel of Fig. 1B. Protein components of the outer Y complex and Nup205, Nup93 and Nup358 molecules are colored the same as those annotated in panel A. The inner Y complex is colored dark gray. Nup155, which connects the CR to the IR, is labeled, but not included in the table in panel A.

The CR subunit of the human NPC was reconstructed through sub-tomogram averaging (STA) to the highest resolution of ∼ 15 Å (*10*). Each CR subunit is featured with two Y-shaped multicomponent complexes, known as the inner and the outer Y complexes. The entire CR was shown to use sixteen Y complexes that assemble in a head-to-tail fashion to build the scaffold in the form of the proximal and distal concentric rings constituted by the inner and outer Y complexes, respectively (Fig. 1A, B) (*11, 12*). The tri-partite Y complexes each consist of a short arm (Nup85, Nup43, and Seh1), a long arm (Nup160 and Nup37), and a stem (Nup96, Sec13, Nup107, and Nup133) (*10*).

To achieve higher resolutions for an advanced understanding of the CR assembly, we adapted single particle cryo-EM reconstruction to examine the CR subunit from the *Xenopus laevis* (*X. laevis*) NPC. During data processing, two relatively independent and rigid structural entities stood out, designated as the core region and the Nup358 region (*13*). The core region contains the long arm from the inner Y complex and the short arms from both Y-complexes, and the Nup358 region comprises the stems of the two Y complexes (Fig. 1B). An overall resolution of 5-8 Å was achieved after masked refinement for these two regions, enabling assignment of some components, such as two Nup205 and two Nup358 molecules (*13*). However, reliable docking and modeling requires higher resolution.

In this study, we improved the resolutions of the core and the Nup358 regions to residue level. We also solved a cryo-EM structure of the N-terminal α-helical domain of Nup358 at 3.0 Å resolution. Relying on the improved resolutions and facilitated by AlphaFold structural prediction, we were able to generate a structural model for the CR of the *X. laevis* NPC. In addition to the previously characterized Y complexes, five Nup358, two Nup205 and two Nup93 molecules were identified to be constituents of the CR scaffold. Our current structure expands the molecular mass by 80% compared to the previously reported composite model of the human CR (*12*).

## Overall structure of a CR subunit

To reduce conformational heterogeneity, we spread the opened nuclear envelope onto the grids with minimal force and used the chemical crosslinker glutaraldehyde to stabilize the NPC. For data processing, please refer to Methods, figs. S1-S4, and Table S1 for details of cryo-sample preparation, image acquisition, data processing, and structural modeling and refinement. We previously defined the core and the Nup358 regions in each CR unit based on their spatial arrangement, but were not able to reliably assign them with nucleoporins (Fig. 1B) (*13*). The average resolutions for these two regions have reached 4.1 Å and 4.7 Å, with the highest local resolution of the core region up to 3.8 Å, in our present study (Fig. 1C). It is now feasible to assign 28 nucleoporins into these regions.

Our reconstruction of the core region displays residue-level features (figs. S5 to S10). For the less well resolved periphery of the core region and the Nup358 region, modeling was assisted by our newly resolved structure of Nup358-NTD2 (222-738) (fig. S11 to S15), previous analyses of vertebrate NPC (*13*) and Alphafold predictions (*14*). Together, these data allow reliable adjustment of the X-ray structures of individual human or fungi nucleoporins and assignment of the *X. laevis* amino acid sequences based on sequence alignment among functional orthologs of each nucleoporin (figs. S16 to S26). A portion of the bulky side chains are assigned. Altogether, our final atomic model of the CR subunit includes 19,037 amino acids in 749 α-helices and 380 β-strands that belong to 30 nucleoporins (Fig. 1A,D, table S2 and S3). Except for Nup358, Nup88, Nup155 and Nup98, all protein components in the current model possess two distinct copies in each CR subunit (*13*). For ease of discussion, the copy closer to and away from the central channel will be denoted with -I and -O, respectively. In the following sections, we will focus on previously uncharacterized structural features of the CR for illustration.

### Nup160-CTF at the vertex of the Y complex

The Y complexes have been most extensively characterized, with the majority of their constituents known. Only in the high-quality map of the core region, is it found that the previously unknown C-terminal fragment of Nup160 (Nup160-CTF), comprising helices α38– α47, sits at the center of the vertex, where the short arm, long arm and stem of the Y complex meet (*11, 15*) (Fig. 2, A and B). With this piece of puzzle solved, the organization of the vertex is defined.

**Fig. 2.**
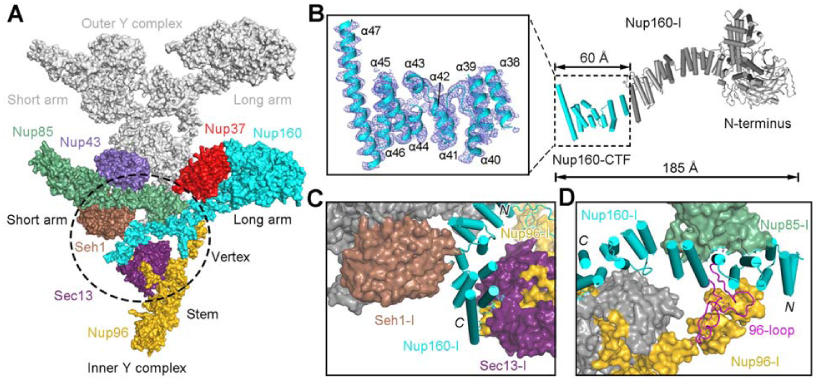
Nup160-CTF is an organizing center for the vertex of the inner Y complex in the CR. (*A*) Nup160-CTF interacts with several components at the vertex of the inner Y complex. In each Y complex, Nup85, Nup43, and Seh1 constitute the short arm; Nup160 and Nup37 form the long arm; and Nup96, Sec13, Nup107, and Nup133 comprise the stem. Shown here is a surface representation for the partial inner and outer Y complexes in the CR. Protein components in the inner and outer Y complexes will be denoted with -I and -O, respectively. Components from the inner Y complex are color-coded, and the outer Y complex is colored silver. Nup107 and Nup133 are not shown for visual clarity. (*B*) Structure of full-length Nup160. Nup160 consists of a 7-bladed β-propeller followed by an extended α-helical domain. *Inset*: Helices α38-α47 of Nup160-CTF are well defined in the EM map. The EM density, shown as blue mesh, is contoured at 6 σ for depiction. (*C*) Nup160-I-CTF is sandwiched by Seh1-I and Sec13-I. (*D*) The central segments of the Nup160-I α-solenoid is pinched by Nup85-I and Nup96-I. The 96-loop from Nup160, colored magenta, spreads on the surface of Nup96.

Seh1 from the short arm and Sec13 from the stem are connected to Nup160-CTF (Fig. 2, A and C; fig. S27A). For inner Y complex, the 4CD loop (the loop between strands C and D of blade 4) of Seh1 contacts helix α40 of Nup160-CTF, whereas the β-3D strand (the D strand of blade 3) of Seh1 interacts with helix α42 of Nup160-CTF (Fig. 2C; Fig. S27A). The N-terminal α1 helices from Nup96 and Sec13 pair up to wedge into the crevice between helices α46 and α47 of inner Nup160 (Fig. 2C; Fig. S27A). These interactions may strengthen the Y complex.

A conserved surface loop between α36 and α37 of Nup160-I, referred to as the 96-loop, closely associates with the ridge of the C-terminal helices from Nup96-I (Fig. 2D, fig. S27B). Similar to that in fungi (*16, 17*), two C-terminal helices of Nup85-I contact the lateral side of the middle portion of the α-helical domain of Nup160-I (Fig. 2D, fig. S27B). The interactions described here for inner Y complex are also recapitulated in the outer Y complex (fig. S27C).

### Two Nup205 bind the inner and outer Y complexes through distinct interfaces

Our structure identifies two Nup205 molecules, defined as Nup205-I and Nup205-O, in each CR subunit (Fig. 3A; figs. S9 and S28 and S29). The two Nup205, which do not contact each other, closely associate with the inner and outer Y complexes through different interfaces (Fig. 3A, fig. S30A).

**Fig. 3.**
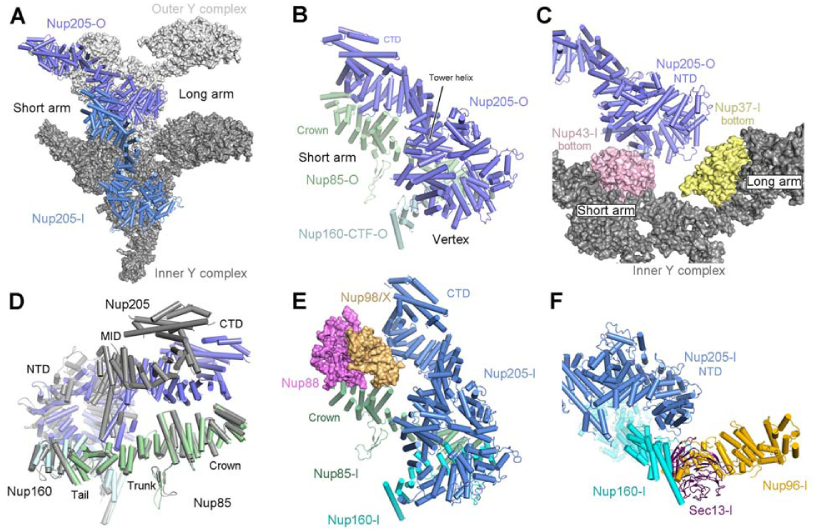
Nup205 wraps around the Y complexes. (*A*) Nup205-O associates with both the inner and outer Y complexes, whereas Nup205-I only interacts with the inner Y complex. Nup205-O interacts extensively with the short arm and the vertex of the outer Y complex as well as both arms of the inner Y complex. In contrast, Nup205-I only interacts with the short arm and the vertex of the inner Y complex. (*B*) Extensive binding interface between Nup205-O-CTD and Nup85-O. (*C*) Association between Nup205-O and the inner Y complex. (*D*) Structural elasticity of Nup205. When superimposed relative to Nup-205 associated Nup160 and Nup85, the CTD of the two Nup205 molecules exhibit pronounced conformational variations. The inner nucleoporins are color coded, and the outer counterparts are colored grey. (*E*) Nup88 and Nup98/X wedge into the space between Nup205-I-CTD and Nup85-I, disrupting their direct association. (*F*) Nup205-I binds to Sec13-I and Nup96-I from the vertex.

Nup205-O directly contacts the short arm and the vertex of outer Y complex (Fig. 3B). On one side, Tower helix α52 and six additional helices associate with the ridge of Nup85-O in the short arm. On the other side, five helices contact Nup160-CTF-O at the vertex (Fig. 3B, fig. S30B). Notably, Nup205-O also associates with both arms of the inner Y complex through interactions with the bottom faces of the β-propellers in Nup43-I and Nup37-I (Fig. 3C, fig. S30C). Through these interactions, Nup205-O connects the two Y complexes within the same CR subunit.

The interactions of Nup205-I with the middle helices of Nup85-I and Nup160-CTF-I are generally conserved as their outer counterparts (Fig. 3D-F; fig. S30D). However, there is no contact between the C-terminal helices of Nup205-I and Nup85-I, which are separated by the N-terminal propeller of Nup88 and Nup98/X from the Nup214 complex of the CF (Fig. 3E) (*13*).

These structural observations uncover previously unrecognized interfaces in the organization of the CR scaffold and identify Nup205 as a new core component of the CR. Distinct conformations of the two Nup205 demonstrates the structural elasticity that underlies their versatility for binding to different nucleoporins in the proximal and distal CR rings and the CF.

### Nup93 bridges the Y complexes and Nup205

Nup93 consists of an N-terminal extended α-helix (α5) and an ACE1 domain that are connected by flexible linkers (figs. S31). Our EM maps unveil two Nup93 ACE1 and two extended helices α5 that are inserted into the CTD of Nup205 (Fig. 4A, fig. S31).

**Fig. 4.**
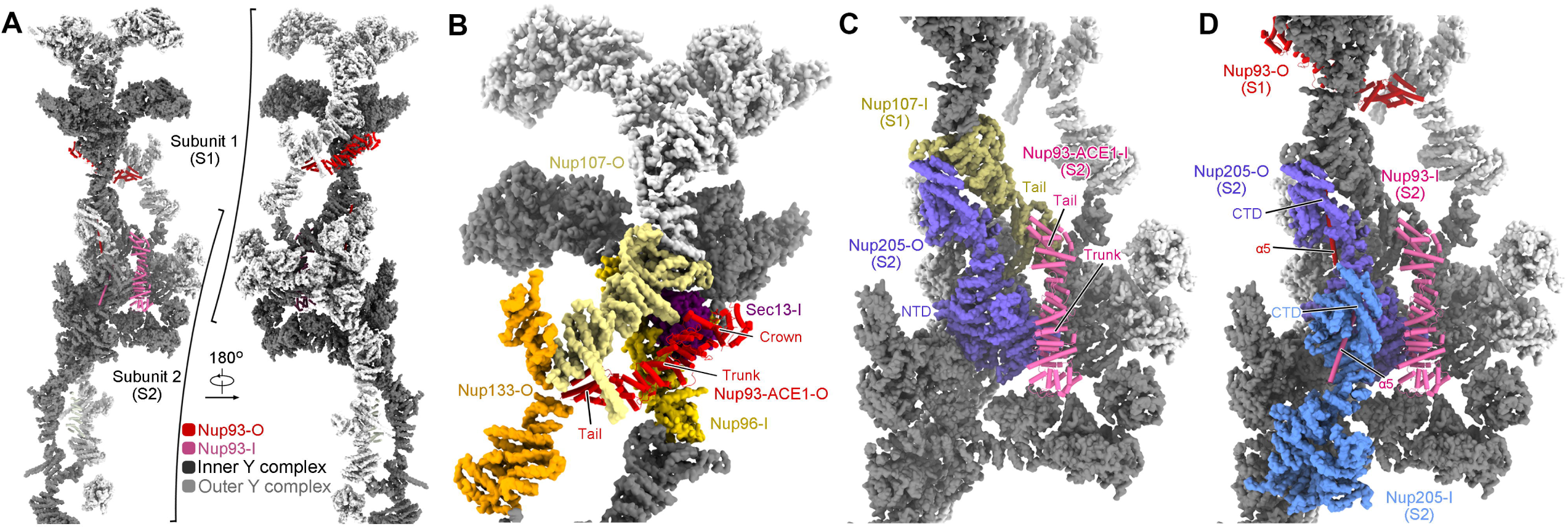
Nup93 bridges Y complexes and Nup205 molecules. (*A*) Nup93 molecules bridge adjacent Y complexes as well as Nup205 molecules. Identical components from two adjacent CR subunits, designated S1 and S2, are presented. Nup205 and the Y complexes are shown as surface. Inner and outer components are colored dark and light gray, respectively. Two copies of Nup93 are present in each CR subunit, with the one closer to the central pore designated Nup93-I (hot-pink) and the other as Nup93-O (red). For visual clarity, only Nup93-O from S1 and Nup93-I from S2 are shown. (*B*) Nup93-ACE1-O connect the stems of the inner and outer Y complexes in each CR subunit. The bridging role is achieved through direct binding to the ACE1 domains in Nup96-I and Nup107-O. These proteins form a triangular ACE1-core at the stem of the Y complexes, which is bolstered by Sec13-I and Nup133-O. Previously defined module names of the ACE1 domain, Crown, Trunk, and Tail, are labeled. (*C*) Nup93-ACE1-I bridges two adjacent subunits through direct binding to the ACE1 of Nup107-I from S1 and the NTD of Nup205-O from S2. (*D*) The α5 helices from the Nup93 molecules each insert into the axial groove of one of the two Nup205-CTDs.

Nup93-ACE1-O resides at the stem regions of both Y complexes (Fig. 4, A and B). The interfaces between Nup93-ACE1-O and surrounding nucleoporins are discernible (figs. S32 and S33). The middle portion, known as the Trunk, of Nup93-ACE1-O associates with the Trunks of Nup107-O and Nup96-I (fig. S32, A and B). The Nup93-Nup96 association appears to be strengthened by Sec13-I, which co-folds with Nup96 and uses its blades to contact the Crown of Nup93 (Fig. 4B; fig. S32B). Through these interfaces, Nup93-ACE1-O connects the two Y complexes within one CR subunit.

Nup93-ACE1-I connects Nup205-O with Nup107-I from the neighboring subunit (Fig. 4, A and C). On one end, the Trunk of Nup93-ACE1-I associates with the NTD of Nup205-O from the same subunit. On the other end, the Tail of Nup93-ACE1-I contacts the lateral side of the Tail in Nup107-I from the adjacent subunit (Fig. 4C). Consequently, Nup93-ACE1-I sews two adjacent CR subunits.

Previous biochemical characterization suggested interactions between an N-terminal region of the fungal orthologue of Nup93 and Nup205-CTD (*12*). Indeed, the predicted structures of helices α5 of Nup93 fit exceptionally well into our EM density (fig. S9C). The extended helix α5 of Nup93-ACE1-I may insert into the axial groove of the CTD of Nup205-I, and that in Nup93-ACE1-O in one subunit (S1) is likely to bind with the CTD of Nup205-O in the neighboring subunit (S2) in the same way (Fig. 4D). The association between Nup93-α5 and Nup205-CTD is almost identical to that observed in the IR and NR subunits (fig. S33) (*18-21*).

### Five Nup358 clamps in each CR subunit

Five densities each in the shape of a shrimp tail wrap about the stems of the two Y complexes (Fig. 5A). They likely belong to five copies of Nup358, a notion corroborated by our 3.0 Å cryo-EM structure of the middle α-helical region of Nup358 (residues 222-738, designated NTD2) (Fig. 5B, fig. S15). Please refer to SI for details of the assignment of the five Nup358 molecules, which will be hereafter referred to as Clamps 1-5.

**Fig. 5.**
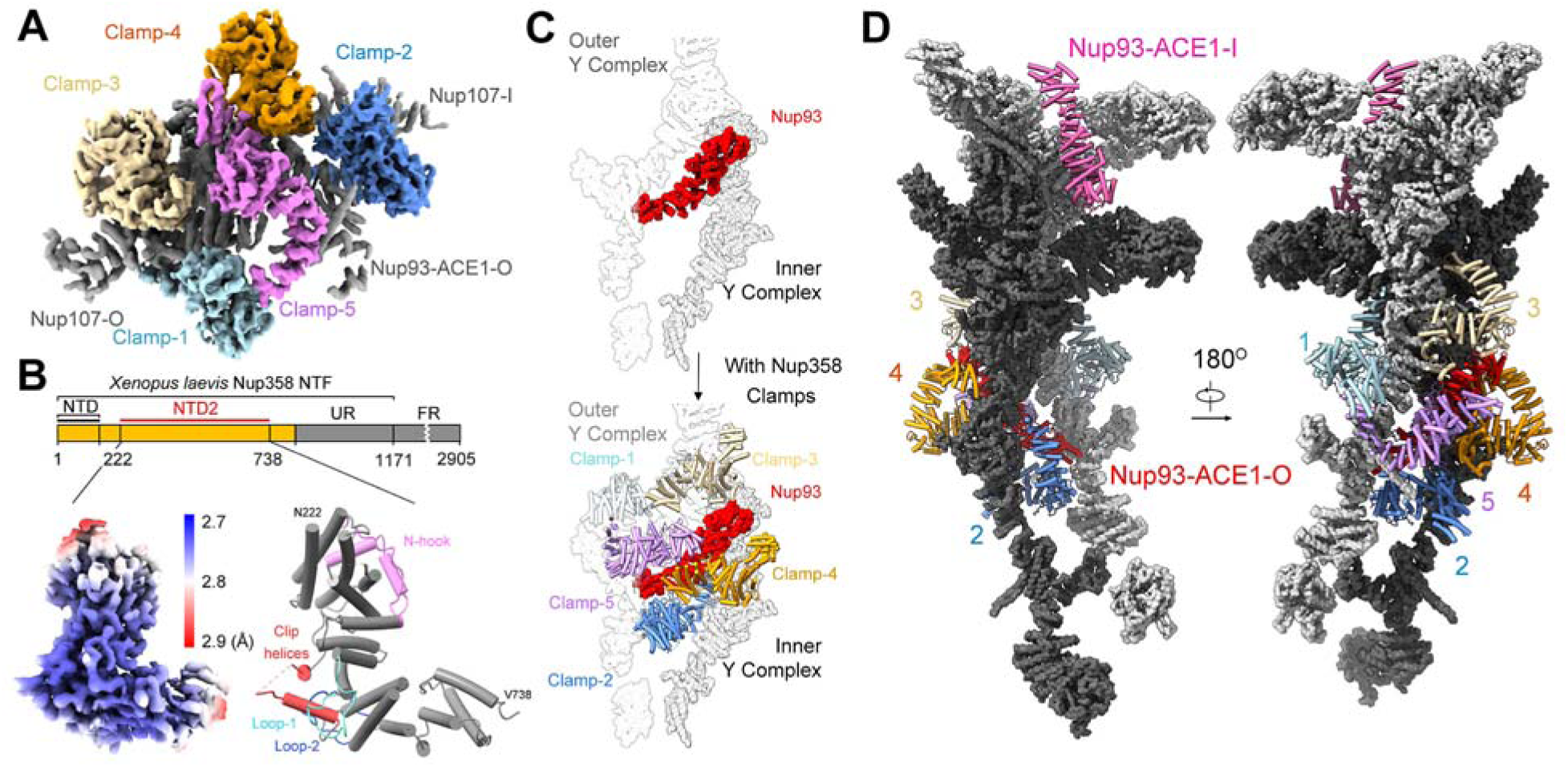
Five Nup358 clamps connect ACE1 proteins around the stems of the Y complexes. *(A)* EM density for the five Nup358 clamps within one CR subunit. The five Nup358 clamps are color coded, and Nup93 and Nup107 are colored gray. (*B*) Cryo-EM structure of Nup358-NTD2 at 3.0 Å resolution. *Left*: Resolution map of Nup358-NTD2. *Right*: Domain structure of Nup358. UR, unstructured region; FR, functional region. The terminal residues are labeled. Four surface motifs of Nup358-NTD2 (N-hook, Loop-1, Clip helices, Loop-2), which mediate the crosstalk between Nup358-NTD2 and ACE1 proteins within the CR, are color coded. (*C*) Nup93-ACE1-O and Nup358 clamps bridge the two Y complexes. Nup93-ACE1-O binds the stems of both Y complexes (upper panel). The five clamps encage the Nup93-ACE1-O (lower panel). Clamps-4 and -5 directly interact with each other to bridge the inner and outer Y complexes. (*D*) Clamp-1/3 and Clamp-2/4 pairs hold the stems of the outer and inner Y complexes, respectively. The overall CR subunit is shown in two opposite views to highlight the relative positions of the five Nup358 clamps and two Nup93 molecules. The inner and outer Y complexes are shown as light and dark grey surfaces, respectively. The Nup358 clamps and Nup93 are color coded and highlighted as cylindrical cartoons.

Clamps-1 through -4 do not interact with each other. They constitute two pairs, Clamp-1/3 and Clamp-2/4, which hold the stems of the outer and inner Y complexes, respectively (Fig. 5C,D, fig. S34). Clamp-5 connects the two pairs by binding to Clamps-1 and -4 (Fig. 5C,D, fig. S34A).

The extraordinary conformational elasticity of Nup358 allows each of these five clamps to adapt to a distinct local environment for optimal interactions with neighboring nucleoporins (fig. S34, B to D, fig. S35). Aside from the association with the Y complexes (fig. S34, C to D, fig. S35A), Clamps -2 through -5 make distinct contacts with Nup93-ACE1-O (Fig. 5C,D, fig. S35, B to E). Please refer to SI for details of the interactions. Of note, Clamp-5, whose N-terminus contacts Clamp-1 and Nup107-O (fig. S34E), places its C-terminus in close proximity to the Central domain of Nup93 (Fig. 5C,D). Together, these Nup358 molecules encage Nup93-ACE1-O within a cleft between the stems of the two Y complexes (Fig. 5D).

The interaction between Nup358 and ACE1-containing proteins mediates the crosstalk between the Y complexes (fig. S35A). Previous studies identified five candidate interfaces between Y complexes in the CR of vertebrate NPC (*10-12, 16*). Our current study uncovers additional interfaces that are clustered in two regions (figs. S35A & S36A, table S6). The first region involves Nup96-I and Nup358 Clamp-1, where the Tail of Nup96-I contacts the Crown of Nup107-O and stacks against the N-terminal α-solenoid of Clamp-1 (fig. S35A, middle and left panels). The second one involves Nup205 and the ACE1 of Nup93 molecules, whose interfaces are elaborated below.

### Interfaces between CR subunits

The 4.1-Å EM map of the core region was individually aligned to each of the eight subunits in the 22.2-Å reconstruction, generating a composite model of the CR consisting of the proximal and distal rings (Fig. 6A) (*11, 12, 16*). For simplicity, we will refer to two adjacent subunits within the CR as Subunit 1 (S1) and Subunit 2 (S2) (Fig. 6A). Nup205 plays a critical role in sewing the neighboring subunits.

**Fig. 6.**
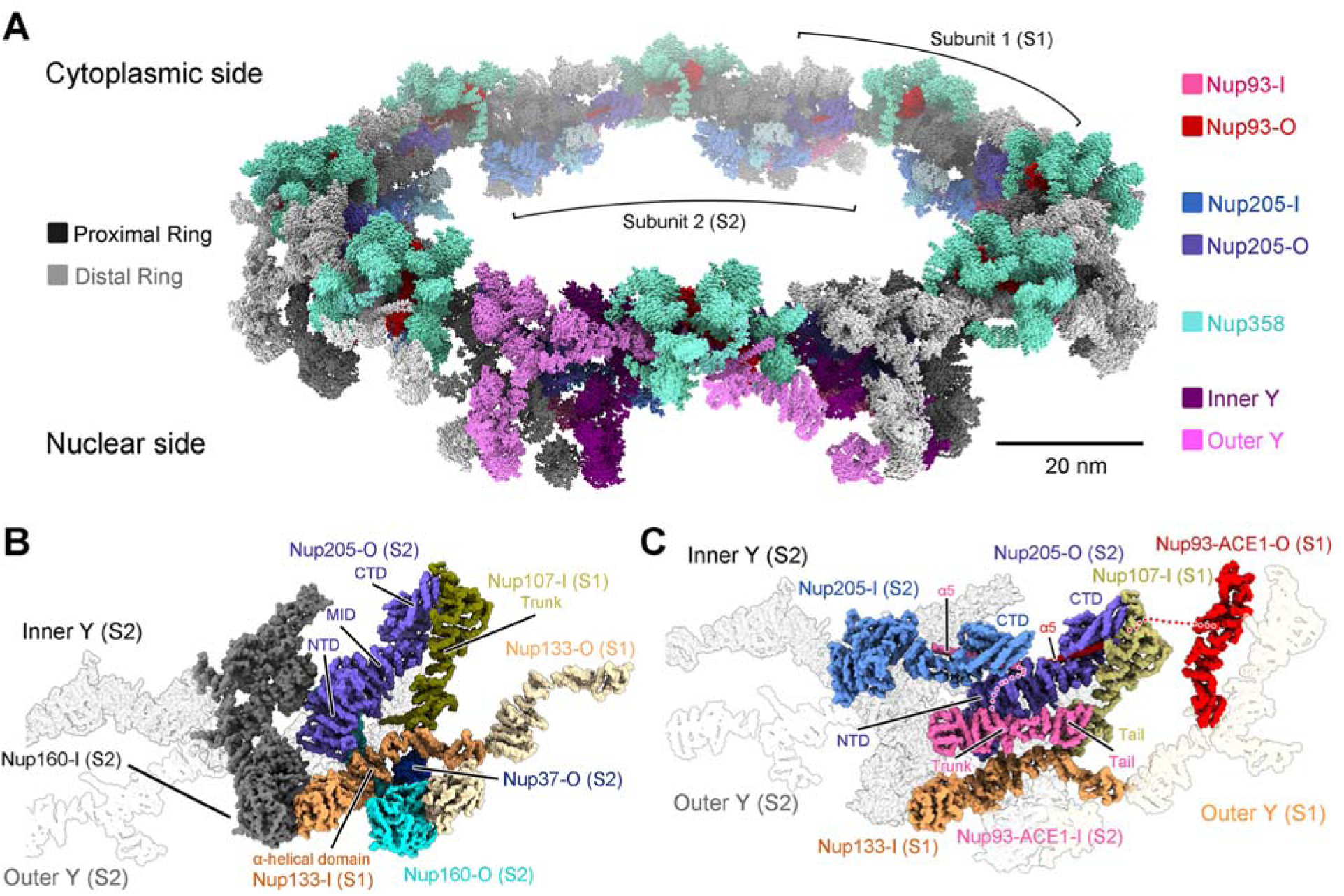
Structure of the double-layered CR scaffold. (*A*) A molecular model of the *X. laevis* CR. In addition to the two concentric Y-complex rings that each comprise eight head-to-tail assembled Y complexes, our present study identifies Nup358, Nup205, and Nup93 to be constituents of the CR scaffold. (*B*) Nup205-O is a key bridge between neighboring CR subunits. The indicated nucleoporins that map to the interface between CR subunits are color coded and labeled. (*C*) Nup93 molecules connect neighboring subunits. The ACE1 of Nup93-I (S2) connects the ACE1 of Nup107-I (S1) with the NTD of Nup205-O (S2). Nup93-α5 may sew the neighboring subunits by inserting into the axial groove of the Nup205-CTDs. Similar views are shown in panels B&C.

On one side, the NTD of Nup205-O (S2), together with Nup37-O (S2) and Nup160-O (S2), clenches the α-helical domain of Nup133-I (S1) (Fig. 6B). The Tower helix from the middle domain of Nup205-O and the Tail of Nup85 (S2) interact with the extended helix α33 from the Tail of Nup107-I (S1) (fig. S36B,C). This interface is further buttressed by the β-propeller of Nup43-O (S2) (fig. S36B). At the C-terminal end of Nup205-O (S2), its vertebrate specific Tail-C associates with the Trunk of Nup107-I (S1) (Fig. 6B, fig. S36B,C). Together, these two α-helical domains harbor the docking site for the outer copy of Nup93-α5 (Fig. 6C), which is likely linked to Nup93-ACE1-O from S1 (fig. S36C).

Nup93-ACE1-I from S2 also connects two adjacent subunits, with its Tail contacting the lateral side of Nup107-I Tail from S1 on one side, and its trunk interacting with the NTD of Nup205-O on the other terminus (Fig. 6C, fig. S36B,C). Furthermore, the extended α5 helix of Nup93-I potentially reaches into the axial groove of the CTD in Nup205-I (S2) (fig. S36C). These structural discoveries establish Nup205 and Nup93 as crucial components in ring formation.

## Discussion

For the past two decades, structural studies of the NPC at moderate resolutions have been complemented by other biochemical strategies exemplified by chemical crosslinking and mass spectrometry (*22-24*). The integrated approach has yielded a wealth of information on the overall organization of NPC (*10-12, 25, 26*). However, it is difficult to obtain accurate distance information, to determine precise stoichiometry, or to pinpoint an interface through these indirect methods. The improved EM map in this study reveals numerous previously unrecognized features (Fig. S5-S10).

The improved atomic model of the CR subunit, which validates most of the assignment in our previous study (*13*) (Fig. S37), provides a framework for mechanistic understanding of NPC assembly. NPC is subject to assembly and disassembly during a cell cycle. The Y complex has been shown to stay as an intact subcomplex throughout the open mitosis of metazoan (*27*). Our structural finding that Nup160-CTF is at the center of the multi-protein vertex of the Y complex provides a mechanistic explanation for the various conformations of the short arm in Y complexes lacking Nup160-CTF (fig. S38). Of note, missing of Nup160-CTF is related to steroid-resistant nephrotic syndrome (*28*), demonstrating its functional significance.

Our EM maps identify two additional Nup93 molecules within each CR subunit, expanding the list of ACE1 proteins within the CR. In the updated model of the CR subunit, four out of seven α-helical nucleoporins possess the ACE1 fold (*29*). Interestingly, crosstalk among the ACE1 domains of Nup93-O, Nup107-O and Nup96-I, which form an ACE1-core at the stem region of the Y complexes, is through interfaces that differ from the canonical Crown-Crown association (*30*) (Fig. 4B). Our recent analysis on the IR subunit also reveals a homo-dimerization interface of Nup93 through its Crown and Tail (*18*). These observations corroborate a key role of the multifaceted ACE1 proteins within the NPC assembly (figs. S39 and S40) (*29*). Notably, published EM maps (*9, 10*) indicate that the Nup93-ACE1-O is also present in the NR subunit from a vertebrate NPC (fig. S41). Together with four molecules of Nup93 in the IR subunit (*12, 25*), there are at least seven Nup93 molecules within each NPC spoke.

Both Nup205 and Nup188 adopt a super-helical fold that is structurally reminiscent of the karyopherin family (*20, 31*). Our EM map of the core region reveals that two Nup205, instead of Nup188, are present within each CR subunit. It is noted that the IR subunit of the *X. laevis* NPC also comprises two Nup205 molecules (*18*), whereas only one Nup205-O is found in the EM maps of the NR subunit from *X. laevis* (*9*) and *H. sapiens* (*32*) (fig. S42). These data suggest that at least five Nup205 molecules are present within each NPC spoke. The potentially differential placement of Nup205 in the CR and NR mark one point of structural asymmetry between these two rings.

Except Nup358, all other known helical nucleoporins in the CR contains a single helical solenoid. Nup358 contains multiple TPR-like repeats (*33*) that are clustered to two distinct super-helical solenoids. Relative movements of the two solenoids confer additional conformational elasticity on Nup358 (fig. S34B). Identification of the fifth copy of Nup358 expands our understanding of the role of Nup358 and highlights its complex modality in the assembly of CR. Notably, both Nup358 Clamp-5 and Nup93-ACE1-O engage in direct interfaces that connect the inner and outer Y complexes. Such interweaving may strengthen the association between the two concentric rings of the vertebrate CR (*10*). In total, our current model accounts for less than 30% of the full length Nup358. The invisible functional domains of Nup358 may project into the cytosol for events such as Ran binding (*7*).

Notwithstanding these advances, many questions remain to be addressed. An atomic model of the CR, which is required to understand detailed regulation of NPC by mechanisms like post-translational modification, is yet to be obtained. EM reconstruction of a single ring cannot provide the molecular basis for the crosstalk between the CR and the CF or the LR (*11*). Structure of the CR alone is inadequate to explain many of the NPC functions.

In summary, our final atomic model reported in this study, with more than 200 nucleoporins assigned, accounts for over 90% of the molecular mass for the ordered region of a vertebrate CR. While the majority of the CR scaffold is built up by three distinct types of α-nucleoporins, the CR scaffold is structurally solidified by 12 β-propellers that may serve as bolts and rivets to strengthen the assembly (fig. S43). Our current study markedly expands structural information on the CR.

## Acknowledgments

We thank the Cryo-EM Core and the Computing Core of Westlake University for technical support.

## Funding

The National Natural Science Foundation of China 31930059 (to Y.S.) Start-up funds from Westlake University (to Y. S.)

The China Postdoctoral Science Foundation 2021M692888 (to X. Zhan)

The National Postdoctoral Program for Innovative Talents of China BX2021268 (to X. Zhan)

## Author contributions

Y.S. conceived and supervised the project. X.Zhu., P.W., and C. Z. prepared the sample. G.H., C.Z., X.Zhan., K.L. and J.L. collected the EM data. G.H., C.Z., X.Zhan. and K. L. processed the EM data. G.H. performed the cryo-EM SPA calculation. X.Zhan. built the atomic models. Q.Z., C.Y., Q.T. and M.L. provided critical support. All authors analyzed the structure. G.H., C.Z., X.Zhan., X.Zhu. and Y.S. wrote the manuscript.

## Competing interests

Authors declare that they have no competing interests.

## Data and materials availability

The atomic coordinates of the CR subunit and NTD2 of Nup358 have been deposited in the Protein Data Bank with the accession code 7FIK and 7FIL, respectively. The EM maps for the CR core region, the Nup358 region and the NTD2 for Nup358 have been deposited in the EMDB with the accession codes EMD-31600, EMD-31601 and EMD-31602, respectively.

## Supplementary Materials

Materials and Methods

Supplementary Text

Figs. S1 to S43

Tables S1 to S6

References(*36*–*54*)

## Notes

### Competing Interest Statement

The authors have declared no competing interest.

### Summary of Updates

Author list updated

