## Supplemental Information for "Near-atomic Structure of the Cytoplasmic Ring of the *Xenopus laevis* Nuclear Pore Complex"

Science
AAAS

### Supplementary Materials for

#### Near-atomic Structure of the Cytoplasmic Ring of the *Xenopus laevis* Nuclear Pore Complex

Xuechen Zhu, Gaoxingyu Huang, Chao Zeng, Xiechao Zhan, Ke Liang, Yanyu Zhao, Pan Wang, Qifan Wang, Qiang Zhou, Qinghua Tao, Minhao Liu, Jianlin Lei, Chuangye Yan, and Yigong Shi.

**This PDF file includes:**

Materials and Methods
Supplementary Text
Figs. S1 to S43
Tables S1 to S6

#### Materials and Methods

##### Cryo-EM sample preparation

The nuclear envelope (NE) from *Xenopus laevis* (*X. laevis*) oocytes was prepared largely as previously described (*9, 13*), but with the following improvements. First, to reduce conformational heterogeneity of the NPC caused by mechanical distortion, the opened nuclear envelope was spread onto the grids as gently as possible, with minimal force applied. This practice contrasts that in previous studies (*9, 13*), in which mechanical force was applied to ensure flat placement of the NE on the grids. Second, to stabilize the NPC conformation, the NE on the grids was crosslinked for 30 minutes on ice using 0.5% glutaraldehyde in a low-salt buffer (LSB) (10 mM HEPES, pH 7.5, 1 mM KCl, 0.5 mM $MgCl_2$). Third, copper grids were replaced by gold ones in this study. Last but not least, to improve sample uniformity, each batch of samples was prepared on the same day using oocytes from the same frog. The gold EM grids (R1.2/1.3, R2/1, R2/2; Quantifoil, Jena, Germany) were blotted for 8 seconds with a blot force of 15, vitrified by plunge-freezing into liquid ethane using a Vitrobot Mark IV (Thermo Fisher Scientific) at 8 °C under 100% humidity. The quality of sample was examined using an FEI Glacios microscope (Thermo Fisher Scientific) operating at 200 kV.

##### Data acquisition of intact *X. laevis* NPC

Grids were transferred to a Titan Krios electron microscope (FEI), operating at 300 kV and equipped with a Gatan GIF Quantum energy filter (slit width 20 eV). 46,143 micrographs were recorded with the grids tilting at angles of 0, 30, 45, and 55 degrees (*13, 36*). A K3 detector (Gatan Company) was used in the super-resolution mode with a nominal magnification of 64,000x, resulting in a calibrated pixel size of 0.6935 Å for the movie files (fig. S1 and table S1). The movie images were then binned twice during motion correction, arriving at a pixel size of 1.387 Å for the final motion corrected images. The total dose followed a cosine alpha scheme where the dose is inversely proportional to the cosine value of the tilting angle. Within each stack, the exposure time for each frame and the dose rate were kept the same. Detailed statistics of data collection are reported in Table S1. All frames in each stack were first aligned and summed using MotionCor2 (*37*). Dose weighting was performed using MotionCor2 (*37*). The average defocus values were set between -1.5 and -3.0 μm and were estimated using Gctf (*38*).

##### Initial model of the cytoplasmic ring

We examined all 46,143 micrographs and manually selected 33,747 micrographs for further processing. A total of 800,825 particles were manually selected from these micrographs (fig. S2A). Initial defocus estimation was carried out as previously described (*13*) prior to all other data processing procedures.

We used the 18-Å map of the cytoplasmic ring (CR) from our previous study (*13*) as the initial reference, which was resampled to a pixel spacing of 11.096 Å and low-pass filtered to 40 Å to reduce potential model bias. We then performed one run of global search (K=1) 3D classification, with the solvent mask covering only the CR. The search had 40 iterations. The resulting star files from last several iterations were then subjected to local search 3D classifications using multiple reference seeds for guidance. We then merged all good classes and removed duplicated particles, generating a data set of 660,302 NPC particles. A final round of auto-refinement with a solvent mask on the CR resulted in a reconstruction at 22 Å resolution (fig. S2A). The resulting reconstruction was nearly identical to that in our previous study (*13*) (fig. S2B). C8 symmetry was applied throughout this stage of data processing.

**Data processing and reconstruction of the CR subunit**

We extracted the CR subunit particles based on the alignment parameters of the 22-Å CR reconstruction. We updated the orientation, shift and defocus parameters for each subunit according to a published protocol (*13*). In short, a cropping center for each subunit was defined in the map, and the Euler angles along with 2D shifts for each subunit particle were deduced from the CR particle. The re-centering procedure was repeated eight times, defining a 3D geometric relationship among eight subunits of the same CR. 5,148,474 particles of the CR subunit were extracted using a box size of 256 and a binned pixel size of 2.774 Å (fig. S3A). We then performed three rounds of CTF-refinement with geometrical restraint among subunits of the same CR and five rounds of guided multi-reference 3D classification. This practice allowed selection of 2,477,433 CR subunits, which yielded a reconstruction of the core region at 5.77 Å resolution. To fully utilize the dataset, these subunits were projected back to the original CR particles. The defocus values of all subunit particles within the same CR were pooled together to calculate a corrected average defocus value for the center of mass of each CR particle. All subunits of this CR were then re-extracted using the updated defocus value deduced from the corrected average defocus of this CR particle. Through this procedure, 4,528,642 subunit particles from 581,882 CR particles were extracted, resulting in a reconstruction at 5.6 Å after auto-refinement (fig. S3A).

This data set was then used for re-extraction with a box size of 400 and pixel size of 1.387 Å (fig. S3B). Following our published protocol (*13*), data processing beyond this point was performed individually for four datasets defined by the tilting angles: tilt0, tilt30, tilt45, tilt55. This is due to our observation that reconstruction at relatively high resolution is easily biased towards low-tilt datasets due to their relatively high signal-to-noise ratio (SNR). In fact, the high-tilt and low-tilt datasets exhibit quite different motion statistics (fig. S1). The four datasets were then separately reconstructed to obtain an estimation of SNR for their respective tilting angles. After polishing and CTF refinements, the four datasets were pooled for 3D auto-refinements and 3D classifications.

A second strategy to enhance the quality of the reconstruction is to improve the results of particle polishing. In particle polishing, initial movement tracks are obtained either through global frame alignment or polynomial motion models determined from local motion trajectories (*39*). Magnitude of the sample drifts in the tilted datasets appears much greater than that in the un-tilted ones. This problem, along with increased sample thickness at high-tilt angles, prevents the local motion model from accommodating sample deformation. An initial motion model estimation using MotionCor2 resulted in 18,619 failed attempts to explain local sample deformation in 38,583 movie stacks using the polynomial model. In particular, sample deformation was successfully explained for only 6,825 movie stacks out of a total of 22,132 for the tilt45 and tilt55 datasets. Errors introduced by the polynomial motion model undermine the accuracy of the initial movement tracks for particle polishing, thus affecting the final outcome. To address this issue, we took advantage of the large size of a single NPC particle and attempted to estimate only the local motion around each NPC particle instead of dividing the micrograph into rectangular patches and estimating motion for each patch. A major rationale for this practice is that empty patches, which often occur, may engender misalignment, thereby compromising the accuracy of the polynomial motion model. To improve alignment, we used a polynomial motion model similar to those implemented in MotionCor2 (*37*), RELION (*40*), or WARP (*41*) to regularize movement tracks of particles that belong to the same micrograph. This polynomial model was iteratively refined until convergence. A final round of patch-based alignment using only information around each NPC particle was performed to account for any residual movement of each particle. Finally, any micrograph that failed to converge or had residual motion of more than 15 pixels in any direction was removed.

Using the above strategy, we were able to rescue 13,758 micrographs from 18,619 micrographs that failed to obtain reliable local motion models. We separately reconstructed the four tilt datasets and performed one round of CTF-refinement. The results were directly subjected to auto-refinement, resulting in a reconstruction of the core region at 4.5 Å resolution (fig. S3B). The combined dataset was then re-divided into four subsets according to their tilting angles and subjected to particle polishing. The resulting particles were again pooled together and subjected to auto-refinement and multi-reference 3D-classifications. The above cycle was repeated several times to arrive at a final average resolution of 4.1 Å from 1,279,381 particles (fig. S4A and table S1).

For the Nup358 region, the same set of particles were enlarged during the particle polishing procedure, using the –window and –scale option from relion_motion_refine. The particles with enlarged box size were then subjected to local refinement in cryoSparc (*42*) to reach an average resolution of 4.7 Å (fig. S4B and table S1). The EM maps in the core region display distinct features for secondary structural elements and some of the bulky amino acid side chains (figs. S5 to S10). These features facilitated sequence assignment of the nucleoporins and identification of protein-protein interfaces.

**Expression and purification of Nup358-NTF**

The cDNA encoding the N-terminal 1,300 amino acids of *X. laevis* Nup358 (Uniprot: A0A1L8HGL2) was synthesized with codon optimization (QinglanBiotech). The N-terminal fragment (residues 1-1171, referred to as Nup358-NTF) with an N-terminal Flag tag was cloned into the pCAG vector. This construct was verified by DNA sequencing. HEK293F cells (Invitrogen) were cultured in SMM 293T-II medium (SinoBiological Inc.) supplemented with 5% $CO_2$ in a Multitron-Pro shaker at 130 rpm at 37 °C. The cells were transfected at a density of ~$2\times10^6$ cells per mL. SMS 293-SUPI (SinoBiological Inc.) was supplemented into the culture 24 hours after the transfection. The cells were then cultured for additional 24-36 hours.
The transfected HEK293F cells were harvested by centrifugation at 3,800 g and resuspended in a lysis buffer containing 25 mM Tris-HCl (pH 8.0), 150 mM NaCl, and a cocktail of protease inhibitors (VWR). The final concentrations of the inhibitors were 2 mM for phenylmethylsulfonyl fluoride (PMSF), 5.2 µg/ml for aprotinin, 2.8 µg/ml for pepstatin and 10 µg/ml for leupeptin. The cells were lysed by ultrasonication (Vibra-Cell, SONICS). After centrifugation at 30,000 g for one hour, the supernatant was loaded into an anti-Flag M2 affinity gel (Sigma) column and eluted with the lysis buffer supplemented with 200 µg/ml Flag peptide. The eluted fraction was concentrated using a 50-kDa cut-off Centricon (Millipore) and applied to size exclusion chromatography (Superose-6, GE Healthcare). The peak fractions were analyzed by SDS-PAGE (fig. S11A).

**Cryo-EM data acquisition for Nup358-NTF**

An aliquot of 4 µL freshly purified Nup358-NTF at a concentration of about 2 mg/ml was placed on glow-discharged holey carbon grids (Quantifoil Au 300 mesh, R1.2/1.3). Grids were blotted for 3.5 seconds and plunge-frozen in liquid ethane cooled by liquid nitrogen using Vitrobot Mark IV (Thermo Fisher) at 8 °C under 100% humidity. The grids were transferred to a Titan Krios electron microscope (Thermo Fisher) operating at 300 kV and equipped with a GIF Quantum energy filter (Gatan Company). Using the same setup as that for intact *X. laevis* NPC data acquisition, a total of 17,147 movie stacks were recorded. WARP was used for all subsequent preprocessing including CTF estimation and Motion correction (*41*) (Figs. S11B and S12).

**Image processing for Nup358-NTF**

Out of a subset of 1,176 micrographs, 627,188 particles were first auto-picked using Topaz (*43*) with its default model on micrographs pre-binned by eight times. Particles were extracted using a box size of 100 and a pixel size of 2.774 Å. These particles were then subjected to four rounds of 2D classifications in cryoSparc (*42*), yielding 201,480 particles. The good particles were then converted to RELION compatible star files using the csparc2star.py program from the pyem suite (*44*). These particles were used to train a deep learning model in Topaz. Relying on this model, we picked 2,418,968 particles from 17,147 micrographs using a box size of 100 and a binned pixel size of 2.774 Å. These particles were directly subjected to two additional rounds of 2D classifications in RELION, yielding a data set containing 2,047,018 particles (fig. S12). These particles were then re-extracted using a box size of 200 and a pixel size of 1.387 Å and imported into cryoSparc. Eight jobs of initial model generation were executed and the best initial model was selected based on visual inspection.

We then used csparc2star.py to convert the 213,534 particles assigned to this class into RELION compatible star files and used the orientational parameters contained within to generate a RELION compatible reconstruction. Using this initial model, one round of random phase 3D classification with the random_mask and solvent_mask options turned on was performed on the whole dataset. Similar to a published procedure (*45*), all other references except for class1 were phase randomized to generate a "bad" reference with the user specified phase randomization upper and lower limits. To further distinguish protein from solvent, the EM density outside the user supplied random_mask was kept unchanged in the "bad" references; the EM density inside the random_mask was subjected to the phase randomization procedure. Conversely, the EM density inside the random_mask was kept unchanged in the "good" reference and that outside was subjected to phase randomization.

Using this approach, we were able to perform image alignment for the relatively small Nup358-NTF protein for up to 40 iterations. After the global search random phase classification procedure, data stars from the last several iterations from the global search run were subjected to local search multi-reference 3D classifications. The good classes were merged and duplicated particles were removed. Finally, the remaining 744,385 particles were imported into cryoSparc for a final round of local refinement to yield reconstructions at an average resolution of 3.0 Å for two separate regions that are flexible relative to each other (figs. S12, D and E, and S12, and table S1). The final EM reconstruction exhibits clear features for sequence assignment (fig. S13).

**Structural modeling of the CR subunit**

The 4.7-Å map of the Nup358 region contains five similar, clamp-shaped density patches. To facilitate modeling of these five clamps, we first generated the atomic model and assigned sequence register for Nup358-NTD2 (residues 222-738) on the basis of the 3.0-Å EM map of Nup358-NTF (fig. S14). The atomic model of Nup358-NTD2 was refined against the 3.0-Å map using Phenix real space refine (*46*) with secondary structure restraints.

The final atomic model of Nup358-NTD2 can be placed into each of these five density patches with little adjustment (fig. S15). Based on the X-ray structure of human Nup358-NTD1 (*33*), we generated a homology model for NTD1 (residues 1-145) of Nup358 from *X. laevis*. Nup358-NTD1 and Nup358-NTD2 were separately docked into each of the five clamp-shaped density areas, respectively. In each of these five areas, this practice leaves a similar density patch unfilled between NTD1 and NTD2. This unfilled density patch was assigned to the sequences between NTD1 and NTD2 (residues 146-221), which were predicted to be α-helical. Together, we generated an atomic model for residues 1-738 of Nup358, which was individually fit into each of the five clamp-shaped density areas (fig. S15, B and C). This observation suggests the presence of a fifth Nup358 molecule in addition to four speculated Nup358 molecules in the Nup358 region (fig. S15A) (*32*).

Apart from the newly resolved structure of Nup358-NTD2, modeling for the CR subunit were largely based on The atomic coordinates of *X. laevis* CR from our previous study (PDB: 6LK8) (*13*), which were fitted into the new reconstruction of the CR subunit using Chimera (*47*). For the core region, the secondary structural elements were manually checked and adjusted based on the much improved EM density map using Coot (*48*). The reconstruction of the core region at

4.1-Å resolution allowed us to identify a large number of bulky residues in components of both the inner and outer Y complexes and in Nup205. We *de novo* modeled 10 α-helices in the newly assigned C-terminal fragment of Nup160 (Nup160-CTF), which was manually adjusted based on the sequence conservation and secondary structural elements extracted from AlphaFold model (*14*) (fig. S16). For the less well resolved N-terminal regions of Nup160-O, modeling was carried out through docking of the coordinates of the N-terminal regions of Nup160-I into the EM density of core region lowpass filtered to 10 Å, in which the outer long arm is clearly discernible. In addition, the β-propeller of Nup88, the autoproteolytic domain (APD) of Nup98, and the C-terminal domain (CTD) of Nup155 were also modeled using the predicted structures, respectively. The current EM density does not allow for reliable docking for Nup98. We will refer to this model as Nup98/X throughout the manuscript.

For modeling of Nup205 molecules, we generated the secondary structural elements of Nup205 based on structural alignment of its functional orthologues and the crystal structure of Nup205 orthologues in fungi, this process was assisted by the AlphaFold model (fig. S17) (*21*). Relying on structural features of the IR subunit in various organisms (*18, 20, 21*), previous biochemical characterizations of the Nup205 orthologue in fungi (*16*), and the EM density (fig. S9C), we also modeled two Nup93 α5 helices inserted into the CTD of Nup205 molecules. Together, our model covers the entire α-helical region of Nup205, including a CTD and vertebrate specific TAIL-C that harbor the docking site of Nup93 α5.

In the center of the Nup358 region, the Bridge domain of unknown identity connects the Nup358 clamps and associates with other surrounding nucleoporins (*13*). In both the NR from frog and human NPC, a rod of EM density with nearly identical shape resides in the same place as the Bridge domain within the two concentric Y complex rings. Previous analysis pointed to TPR, a nuclear basket specific nucleoporin as a candidate that may occupy this location (*10*). The 4.7-Å EM map of the Nup358 region allowed generation of a poly-Ala model for the Bridge domain. The poly-Ala model was used to search the PDB using the DALI server (*49*). This search led to identification of a number of potential candidates, of which the yeast NPC component Nic96 tops the list (table S5), suggesting the Bridge domain to be Nup93, the *X. laevis* ortholog of Nic96.

Near completion of this manuscript, DeepMind released a database of 20,000 predicted structures of the human proteome, which includes that of human Nup93 (*50*). To our satisfaction, the poly-Ala model of the Bridge domain and the predicted model of human Nup93 display an

RMSD of 2.43 Å over 416 aligned Cα atoms (fig. S31A). In fact, the predicted structure of human Nup93 can be directly docked into the EM density map for the Bridge domain without adjustment (fig. S31B). The nearly perfect fitting of human Nup93 at the secondary structure level strongly supports the identification of the Bridge domain to be Nup93. This conclusion is further supported by mass spectrometric analysis of the crosslinked *X. laevis* NE, in which Nup93 was found to be mainly crosslinked to Nup358 among the nucleoporins (fig. S30C).

Using AlphaFold (*14*), we generated a predicted atomic model for *X. laevis* Nup93, which fits the EM density map exceedingly well (fig. S30D). The surface loop region of the predicted model for *X. laevis* Nup93 was manually adjusted into the EM density. This final atomic model of *X. laevis* Nup93 is almost identical to our initial poly-Ala model of the Bridge domain but contains more features. The featured atomic model of *X. laevis* Nup93 (fig. S30E) was used for all subsequent structural analysis. The structure of *X. laevis* Nup93 α-solenoid (residues 180-820), with a fold-back conformation characteristic of ACE1 (*29*), consists of a Trunk module (α6-α9&α18-28), a Crown module (α10-α17) and a Tail module (α29-α37) (Fig. 5*B*; Fig. S30E). In addition, in the N-terminus of the Nup93 α-solenoid, an extended α-helix (α5) inserted in the CTD of Nup205 was also modeled (figs. S9C and S30E). The corresponding region in yeast Nic96 was reported to bind the CTD of Nup192 (fungal homolog of Nup205) (*12*). Secondary structural elements were assigned based on sequence alignment of functional orthologues of Nup93 and the AlphaFold model (fig. S31). The EM maps also allow identification of protein-protein interfaces mediated by Nup93 (figs. S9C, S32 and S33).

With one Nup93-ACE1 assigned to the Bridge domain and two Nup93 α5 helices identified to be inserted into the CTD of Nup205 molecules, we sought whether a second copy of Nup93-ACE1 is present within the CR subunit. The current molecular model left a chunk of unassigned density spanning the Nup107-I from one subunit and Nup205-O from the adjacent subunit that adopts a similar shape as those of the Bridge domain. Local-refinements focused on this chunk of density revealed that a second copy of Nup93-ACE1 is indeed present in the CR subunit. Direct docking of the model of Nup93-ACE1-O into this region resulted in a nearly-perfect match (fig. S31F).

Modeling and structural analyses of other nucleoporins were facilitated by sequence alignment (*51, 52*) of functional orthologues of the structurally defined CR components (Fig. 1D, figs. S16-S25) and AlphaFold predicted models. Overall, the final atomic model of the CR subunit includes the inner and outer Y complexes, two Nup205 molecules, five Nup358 Clamps, two Nup93 molecules, and one copy of Nup88, Nup98/X and Nup155 each. This atomic model includes 19,037 amino acids in 749 α-helices and 380 β-strands (tables S2 and S3). The final model was refined using Phenix (*46*) with secondary structure restraints and validated through examination of the Molprobity scores and statistics of the Ramachandran plots (*53*) (table S1). Together with the EM density map, this model allows structural analysis of Nup205 (Fig. 3, Fig. figs. S27 to S29), Nup93 (Fig. 4, figs. S30 to S33), and Nup358 clamps (Fig. 5, figs. S34 and S35). In addition, the final model provides basis for structural comparison with previous results (fig. S37), among Y complexes from various species (fig. S38), and between individual components of the CR (figs. S39, S40, and table S4),

The figures for structures and maps were generated using Pymol, Chimera or ChimeraX (*35, 47*). Sequence alignments were done in Clustal Omega (*51*) and ESPript 3.0 (*52*).

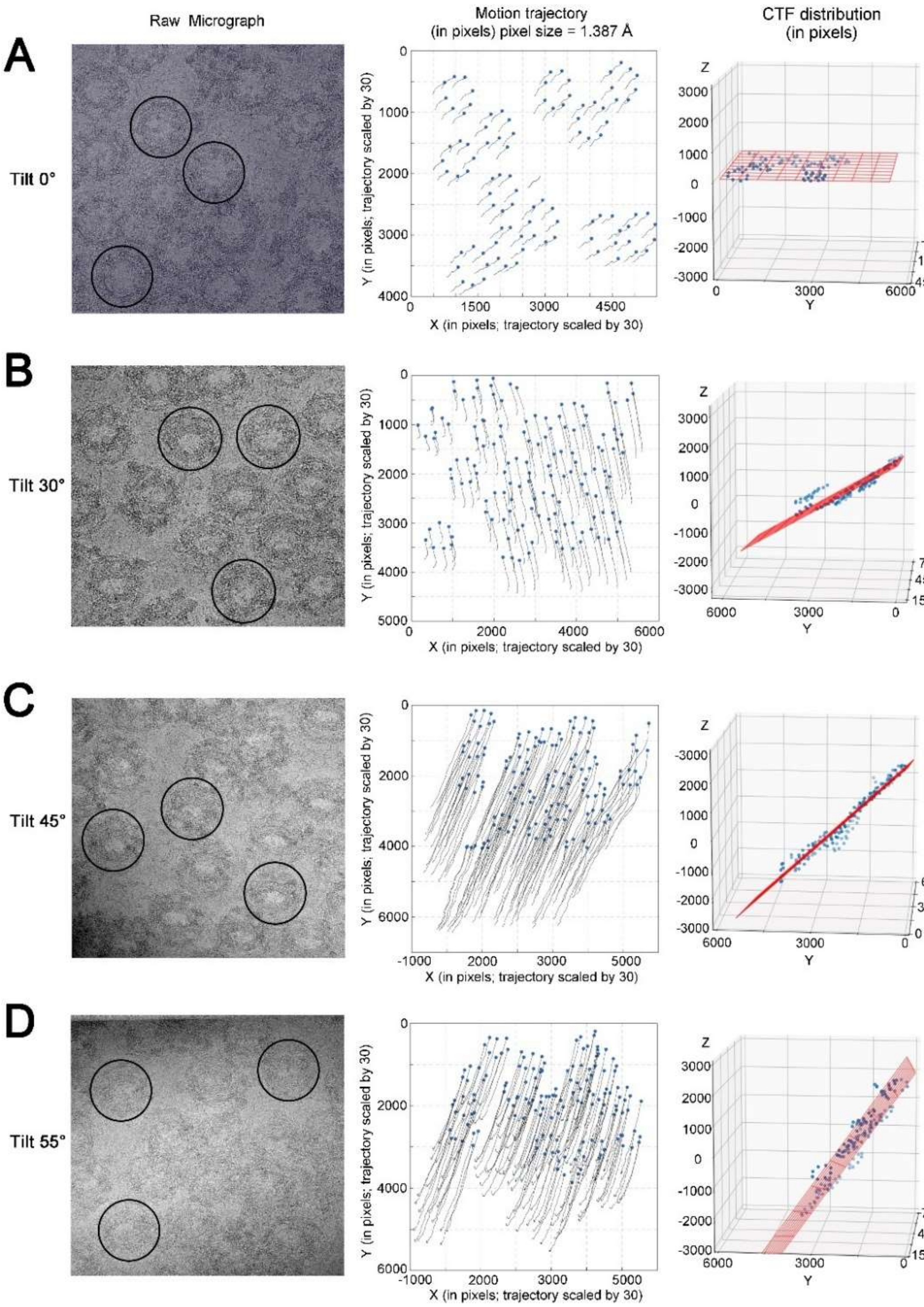
Raw Micrograph
Motion trajectory
(in pixels) pixel size = 1.387 Å
CTF distribution
(in pixels)
A
Tilt 0°
B
Tilt 30°
C
Tilt 45°
D
Tilt 55°
Y (in pixels; trajectory scaled by 30)
X (in pixels; trajectory scaled by 30)
Z
X
Y

**Fig. S1.**

**Analysis of the quality of the cryo-EM data for the NPC from *X. laevis* oocytes.** (**A-D**) Representative micrographs from the datasets for tilt-0$^{o}$ (***A***), tilt-30$^{o}$ (***B***), tilt-45$^{o}$ (***C***), and tilt-55$^{o}$ (***D***) and their quality assessment. A summed micrograph with dose weighting applied, the motion trajectory, and the CTF distribution of the CR subunit core region are shown on the left, middle, and right, respectively, in each panel. Compared to the tilt-0$^{o}$ dataset, micrographs of the tilt-30$^{o}$, tilt-45$^{o}$ and tilt-55$^{o}$ datasets display increasingly large degrees of motion and wide distribution of per-particle defocus values.

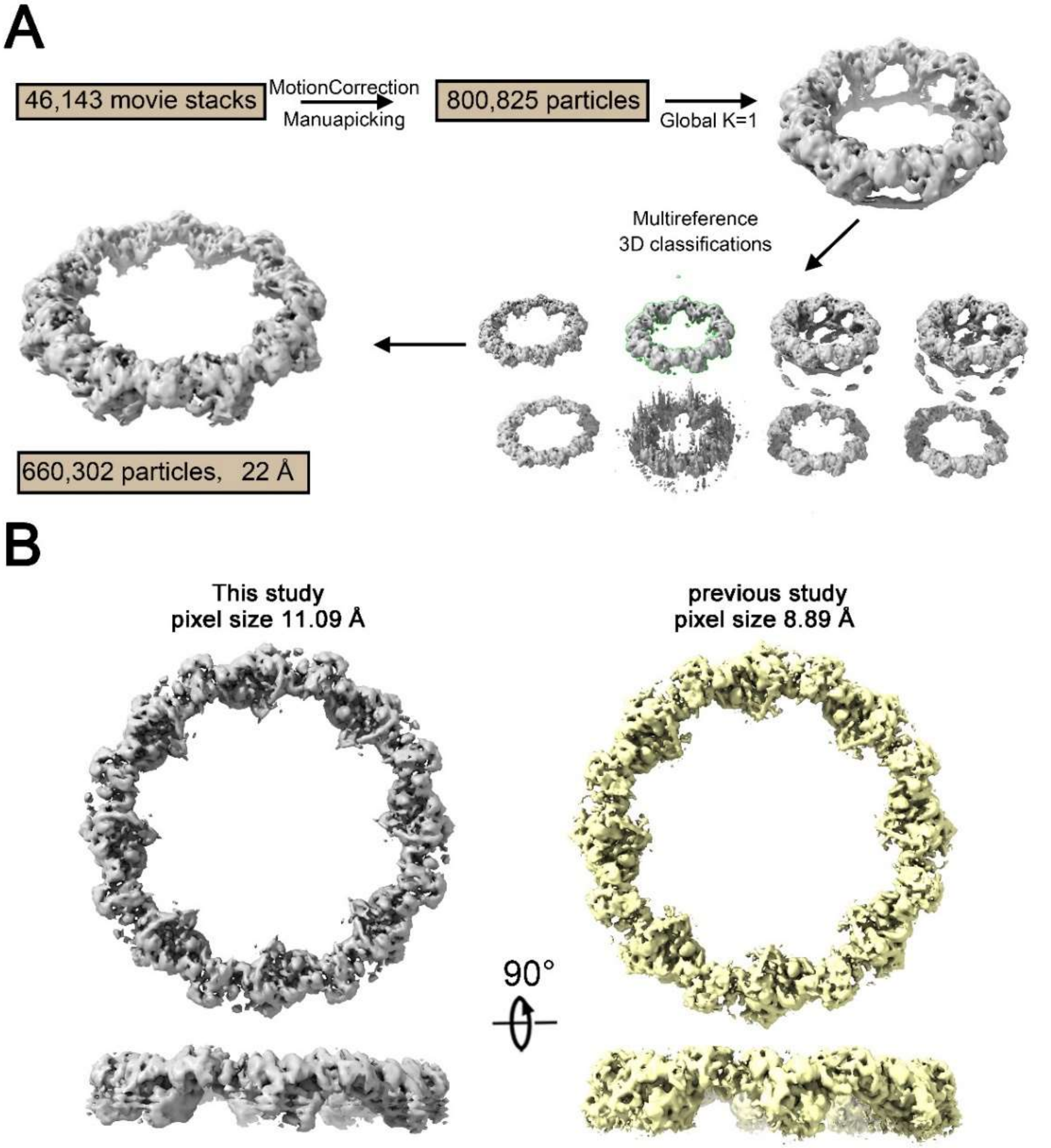


**Fig. S2.**

**Preliminary processing of the cryo-EM data for the reconstruction of the intact CR from *X. laevis* NPC.** (**A**) Flowchart of data processing towards generation of the reconstruction for the complete CR of the *X. laevis* NPC. For details, please refer to the "Initial model of the cytoplasmic ring" section in Material and Methods. (**B**) Structural comparison of the CR from the crosslinked *X. laevis* NPC (left panel) with that without crosslinking in our previous study (*13*) (right panel). Two representative views are shown. The overall structural features are nearly identical to each other.

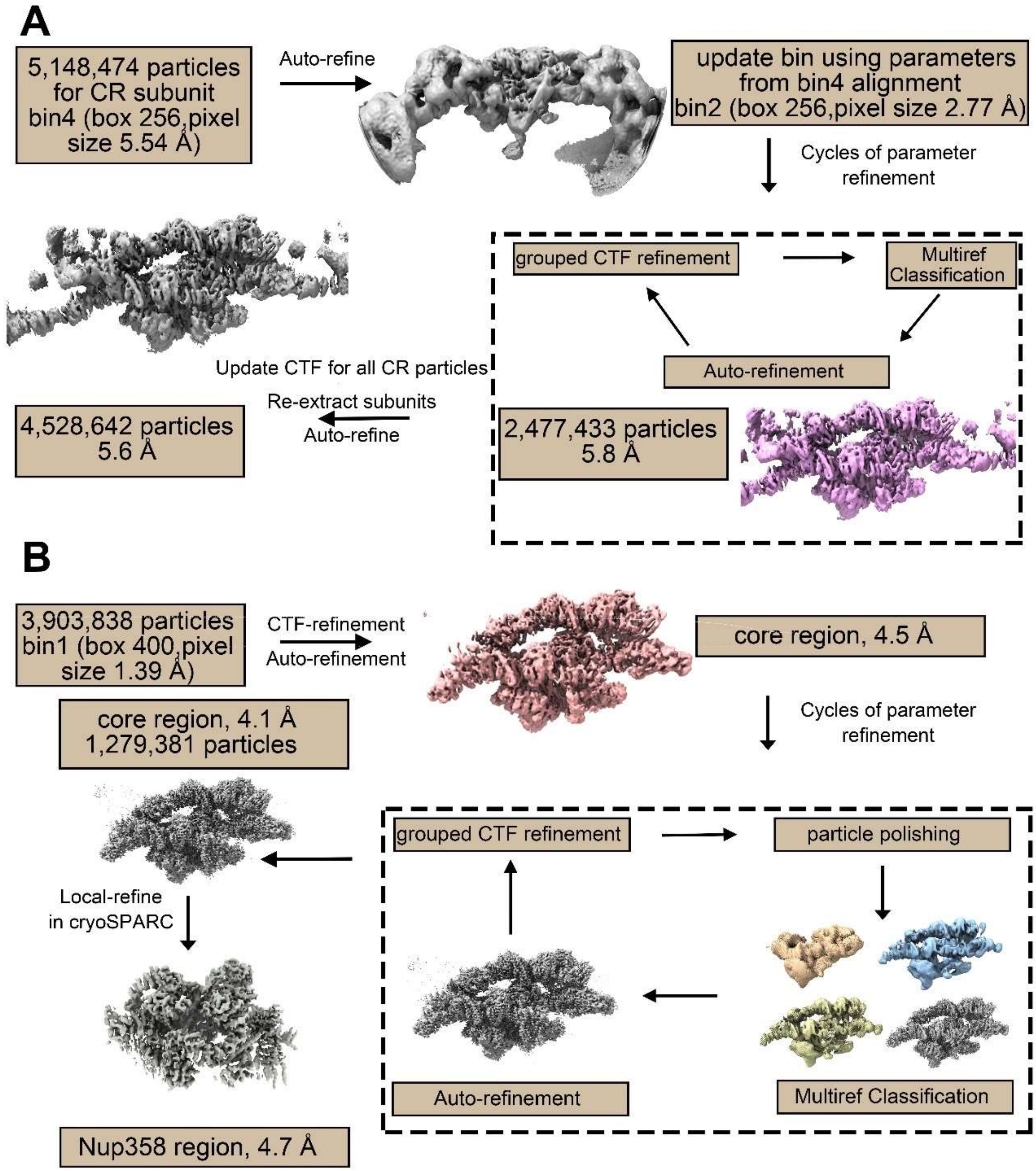


**Fig. S3.**

**Cryo-EM data processing for the reconstruction of the CR subunit from *X. laevis* NPC.** (**A**) Flowchart of bin2-level data processing for the reconstruction of the core region of the CR subunit. The core region was defined based on its central position and relative structural stability in the CR subunit. The remaining visible and continuous chunk was defined as the Nup359 region. Please refer to Fig. 1B for the approximate demarcation of the two regions. This practice resulted in a reconstruction of the core region at an average resolution of 5.6 Å. (**B**) Flowchart of bin1-level data processing for the reconstruction of the core region and Nup358 region of the CR subunit. This practice led to a reconstruction of the core region at an average resolution of 4.1 Å and the reconstruction of the Nup358 region of the CR subunit at an average resolution of 4.7 Å. For details, please refer to the "Data processing and reconstruction of the CR subunit" section in Material and Methods.

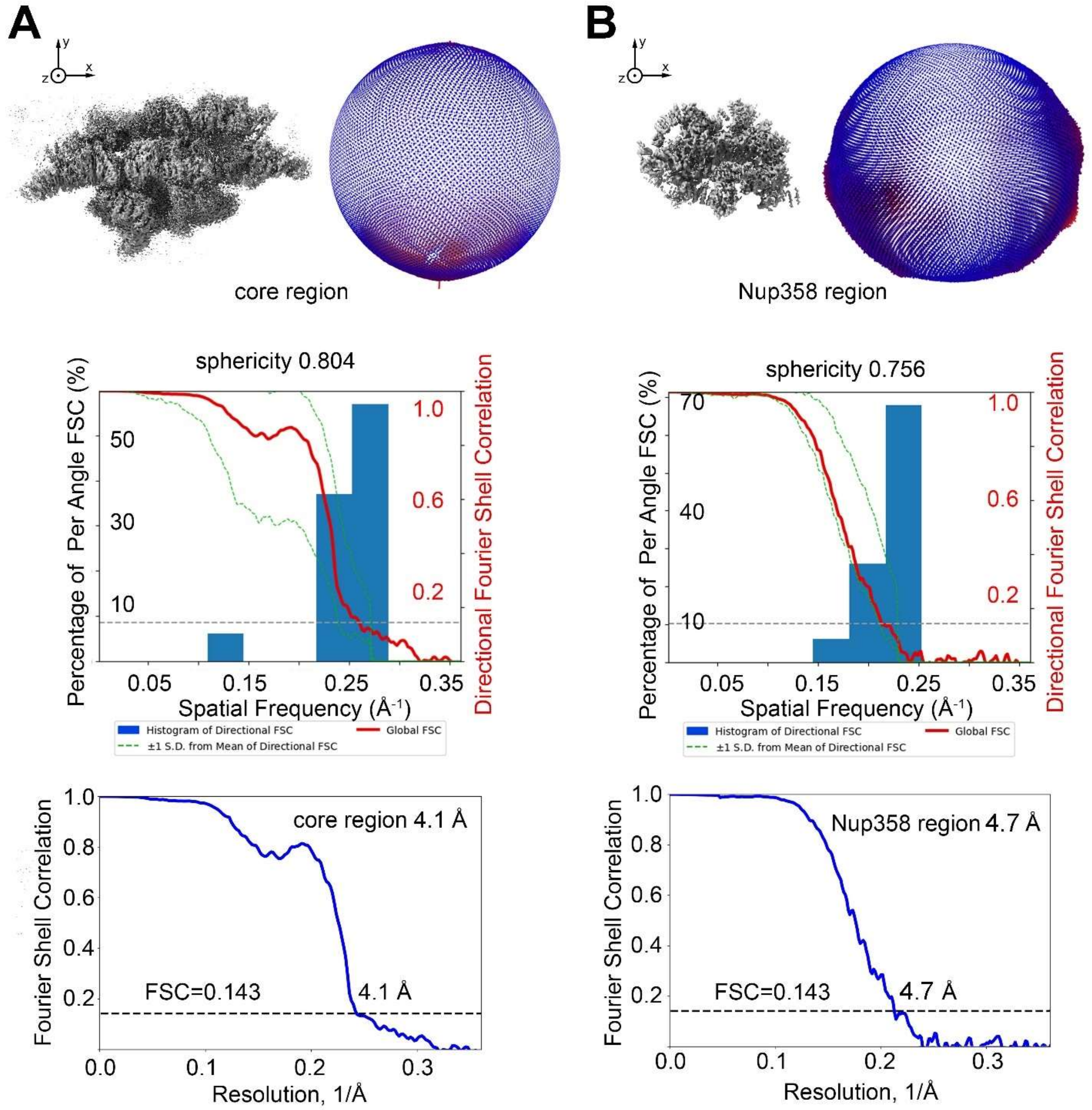


**Fig. S4.**

**Cryo-EM reconstruction of the core region (A) and the Nup358 region (B).** Angular distribution of the particles used for the final reconstruction, the directional FSC of the core region, and the final FSC for the core region cryo-EM reconstruction are shown in the top, middle and bottom panels, respectively. All directional FSC curves were prepared using the following website: https://3dfsc.salk.edu (*36*).

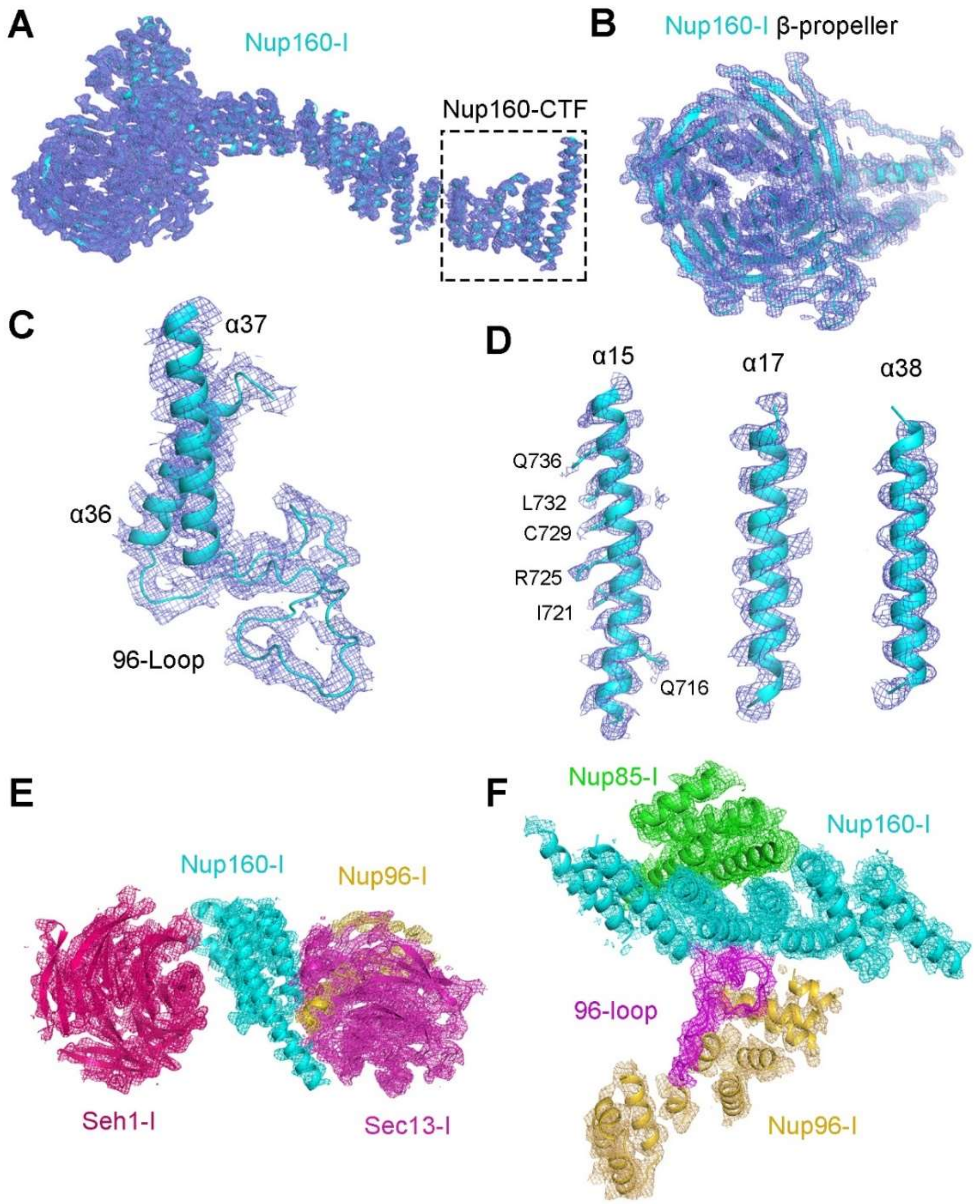


**Fig. S5.**

**Representative EM maps for Nup160.** (**A**) Overall EM map of Nup160 from the inner Y complex (Nup160-I). (**B**) EM map for the N-terminal β-propeller domain of Nup160-I. The quality of EM map allows identification of individual β-strands in the seven blades. (**C**) EM map for a surface loop of Nup160 that interacts with Nup96 (known as the 96-loop). (**D**) EM maps for three representative α-helices in Nup160-I. The helical features of the α-helices are clearly visible and some of the side chains are well defined by the density. (**E**) EM map around Nup160-I-CTF. Nup160-I-CTF is sandwiched by Seh1-I and Sec13-I. This panel corresponds to Fig. 2C. (**F**) EM map around the central portion of the α-helical domain of Nup160-I, which interacts with Nup85-I and Nup96-I. This panel corresponds to Fig. 2D. All EM maps in this figure were prepared using the reconstruction of the core region with a contour level between 5 – 10 σ in PyMol. The same applies to all density figures hereafter if not otherwise indicated.

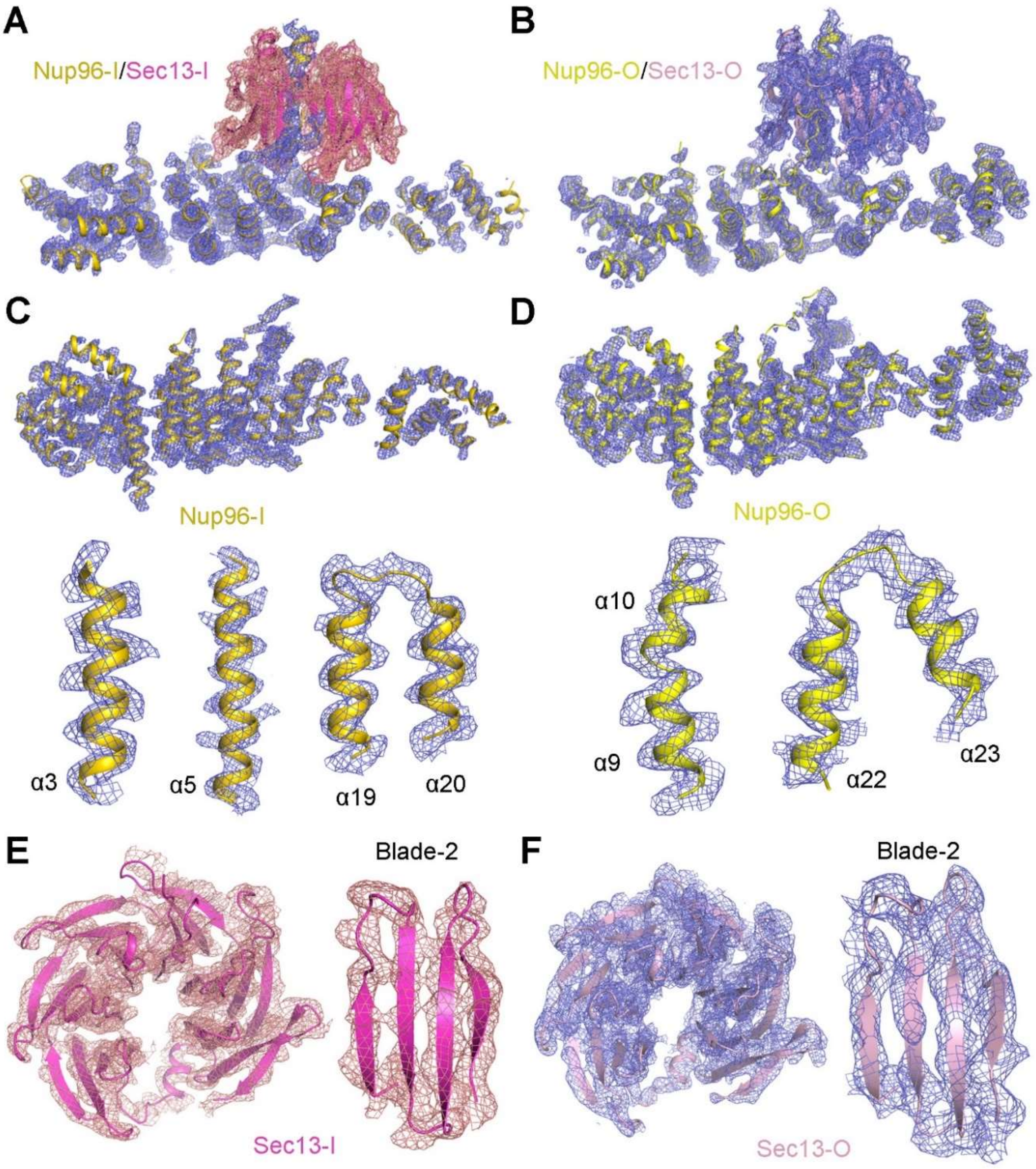


**Fig. S6.**

**Representative EM maps for Nup96 and Sec13.** (**A,B**) Overall EM maps of Nup96-I/O and Sec13-I/O. (**C,D**) EM map for Nup96-I/O. The overall views in the upper row are related to the corresponding ones in (A,B) by a 45-degree rotation around a horizontal axis. The EM maps for representative α-helices are shown in the lower row. (**E**) EM maps for the β-propeller domain of Sec13-I. The intact β-propeller has seven blades, six of which come from Sec13. Blade-2 is shown on the right. (**F**) EM maps of the β-propeller domain of Sec13-O. Blade-2 is shown on the right.

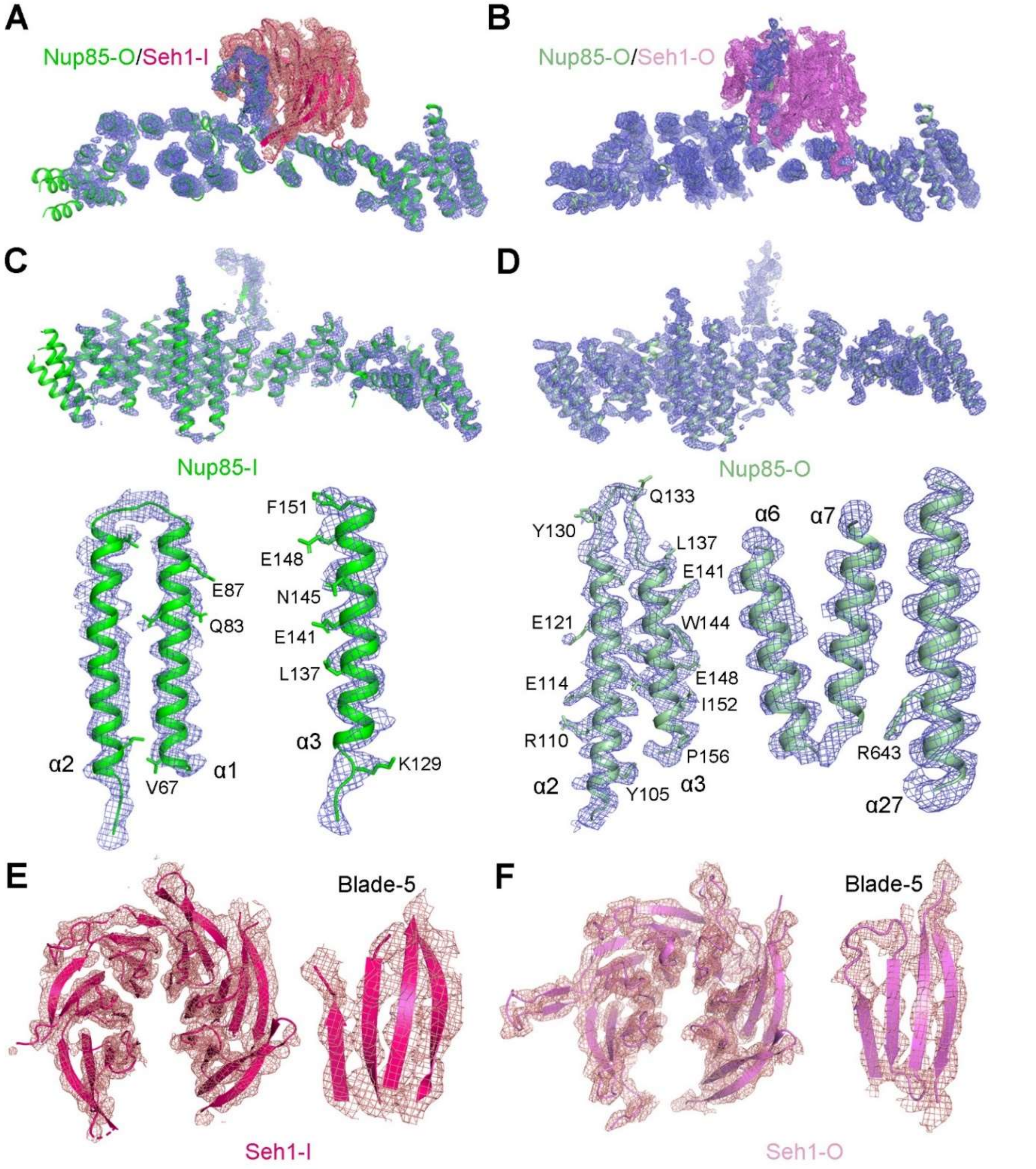


**Fig. S7.**

**Representative EM maps for Nup85 and Seh1.** (**A,B**) Overall EM maps for Nup85-I/O and Seh1-I/O. (**C,D**) EM maps for Nup85-I/O. The overall views in the upper rows are related to the corresponding ones in (A,B) by a 45-degree rotation around a horizontal axis. EM densities for representative α-helices are shown in the lower row. (**E,F**) EM map of the β-propeller domain of Seh1-I/O. Six blades of the β-propeller come from Seh1. The EM map for Blade-5 is shown on the right in each panel.

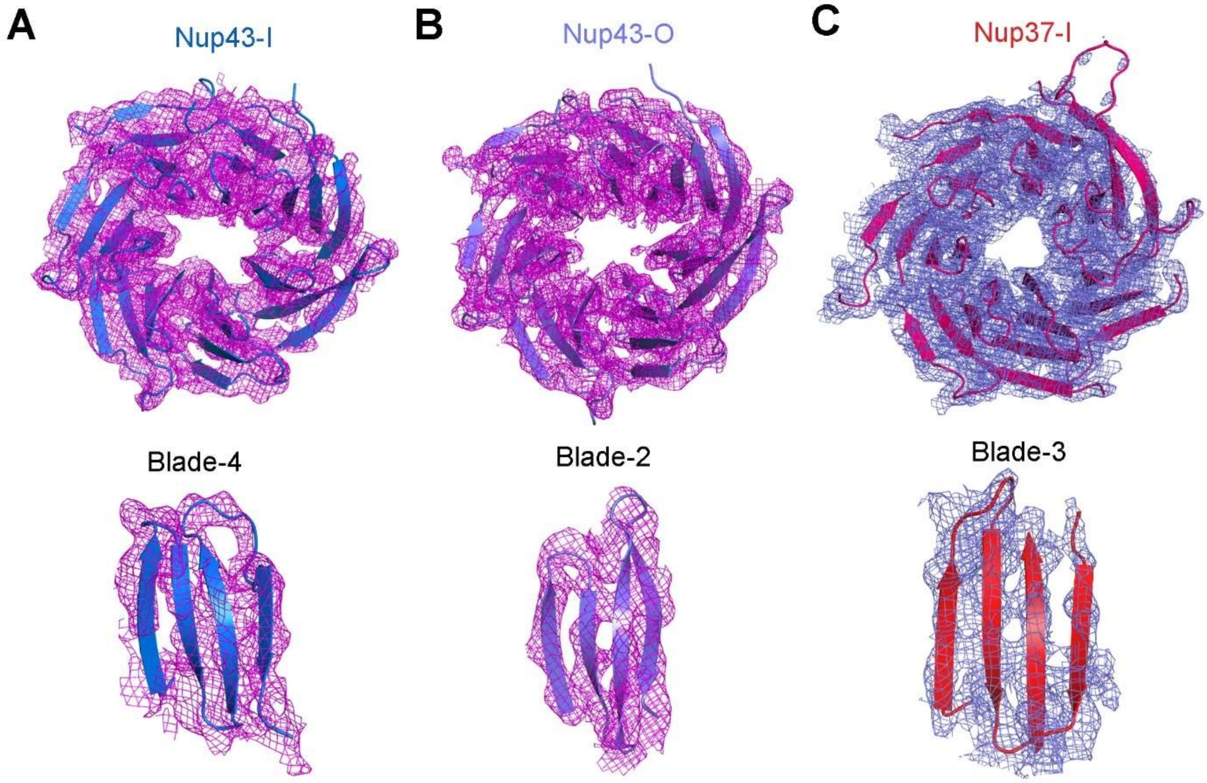


**Fig. S8.**

**Representative EM maps for Nup43 and Nup37.** (**A,B**) EM maps of the β-propeller domain from Nup43-I/O. Densities for representative blade are shown on the bottom. (**C**) EM map for the β-propeller domain of Nup37-I. The EM map for blade-3 is shown below.

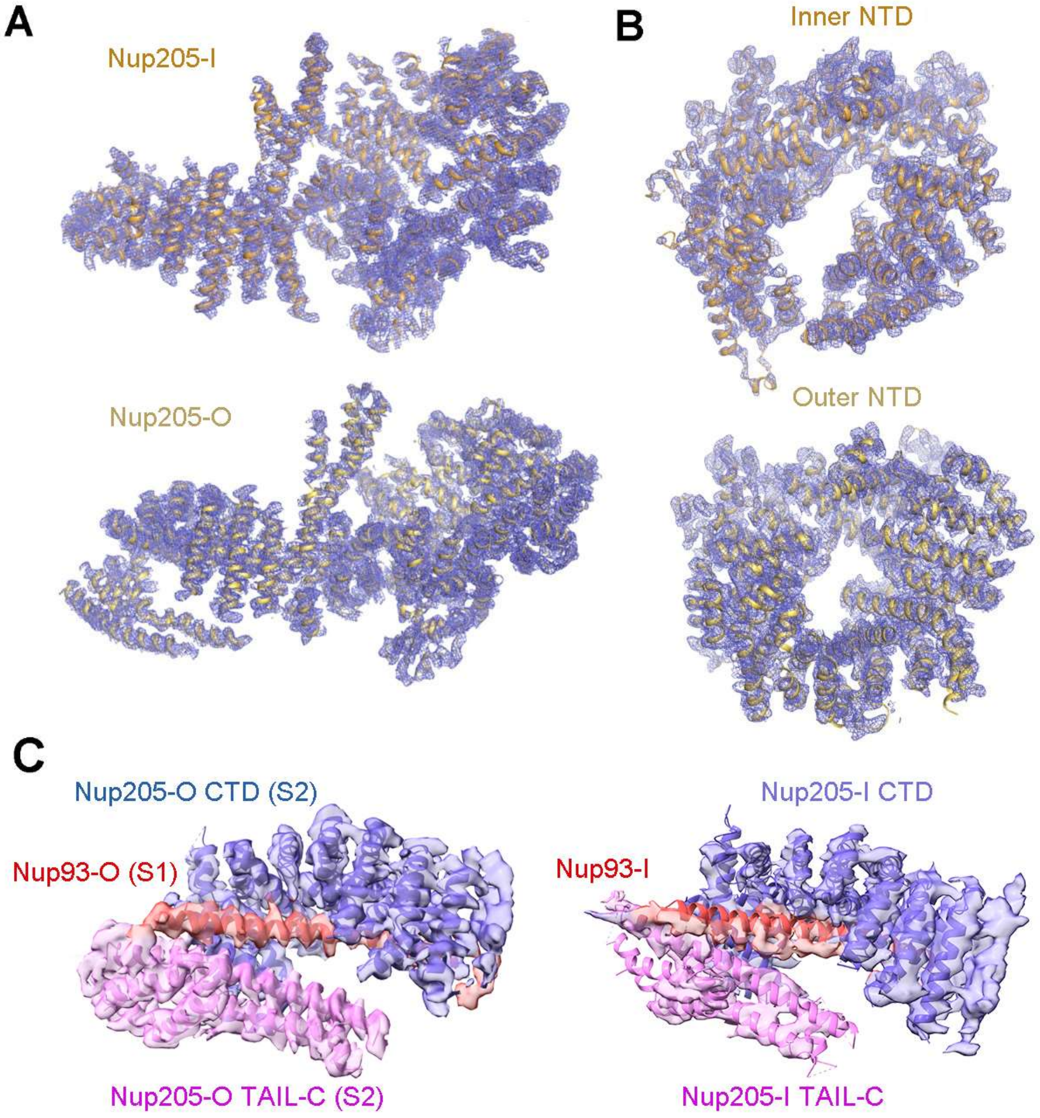


**Fig. S9.**

**Representative EM maps of Nup205.** (**A**) Overall EM maps of Nup205-I (upper panel) and Nup205-O (lower panel). (**B**) EM maps of the N-terminal domain (NTD) of Nup205-I (upper panel) and Nup205-O (lower panel). (**C**) EM maps of the CTD of Nup205-O bound to Nup93-O from the adjacent subunit (left panel) and the CTD of Nup205-I bound to Nup93-I from the same subunit (right panel). The EM maps in panel C, shown as transparent surface, are contoured at 9 σ and presented in Chimera X.

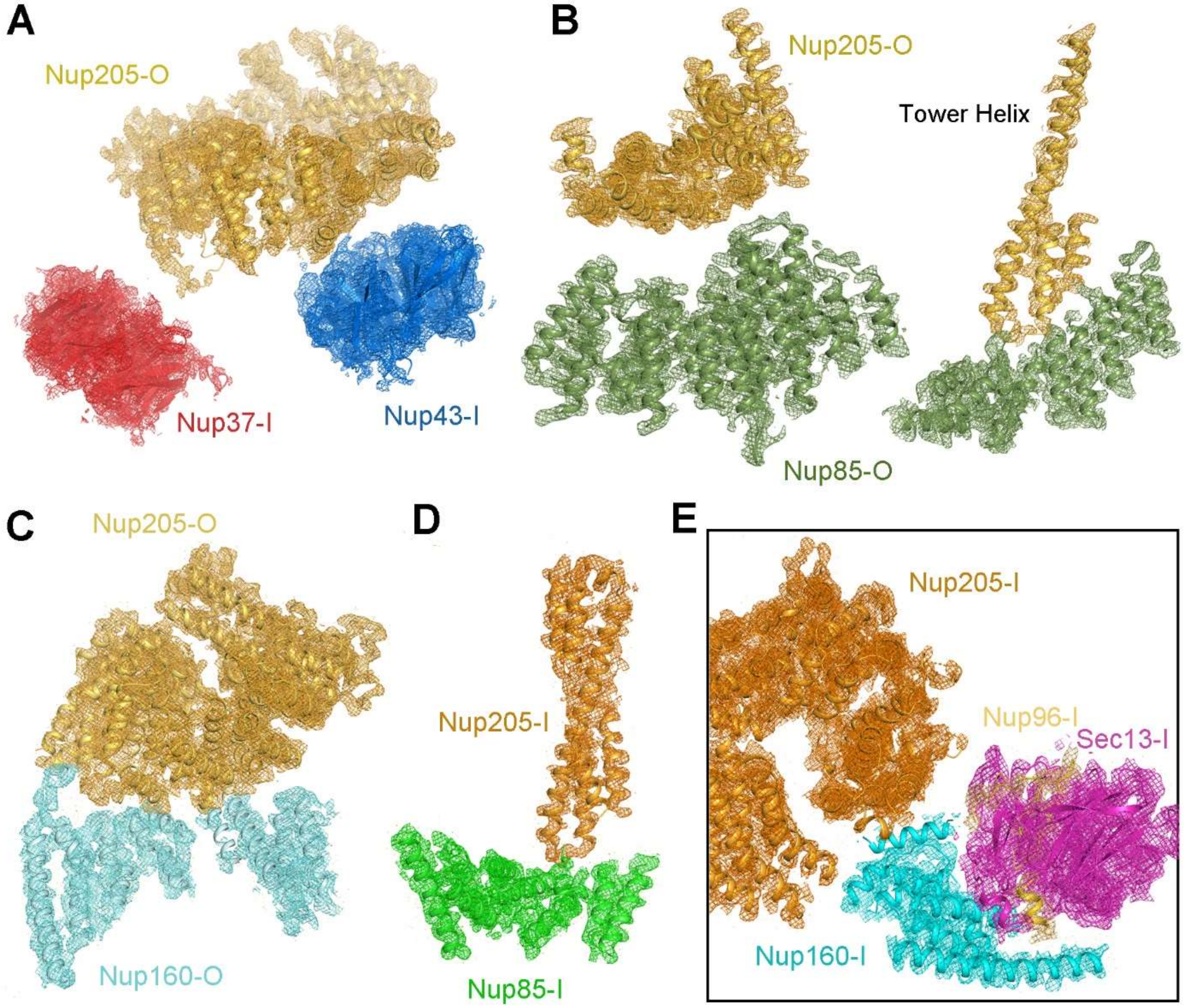


**Fig. S10.**

**EM maps of the interfaces between Nup205 and its interacting nucleoporins.** (**A**) EM map highlighting the interface of Nup205-O with Nup43-I or Nup37-I. (**B**) EM maps highlighting the interface between Nup205-O and Nup85-O. The densities for the association between the TAIL of Nup205-O and Nup85-O (left) and that between the Tower helix of Nup205-O and the α-solenoid of Nup85-O (right) are shown. (**C**) EM map highlighting the interface between Nup205-O and Nup160-O. (**D**) EM map highlighting the interface between the Tower helix of Nup205-I and the α-solenoid of Nup85-I. (**E**) EM map highlighting the association of Nup205-I, Nup160-I, Nup96-I, and Sec13-I.

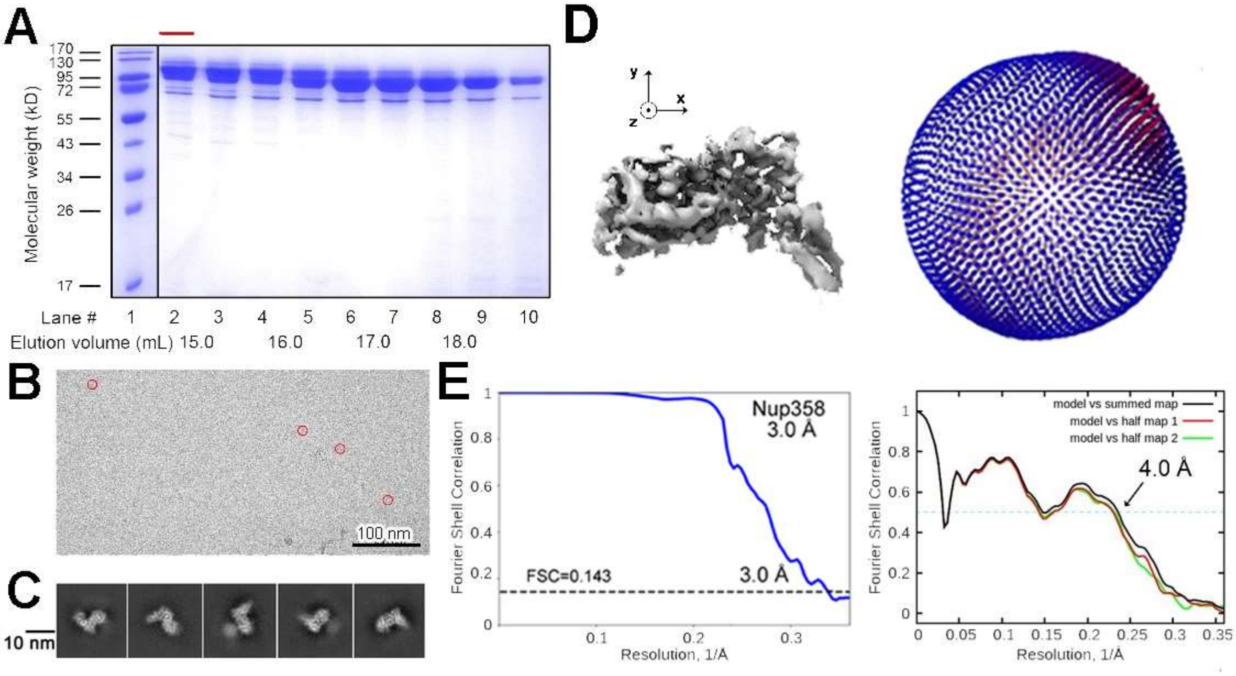


**Fig. S11.**

**Cryo-EM analysis of the N-terminal fragment (NTF) of Nup358.** (**A**) Purification of Nup358-NTF (residues 1-1171). Following gel filtration, the peak fractions were visualized on SDS-PAGE by Coomassie blue staining. Fraction 2 (indicated by red bar) was used for cryo-sample preparation. (**B**) A representative EM micrograph of Nup358-NTF. (**C**) Representative 2D class averages of Nup358-NTF. Scale bar, 10 nm. (**D**) Angular distribution of the final reconstruction. (**E**) The FSC curves. Shown on the left is the final FSC curve for Nup358-NTD2, and on the right are the FSC curves of the final reconstruction against the refined atomic coordinates. The coordinates were first refined against the map from the whole data set and the map-vs-model FSC was calculated against the first half map (green), second half map (red) and final map (black). Small differences among the three FSC curves suggest there is no apparent overfitting in the final model.

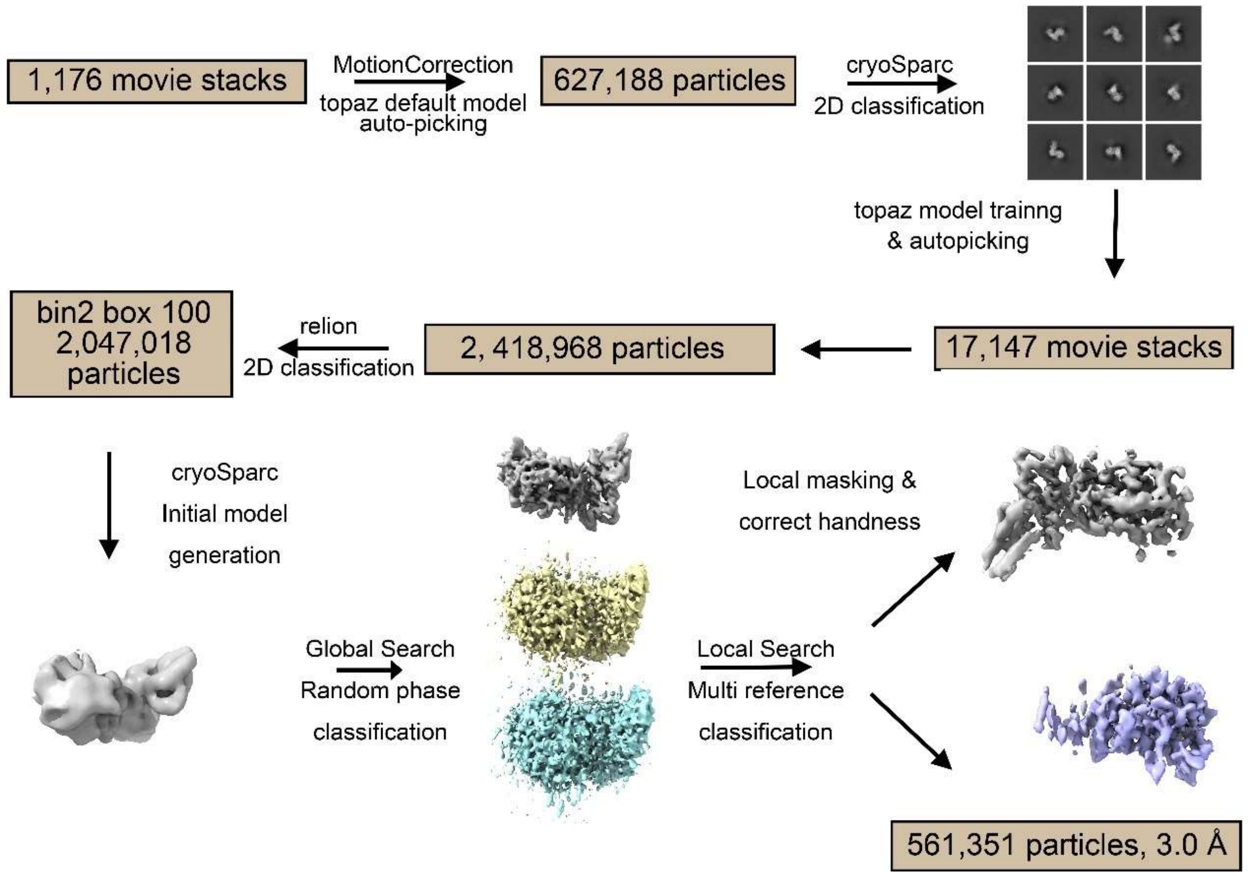


**Fig. S12.**

**Cryo-EM data processing for Nup358-NTF.** For details, please refer to the section "Image processing for Nup358-NTF" in Material and Methods.

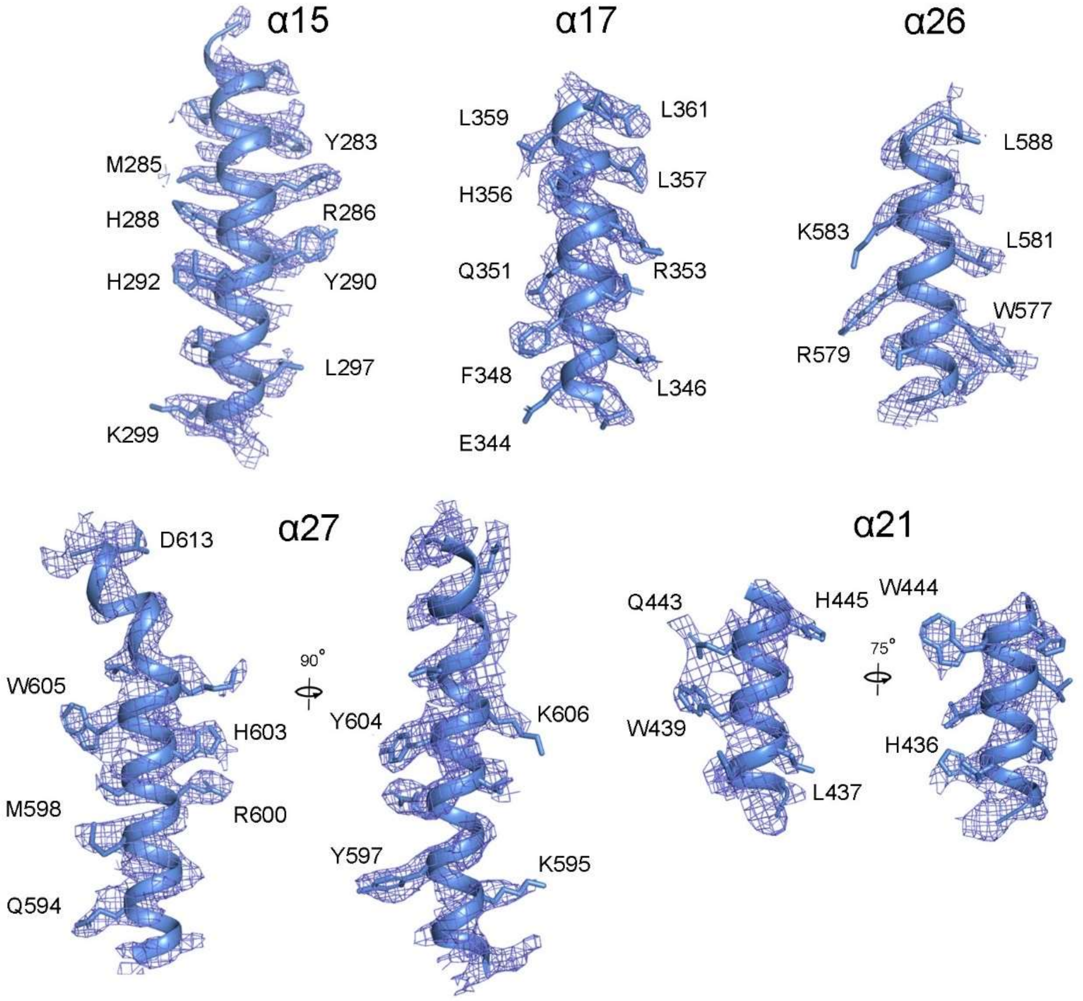


**Fig. S13.**
**Representative EM maps for Nup358-NTD2.**

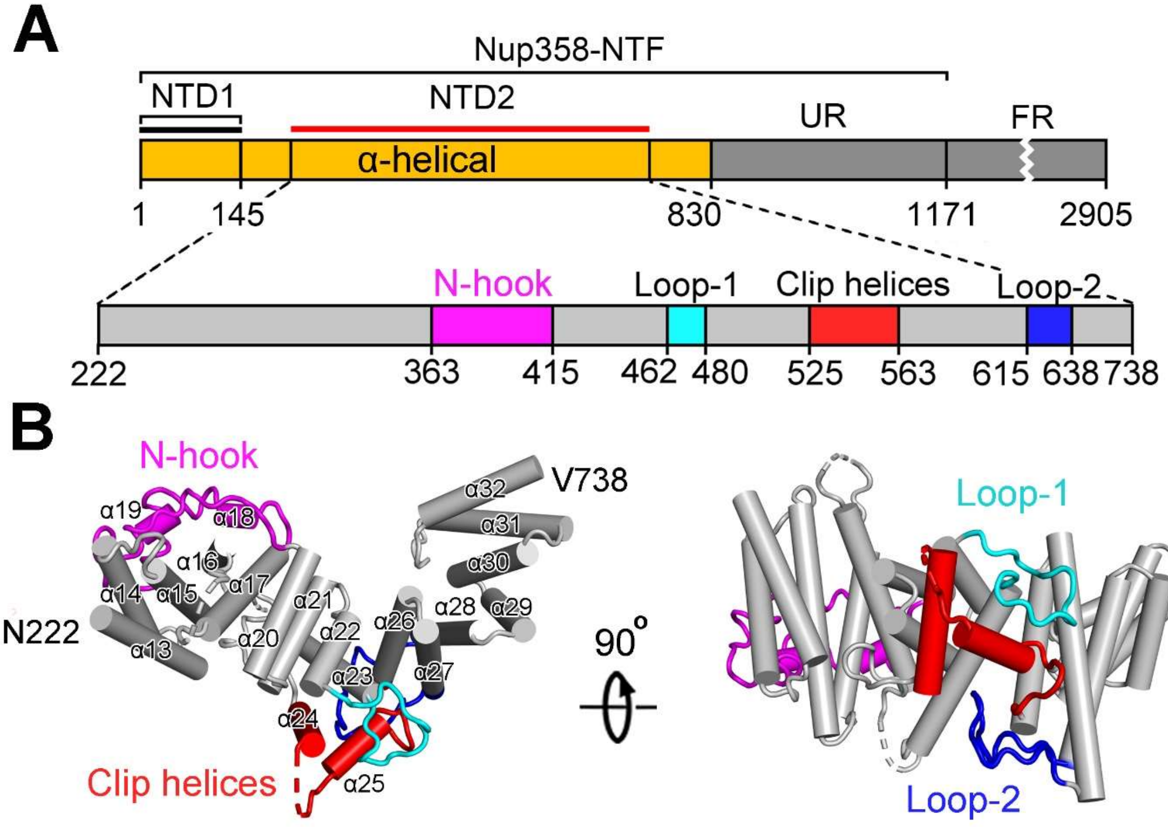


**Fig. S14.**

**Structure of Nup358-NTD2.** (**A**) The topological structure of Nup358-NTF. Nup358-NTF contains NTD1 and NTD2. NTD2 (residues 222-738) is well-defined in the EM map of Nup358-NTF. Four distinct structural motifs revealed by the EM structure, N-hook, Loop-1, Clip helices, and Loop-2, are highlighted. (**B**) Overall structure of Nup358-NTD2. Two perpendicular views are shown.

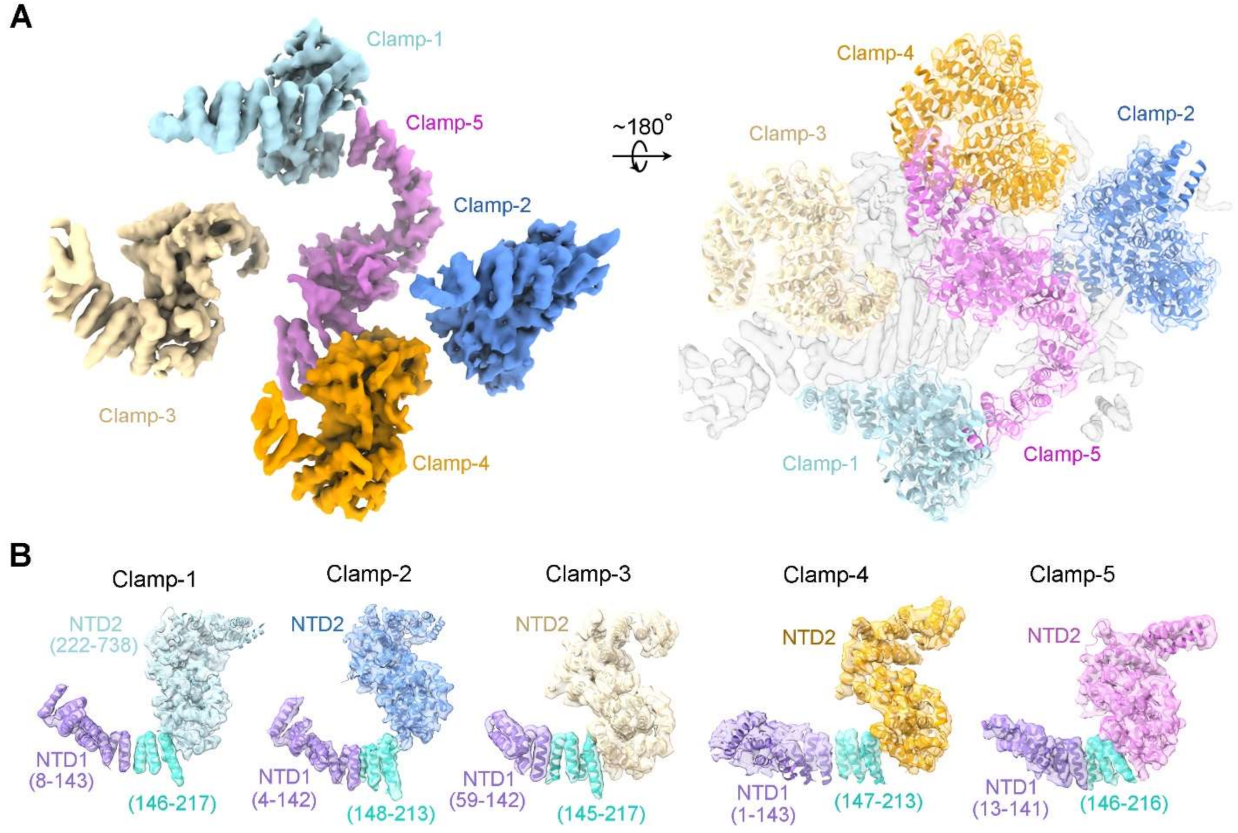


**Fig. S15.**

**EM maps for the individual clamps of the Nup358 region.** (**A**) Five Nup358 molecules are revealed in the EM map. Color-coded maps for the five clamps are shown on the left, and structures of five Nup358 docked into the 4.7-Å EM map of the Nup358 region, presented as semi-transparent surface, are shown on the right. The two view are related by a rotation of approximately 180 degrees. (**B**) Generation of the atomic models for the five Nup358 clamps. The atomic model for each clamp comprises three segments: NTD1 (residues 1-145), linker helices (residues 146-221), and NTD2 (residues 222-738). The exact boundary of modeled sequence for each clamp is adjusted based on the 4.7-Å EM map of the Nup358 region.

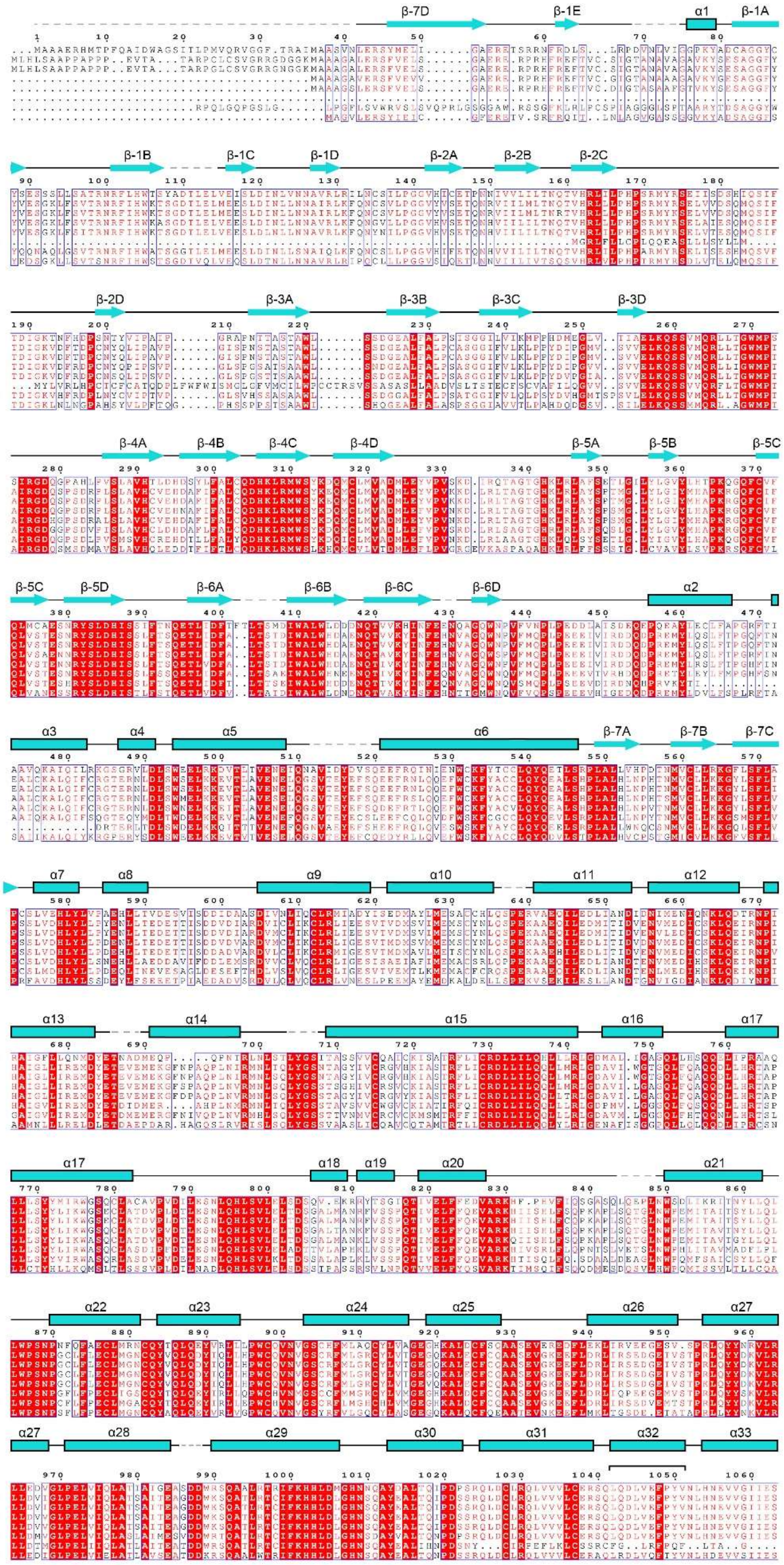
Xl_Nup160
Hs_Nup160
Pt_Nup160
Bt_Nup160
Mm_Nup160
Mg_Nup160
Ac_Nup160
Dr_Nup160

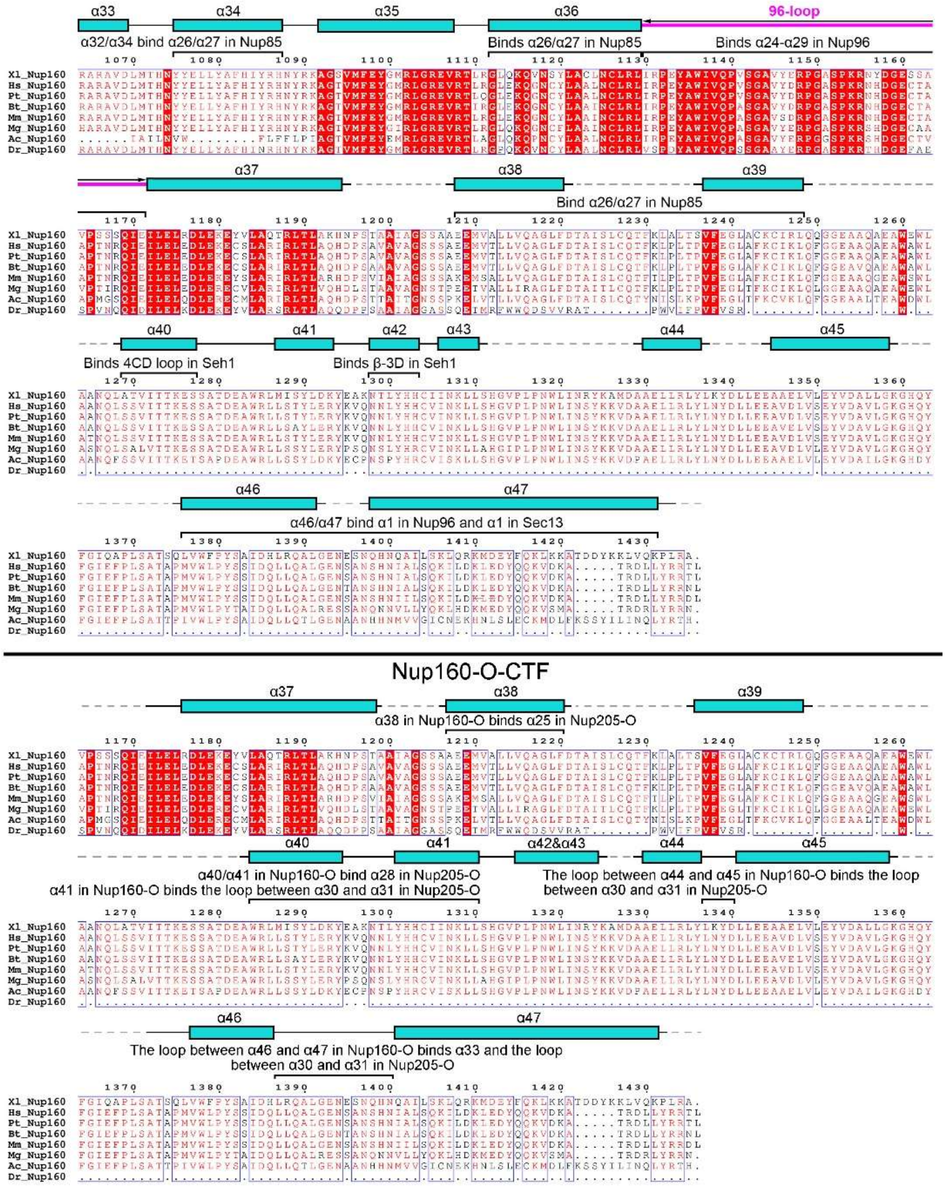

**Fig. S16.**

**Sequence alignment of Nup160 orthologues from *Xenopus laevis* (Xl), *Homo sapiens* (Hs), *Pan troglodytes* (Pt), *Bos taurus* (Bt), *Mus musculus* (Mm), *Meleagris gallopavo* (Mg), *Anolis carolinensis* (Ac), and *Danio rerio* (Dr).** Sequence alignment of full-length Nup160 is shown in the upper panel. Conserved amino acids are boxed, with invariant residues highlighted in red background. The observed secondary structural elements in Nup160-I and Nup160-O-CTF are indicated above the sequences in the upper and lower panel, respectively. Structural elements interacting with other nucleoporins in the CR subunit are indicated in both panels.

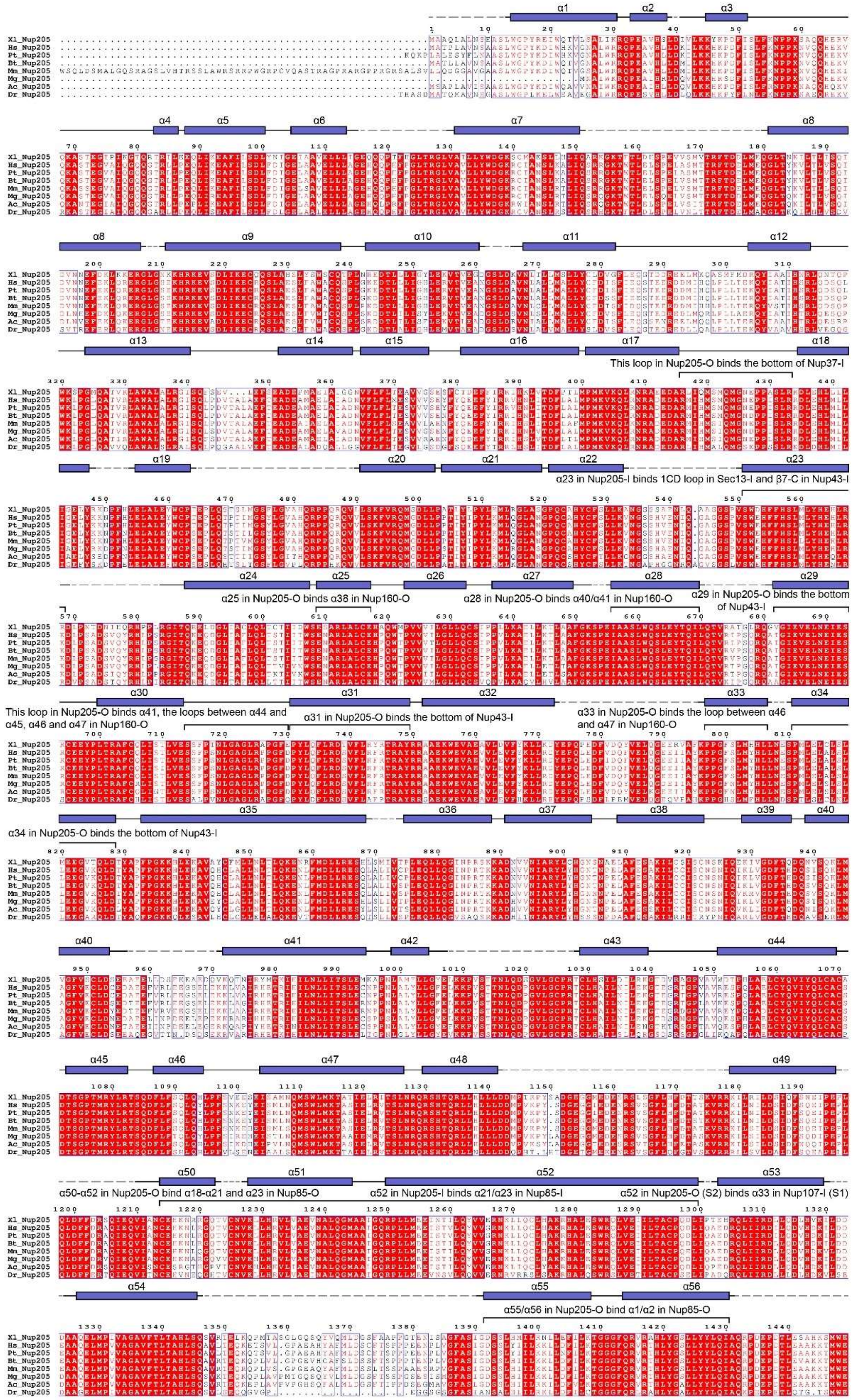
This loop in Nup205-O binds the bottom of Nup37-I
α23 in Nup205-I binds 1CD loop in Sec13-I and β7-C in Nup43-I
α25 in Nup205-O binds α38 in Nup160-O
α28 in Nup205-O binds α40/α41 in Nup160-O
α29 in Nup205-O binds the bottom of Nup43-I
This loop in Nup205-O binds α41, the loops between α44 and α45, α46 and α47 in Nup160-O
α31 in Nup205-O binds the bottom of Nup43-I
α33 in Nup205-O binds the loop between α46 and α47 in Nup160-O
α34 in Nup205-O binds the bottom of Nup43-I
α50-α52 in Nup205-O bind α18-α21 and α23 in Nup85-O
α52 in Nup205-I binds α21/α23 in Nup85-I
α52 in Nup205-O (S2) binds α33 in Nup107-I (S1)
α55/α56 in Nup205-O bind α1/α2 in Nup85-O

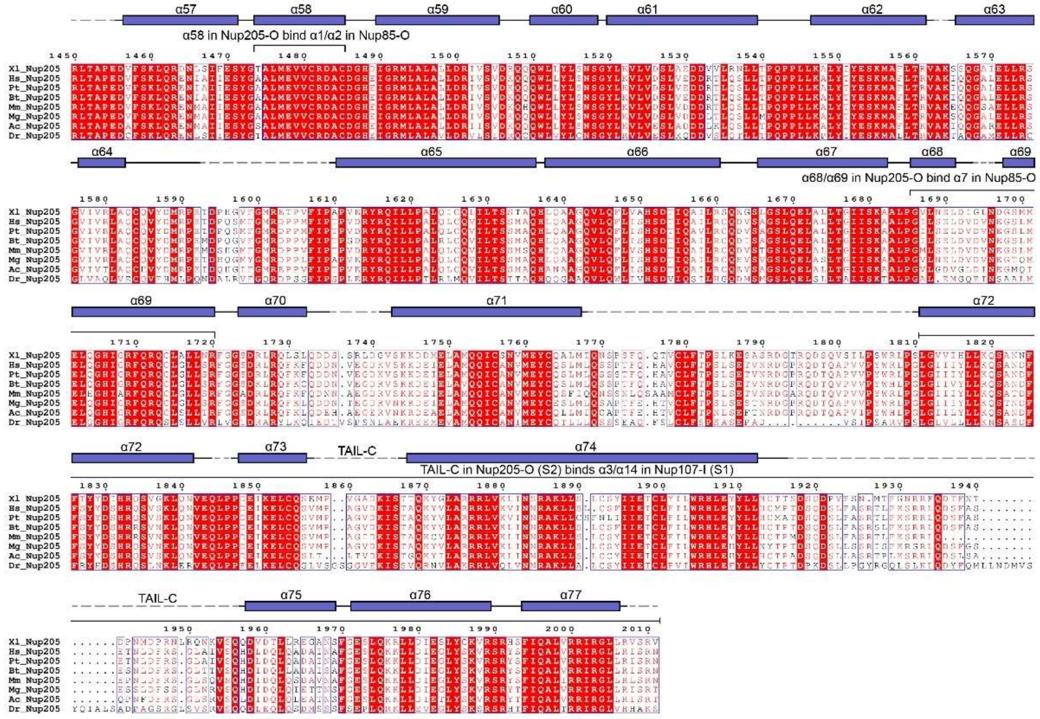


**Fig. S17.**

**Sequence alignment of Nup205 orthologues.** Conserved amino acids are boxed. Invariant residues are highlighted in red background. The observed secondary structural elements in Nup205-O are indicated above the sequences. The structural elements interacting with other nucleoporins in the CR subunit are also indicated.

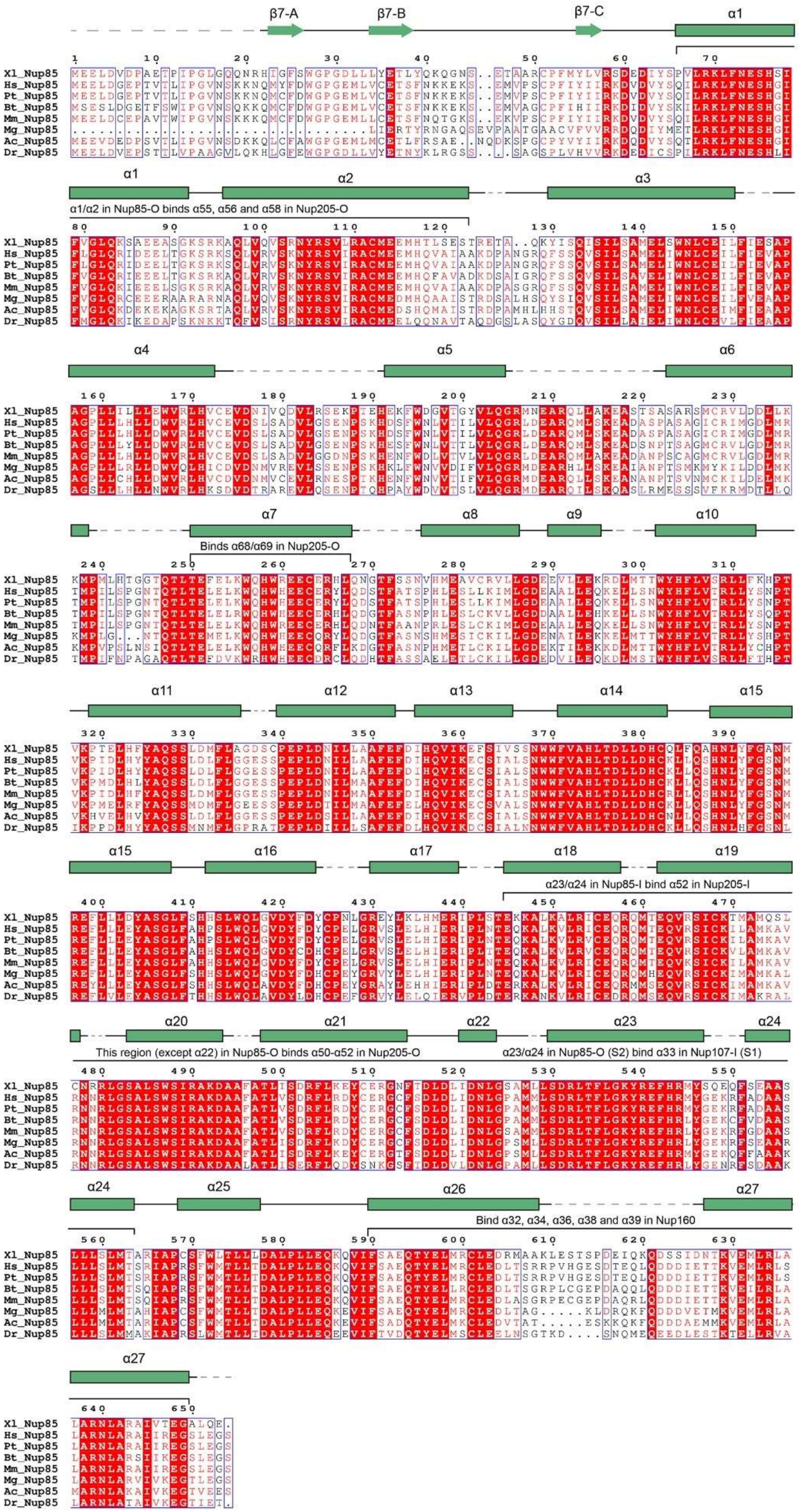
β7-A
β7-B
β7-C
α1
α1/α2 in Nup85-O binds α55, α56 and α58 in Nup205-O
Binds α68/α69 in Nup205-O
α23/α24 in Nup85-I bind α52 in Nup205-I
This region (except α22) in Nup85-O binds α50-α52 in Nup205-O
α23/α24 in Nup85-O (S2) bind α33 in Nup107-I (S1)
Bind α32, α34, α36, α38 and α39 in Nup160
Xl_Nup85
Hs_Nup85
Pt_Nup85
Bt_Nup85
Mm_Nup85
Mg_Nup85
Ac_Nup85
Dr_Nup85

**Fig. S18.**

**Sequence alignment of Nup85 orthologues.** Conserved amino acids are boxed. Invariant residues are highlighted in red background. The observed secondary structural elements in Nup85-I are indicated above the sequences. The structural elements interacting with other nucleoporins in the CR subunit are also indicated.

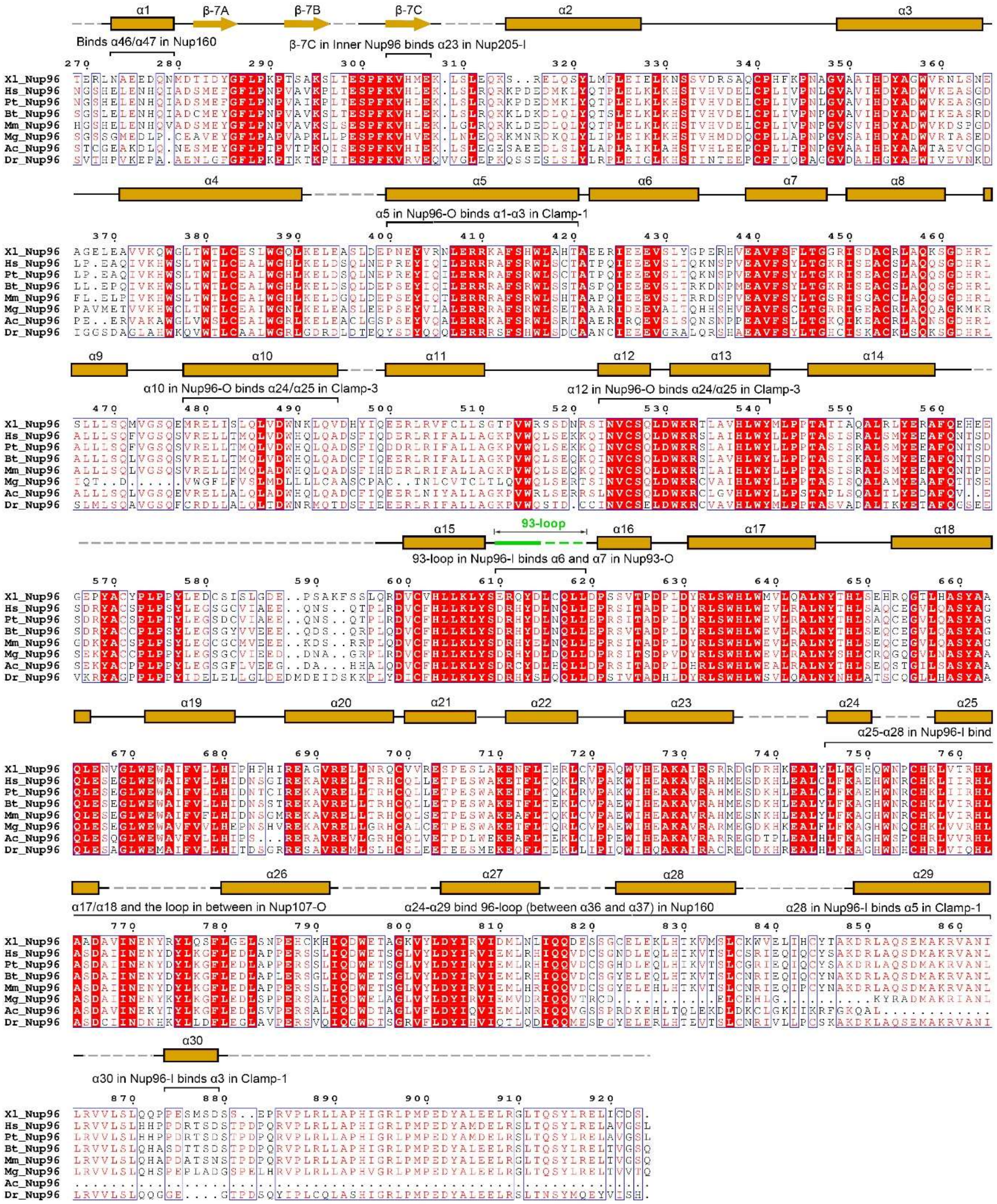


**Fig. S19.**

**Sequence alignment of Nup96 orthologues.** Conserved amino acids are boxed. Invariant residues are highlighted in red background. The observed secondary structural elements in Nup96-I are indicated above the sequences. The structural elements interacting with other nucleoporins in the CR subunit are also indicated.

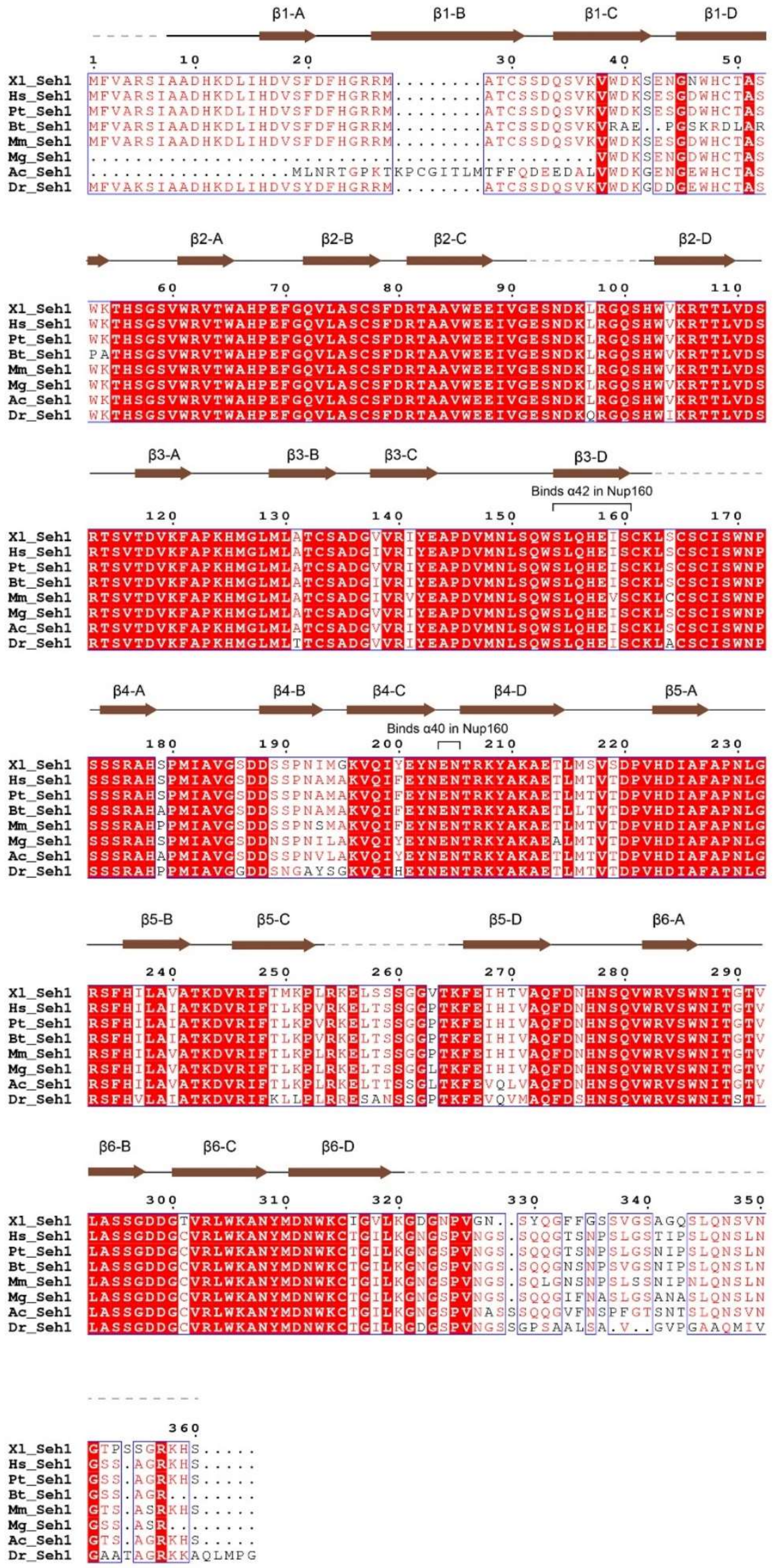
β1-A
β1-B
β1-C
β1-D
β2-A
β2-B
β2-C
β2-D
β3-A
β3-B
β3-C
β3-D
Binds α42 in Nup160
β4-A
β4-B
β4-C
β4-D
β5-A
Binds α40 in Nup160
β5-B
β5-C
β5-D
β6-A
β6-B
β6-C
β6-D
Xl_Seh1
Hs_Seh1
Pt_Seh1
Bt_Seh1
Mm_Seh1
Mg_Seh1
Ac_Seh1
Dr_Seh1

**Fig. S20.**

**Sequence alignment of Seh1 orthologues.** The conserved amino acids are boxed, with invariant residues highlighted in red background. The observed secondary structural elements in Seh1-I are indicated above the sequences. The structural elements interacting with other nucleoporins in the CR subunit are also indicated.

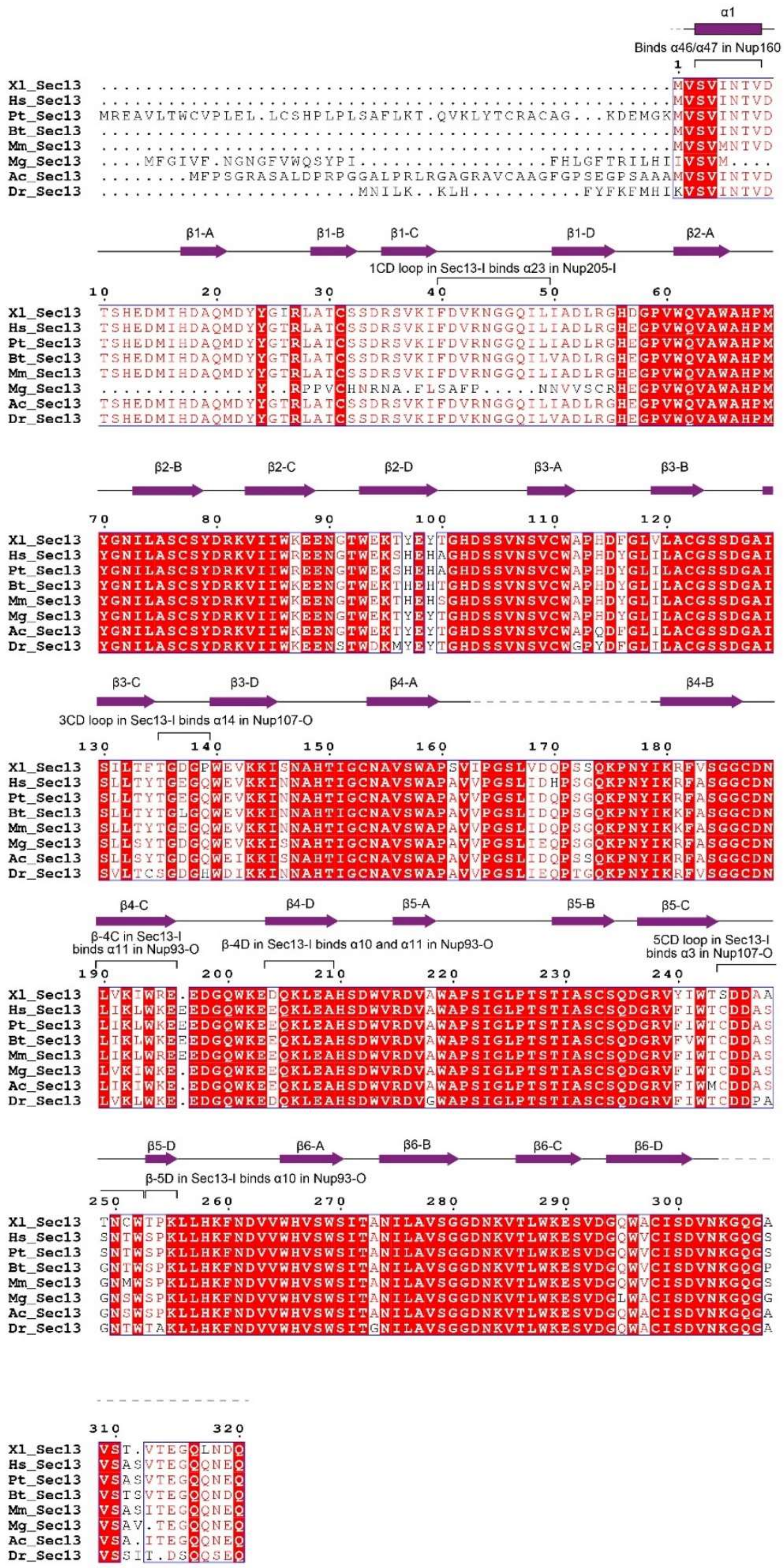
α1
Binds α46/α47 in Nup160
β1-A
β1-B
β1-C
β1-D
β2-A
1CD loop in Sec13-I binds α23 in Nup205-I
β2-B
β2-C
β2-D
β3-A
β3-B
β3-C
β3-D
β4-A
β4-B
3CD loop in Sec13-I binds α14 in Nup107-O
β4-C
β4-D
β5-A
β5-B
β5-C
β-4C in Sec13-I binds α11 in Nup93-O
β-4D in Sec13-I binds α10 and α11 in Nup93-O
5CD loop in Sec13-I binds α3 in Nup107-O
β5-D
β6-A
β6-B
β6-C
β6-D
β-5D in Sec13-I binds α10 in Nup93-O
Xl_Sec13
Hs_Sec13
Pt_Sec13
Bt_Sec13
Mm_Sec13
Mg_Sec13
Ac_Sec13
Dr_Sec13

**Fig. S21.**

**Sequence alignment of Sec13 orthologues.** The conserved amino acids are boxed, with invariant residues highlighted in red background. The observed secondary structural elements in Sec13-I are indicated above the sequences. The structural elements interacting with other nucleoporins in the CR subunit are also indicated.

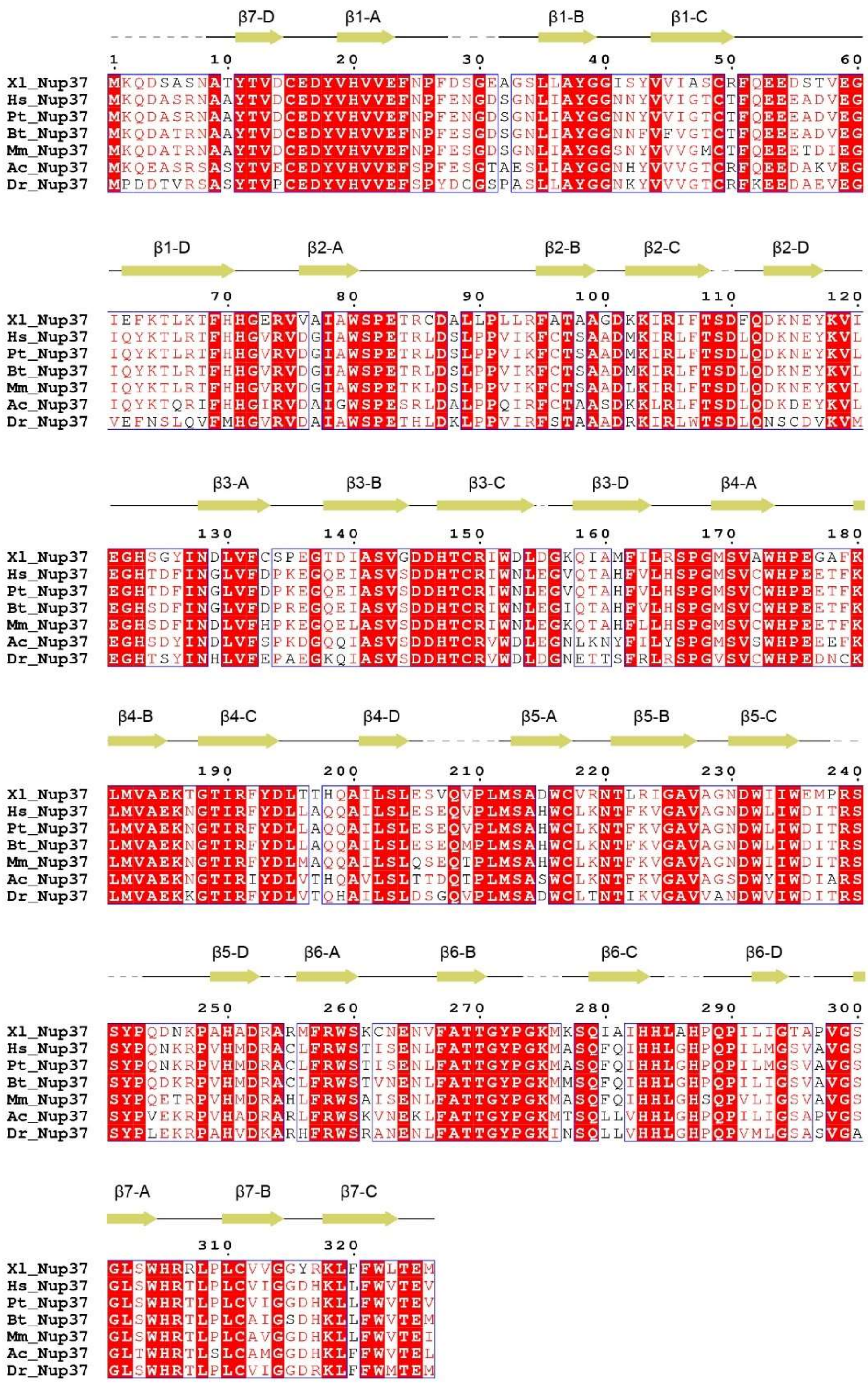


**Fig. S22.**

**Sequence alignment of Nup37 orthologues.** The conserved amino acids are boxed, with invariant residues highlighted in red background. The observed secondary structural elements in Nup37-I are indicated above the sequences.

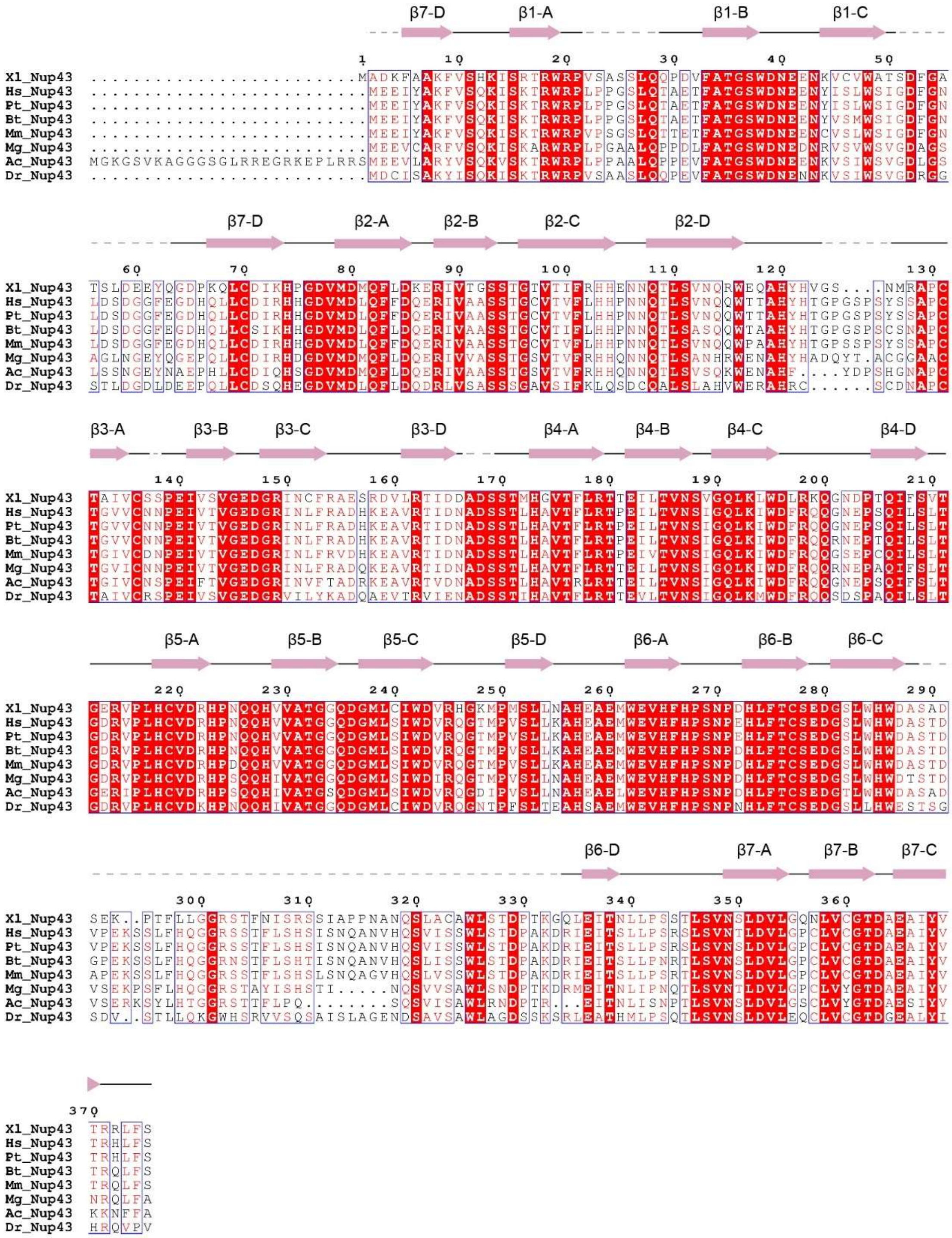

**Fig. S23.**

**Sequence alignment of Nup43 orthologues.** The conserved amino acids are boxed, with invariant residues highlighted in red background. The observed secondary structural elements in Nup43-I are indicated above the sequences.

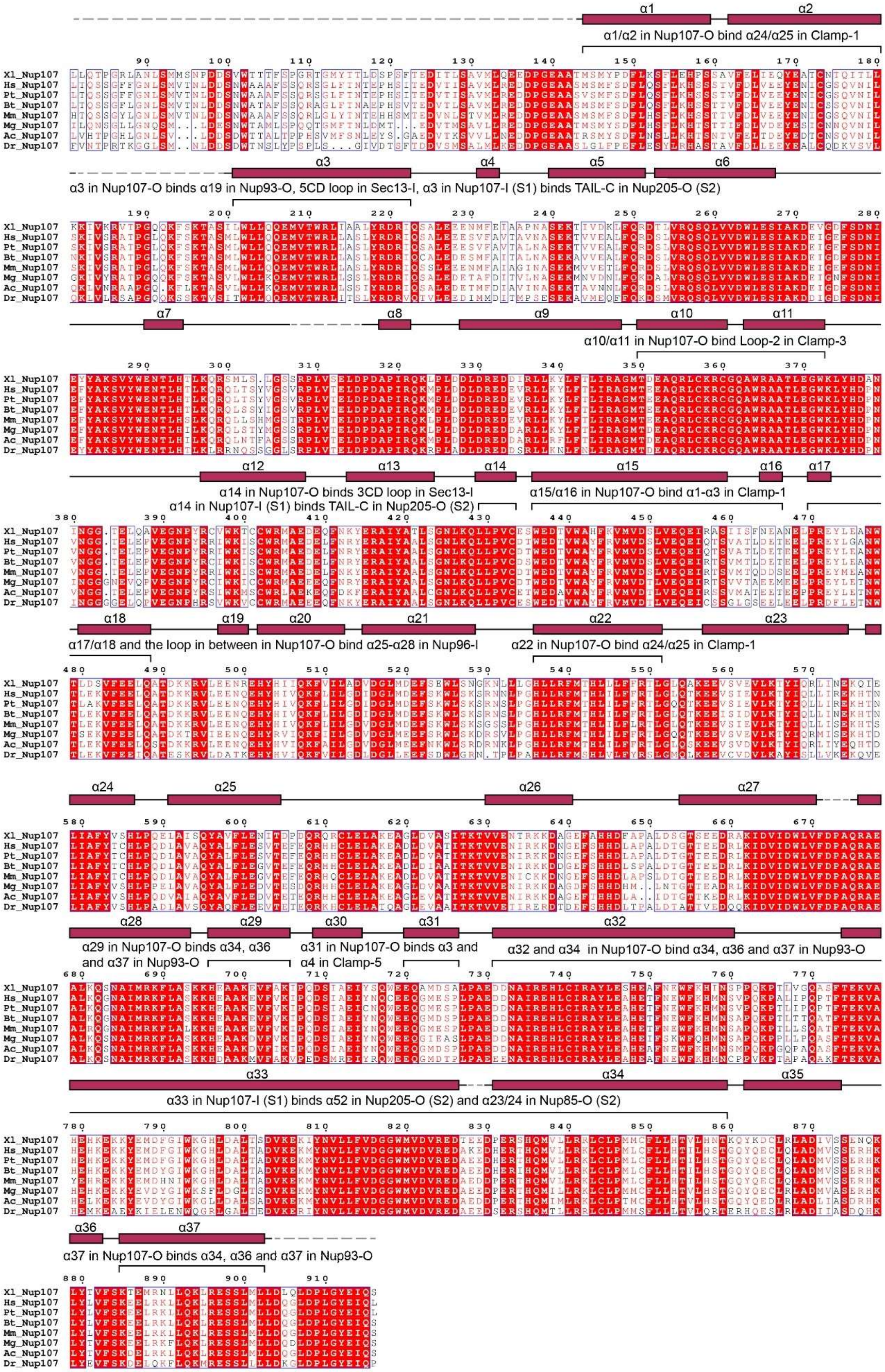
α1/α2 in Nup107-O bind α24/α25 in Clamp-1
α3 in Nup107-O binds α19 in Nup93-O, 5CD loop in Sec13-I, α3 in Nup107-I (S1) binds TAIL-C in Nup205-O (S2)
α10/α11 in Nup107-O bind Loop-2 in Clamp-3
α14 in Nup107-O binds 3CD loop in Sec13-I
α14 in Nup107-I (S1) binds TAIL-C in Nup205-O (S2)
α15/α16 in Nup107-O bind α1-α3 in Clamp-1
α17/α18 and the loop in between in Nup107-O bind α25-α28 in Nup96-I
α22 in Nup107-O bind α24/α25 in Clamp-1
α29 in Nup107-O binds α34, α36 and α37 in Nup93-O
α31 in Nup107-O binds α3 and α4 in Clamp-5
α32 and α34 in Nup107-O bind α34, α36 and α37 in Nup93-O
α33 in Nup107-I (S1) binds α52 in Nup205-O (S2) and α23/24 in Nup85-O (S2)
α37 in Nup107-O binds α34, α36 and α37 in Nup93-O
Xl_Nup107
Hs_Nup107
Pt_Nup107
Bt_Nup107
Mm_Nup107
Mg_Nup107
Ac_Nup107
Dr_Nup107

**Fig. S24.**

**Sequence alignment of Nup107 orthologues.** The conserved amino acids are boxed, with invariant residues highlighted in red background. The observed secondary structural elements in Nup107-I are indicated above the sequences. The structural elements interacting with other nucleoporins in the CR subunit are also indicated.

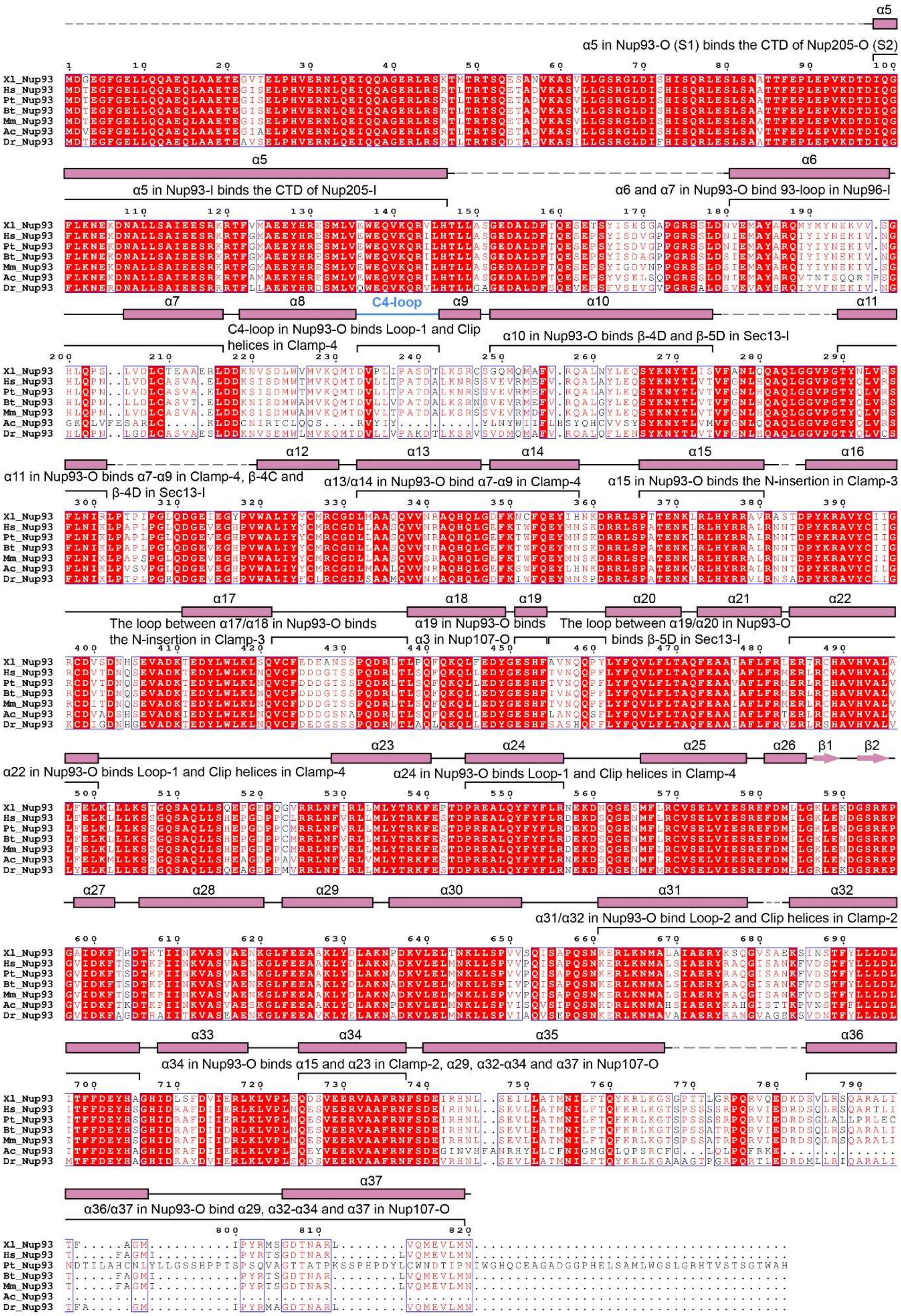
α5 in Nup93-O (S1) binds the CTD of Nup205-O (S2)
α5 in Nup93-I binds the CTD of Nup205-I
α6 and α7 in Nup93-O bind 93-loop in Nup96-I
C4-loop
C4-loop in Nup93-O binds Loop-1 and Clip helices in Clamp-4
α10 in Nup93-O binds β-4D and β-5D in Sec13-I
α11 in Nup93-O binds α7-α9 in Clamp-4, β-4C and β-4D in Sec13-I
α13/α14 in Nup93-O bind α7-α9 in Clamp-4
α15 in Nup93-O binds the N-insertion in Clamp-3
The loop between α17/α18 in Nup93-O binds the N-insertion in Clamp-3
α19 in Nup93-O binds α3 in Nup107-O
The loop between α19/α20 in Nup93-O binds β-5D in Sec13-I
α22 in Nup93-O binds Loop-1 and Clip helices in Clamp-4
α24 in Nup93-O binds Loop-1 and Clip helices in Clamp-4
α31/α32 in Nup93-O bind Loop-2 and Clip helices in Clamp-2
α34 in Nup93-O binds α15 and α23 in Clamp-2, α29, α32-α34 and α37 in Nup107-O
α36/α37 in Nup93-O bind α29, α32-α34 and α37 in Nup107-O
Xl_Nup93
Hs_Nup93
Pt_Nup93
Bt_Nup93
Mm_Nup93
Ac_Nup93
Dr_Nup93

**Fig. S25.**

**Sequence Alignment of Nup93 orthologs.** Conserved amino acids are boxed. Invariant residues are highlighted in red background. The observed secondary structural elements in Nup93-O are indicated above the sequences. The structural elements interacting with other nucleoporins in the CR subunit are also indicated.

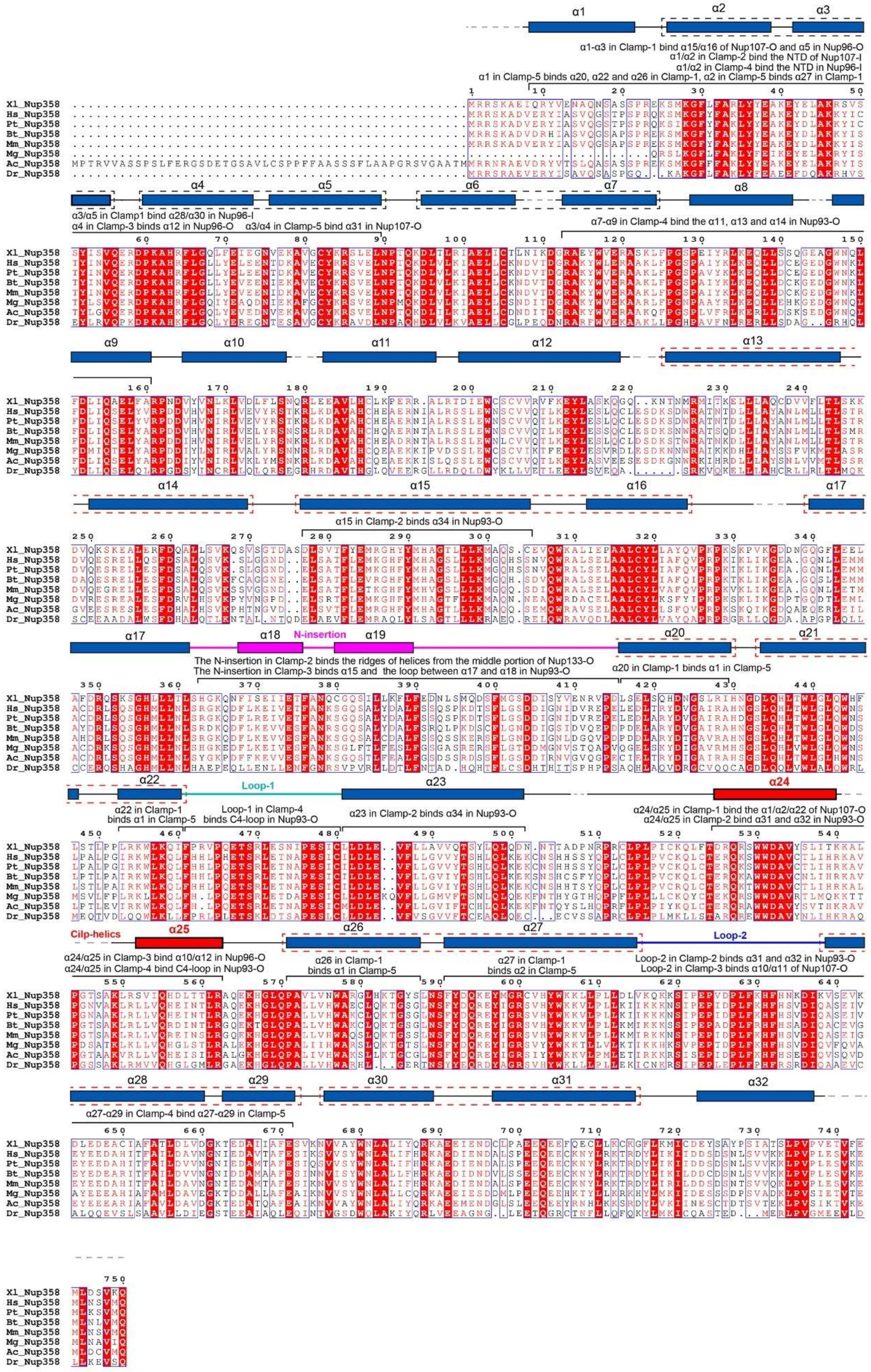
α1-α3 in Clamp-1 bind α15/α16 of Nup107-O and α5 in Nup96-O
α1/α2 in Clamp-2 bind the NTD of Nup107-I
α1/α2 in Clamp-4 bind the NTD in Nup96-I
α1 in Clamp-5 binds α20, α22 and α26 in Clamp-1, α2 in Clamp-5 binds α27 in Clamp-1
α3/α5 in Clamp1 bind α28/α30 in Nup96-I
α4 in Clamp-3 binds α12 in Nup96-O
α3/α4 in Clamp-5 bind α31 in Nup107-O
α7-α9 in Clamp-4 bind the α11, α13 and α14 in Nup93-O
α15 in Clamp-2 binds α34 in Nup93-O
N-insertion
The N-insertion in Clamp-2 binds the ridges of helices from the middle portion of Nup133-O
The N-insertion in Clamp-3 binds α15 and the loop between α17 and α18 in Nup93-O
α20 in Clamp-1 binds α1 in Clamp-5
Loop-1
α22 in Clamp-1 binds α1 in Clamp-5
Loop-1 in Clamp-4 binds C4-loop in Nup93-O
α23 in Clamp-2 binds α34 in Nup93-O
α24/α25 in Clamp-1 bind the α1/α2/α22 of Nup107-O
α24/α25 in Clamp-2 bind α31 and α32 in Nup93-O
Cilp-helics
α24/α25 in Clamp-3 bind α10/α12 in Nup96-O
α24/α25 in Clamp-4 bind C4-loop in Nup93-O
α26 in Clamp-1 binds α1 in Clamp-5
α27 in Clamp-1 binds α2 in Clamp-5
Loop-2
Loop-2 in Clamp-2 binds α31 and α32 in Nup93-O
Loop-2 in Clamp-3 binds α10/α11 of Nup107-O
α27-α29 in Clamp-4 bind α27-α29 in Clamp-5
Xl_Nup358
Hs_Nup358
Pt_Nup358
Bt_Nup358
Mm_Nup358
Mg_Nup358
Ac_Nup358
Dr_Nup358

**Fig. S26.**

**Sequence alignment of Nup358-NTF among Nup358 orthologues.** Conserved amino acids are boxed. Invariant residues are highlighted in red background. The observed secondary structural elements in Clamp-1 are indicated above the sequences. The TPR helix pairs found in previous human Nup358-NTD1 structure (*33*) and our *X. laevis* Nup358-NTD2 structure are indicated by black and red dashed boxes, respectively. It should be noted that helices α17 and α20 are not continuous in sequence, but still form a TPR like helical pair. The structural elements interacting with other nucleoporins in the CR subunit are also indicated.

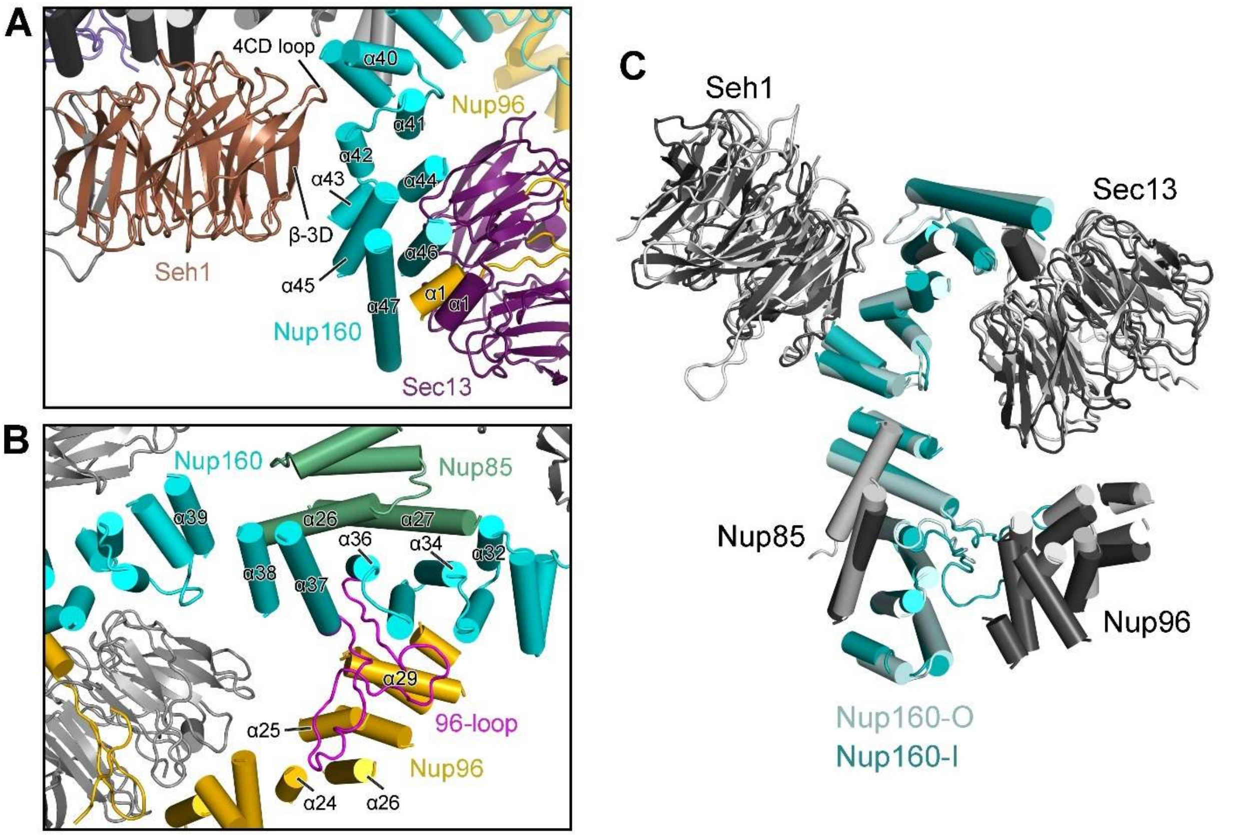


**Fig. S27.**

**Nup160-CTF as an organizing center for the vertex of the Y complex.** (**A**) Nup160-CTF is sandwiched by the two β-propellers from Seh1 and Sec13. Shown here is the inner complex. The 4CD loop (the loop between strands C and D of blade 4) of Seh1 contacts helix α40 of Nup160-CTF, and the β-3D strand (the strand D of blade 3) of Seh1 interacts with helix α42 of Nup160-CTF. The N-terminal α1 helices from Nup96 and Sec13 pair up to wedge into the crevice between helices α46 and α47 of Nup160. These interactions may strengthen the Y complex. This panel is related to Fig. 2C. (**B**) The middle portion of Nup160-I is sandwiched by Nup85-I and Nup96-I. The C-terminal helices of Nup96-I make no direct contact to the α-solenoid of Nup160-I. Instead, a conserved surface loop between α36 and α37 of Nup160-I referred to as 96-loop closely associates with the ridge of helices α24-α29 from Nup96-I. Similar to that in fungi (*16, 17*), two C-terminal helices α26 and α27 of Nup85-I contact the lateral side of helices α32-α39 from Nup160-I. (**C**) Comparison of the vertex structures from the inner and outer Y complexes. The two vertexes are superimposed on their respective Nup160-CTF, with an RMSD of 1.9 Å over 237 aligned Cα atoms. The interactions described for inner Y complex are also recapitulated in the outer Y complex.

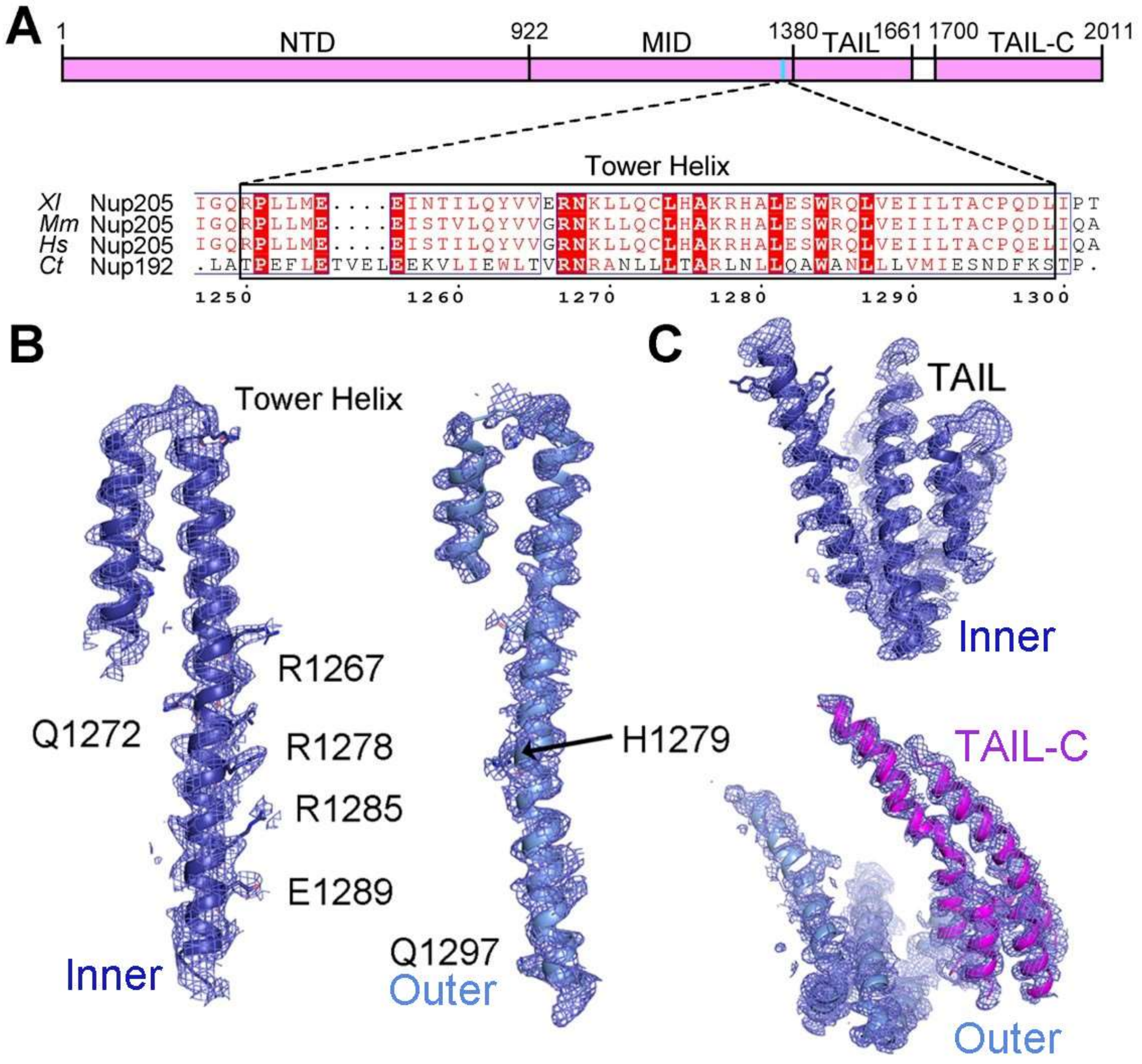


**Fig. S28.**

**Identification of two Nup205 molecules in the CR subunit.** (**A**) Sequence features of Nup205 from *X. laevis*. Shown here is a schematic diagram of Nup205 domain organization, with sequence alignment of the Tower helix region displayed below. The aligned sequences include those from *X. laevis* (Xl), *Mus musculus* (Mm), *Homo sapiens* (Hs) and *C. thermophilum* (Ct). Conserved residues are boxed. (**B**) Two Tower helices are defined by the EM maps and assigned to Nup205-I (left) and Nup205-O (right). (**C**) Identification of the C-terminal helical domains of Nup205. The fine EM maps allow atomic modeling of the TAIL domain in both Nup205-I (upper panel) and Nup205-O (lower panel). In addition, the EM maps allow identification of the TAIL-C domain in Nup205-O. All EM maps in this figure were prepared using the reconstruction of the core region with a contour level between 5 – 10 σ.

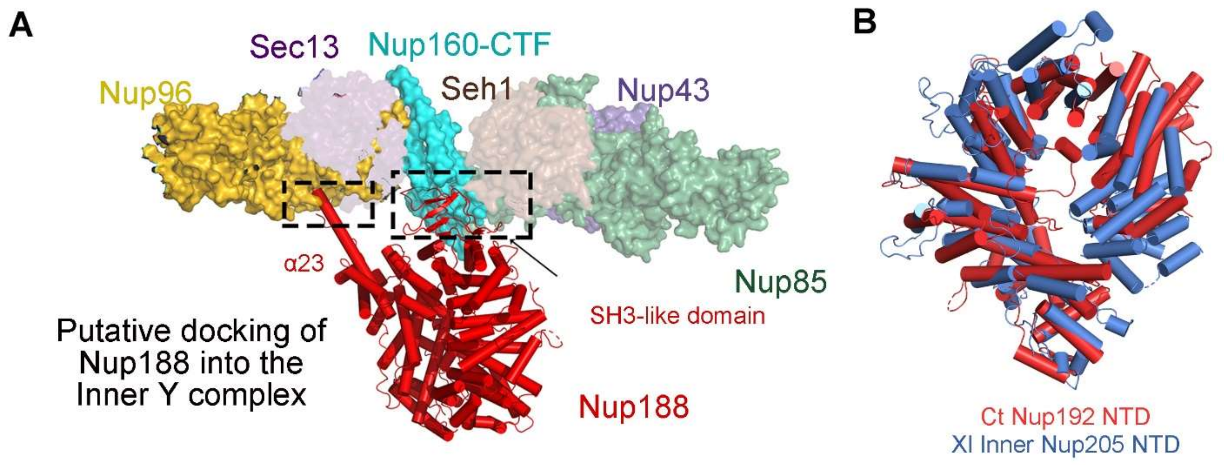


**Fig. S29.**

**Structural comparison of Nup205 with Nup188.** (**A**) Features of the EM map is incompatible with assignment of Nup188. Docking of the X-ray structure of Nup188 from *M. thermophila* (MtNup188) into the EM map of the core region results in significant steric clashes in two areas (marked by dashed boxes). In particular, the conserved SH3-like domain of Nup188 (*31*) clashes with Nup160-I-CTF. (**B**) Structural comparison of Nup205 and CtNup192. The crystal structures of CtNup192 NTD (Nup205 ortholog in *Chaetomium thermophilum*, PDB code: 4KNH) is aligned to the NTD from Nup205-I.

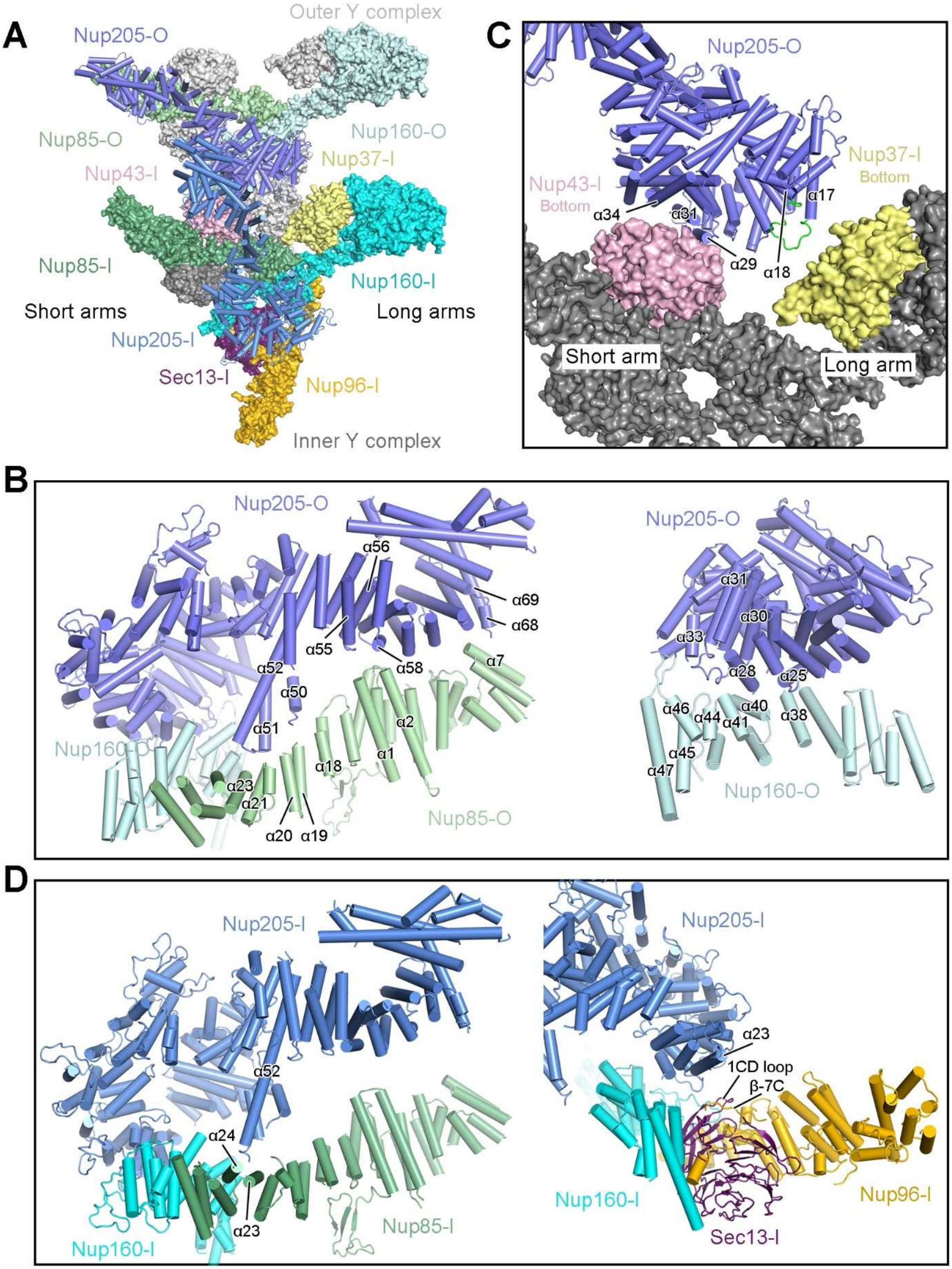
A
Outer Y complex
Nup205-O
Nup160-O
Nup85-O
Nup37-I
Nup43-I
Nup85-I
Nup160-I
Short arms
Long arms
Nup205-I
Sec13-I
Nup96-I
Inner Y complex
C
Nup205-O
Nup37-I
Bottom
Nup43-I
Bottom
α34
α31
α17
α29
α18
Short arm
Long arm
B
Nup205-O
α56
α69
α68
α55
α58
α52
α7
α50
α51
α2
Nup160-O
α18
α1
α23
α21
α20
α19
Nup85-O
Nup205-O
α31
α30
α33
α28
α25
α46
α44
α40
α41
α38
α45
α47
Nup160-O
D
Nup205-I
α52
α24
α23
Nup85-I
Nup160-I
Nup205-I
α23
1CD loop
β-7C
Nup160-I
Nup96-I
Sec13-I

**Fig. S30.**

**Nup205-I and Nup205-O associate with the Y complexes through different interfaces.** (**A**) Nup205-O connects inner and outer Y complexes. Nup205-O interacts with the short arm and the vertex of outer Y complex; it also associates with both arms of inner Y complex. Nup205-I mainly associates with the short arm and the vertex of inner Y complex. (**B**) Two close-up views on the interface between Nup205-O and the outer Y complex. On one side, the Tower helix α52 and six additional helices α50/51/55/56/58/68/69 associate with the ridge of the α-solenoid of Nup85-O in the short arm (left panel). On the other side, five helices α25/28/30/31/33 contact Nup160-O-CTF at the vertex (right panel). (**C**) Association between Nup205-O and the inner Y complex. Three helices α29/31/34 of Nup205-O closely associate with the bottom face of the β-propeller of Nup43-I. A flexible loop between helices α17 and α18 (colored green) is located in close proximity to the bottom face of the β-propeller of Nup37-I. (**D**) Two close-up views on the interface between Nup205-I and the inner Y complex. The Tower helix α52 of Nup205-I associates with helices α23/α24 of Nup85-I in the short arm (left panel). At the interface with the vertex of the inner Y complex, helix α23 of Nup205-I is positioned in the vicinity of the 1CD loop of Sec13-I and strand β-7C of Nup96-I (right panel).

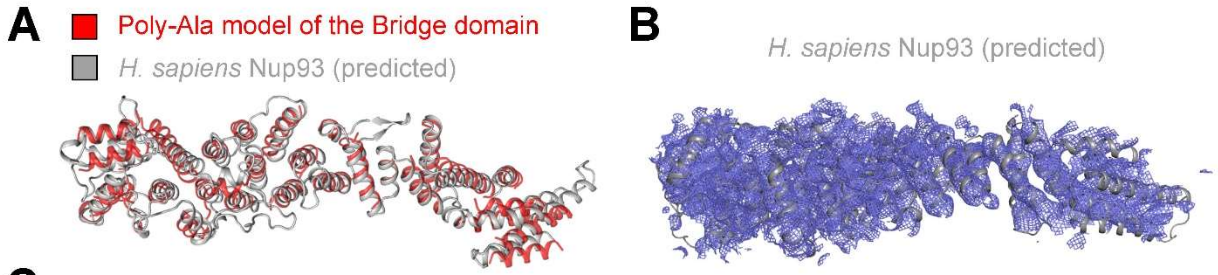


C

| Protein A | Position | Protein B | Position |
|---|---|---|---|
| Nup93 | 371 | Nup96 | 94 |
| Nup93 | 559 | Nup358 | 607 |
| Nup93 | 665 | Nup358 | 808 |
| Nup93 | 590 | Nup358 | 2230 |
| Nup93 | 590 | Nup358 | 2746 |
| Nup93 | 782 | Nup358 | 1207 |
| Nup93 | 782 | Nup358 | 2230 |

| Protein A | Position | Protein B | Position |
|---|---|---|---|
| Nup93 | 103 | TPR | 1351 |
| Nup93 | 388 | TPR | 45 |
| Nup93 | 590 | TPR | 413 |
| Nup93 | 590 | TPR | 563 |
| Nup93 | 782 | TPR | 1023 |
| Nup93 | 782 | ELYS | 2193 |

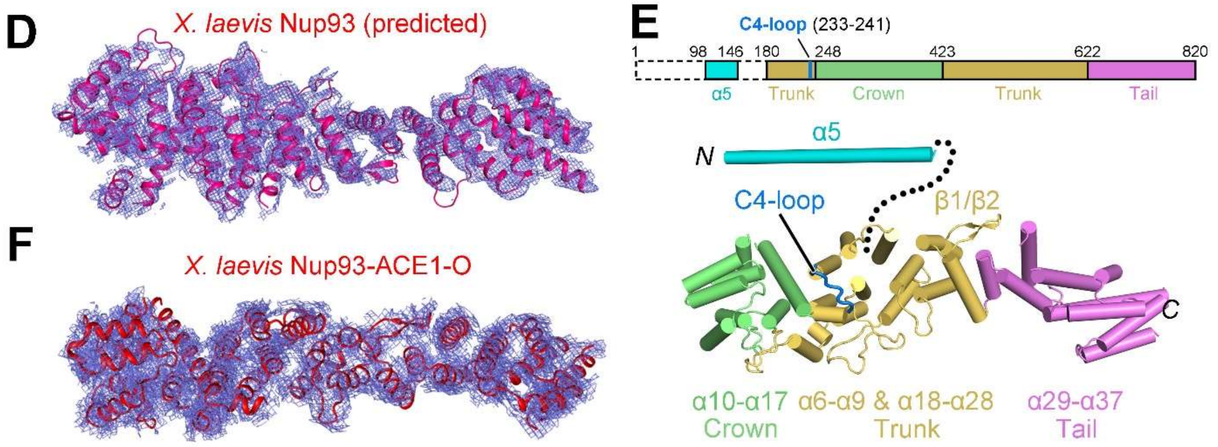


**Fig. S31.**

**Identification of the Bridge domain as Nup93.** (**A**) Superimposition of the poly-Ala model of the Bridge domain and the predicted structure of human Nup93. The predicted structure of human Nup93 comes from the database released by DeepMind (*50*). (**B**) Docking of the predicted structure of human Nup93 into the EM map of the Bridge domain. (**C**) Mass spectrometric analysis of the crosslinked *X. laevis* NE supports the conclusion that Nup93 is the Bridge domain. Shown here are results of mass spectrometric analysis following chemical crosslinking of *X. laevis* NPC using glutaraldehyde. (**D**) Docking of the Alphafold2-predicted structure of *X. laevis* Nup93 (*14*) into the EM map of the Bridge domain. The EM map is derived from the 4.7-Å reconstruction for the Nup358 region at a contour of 5 σ. (**E**) Domain structure of *X. laevis* Nup93. (**F**) Docking of the model of Nup93-ACE1-O into the EM map of the second copy of Nup93-ACE1-I. The local density is prepared with the local refined map of the Nup93-ACE1-I region contoured at 6 σ.

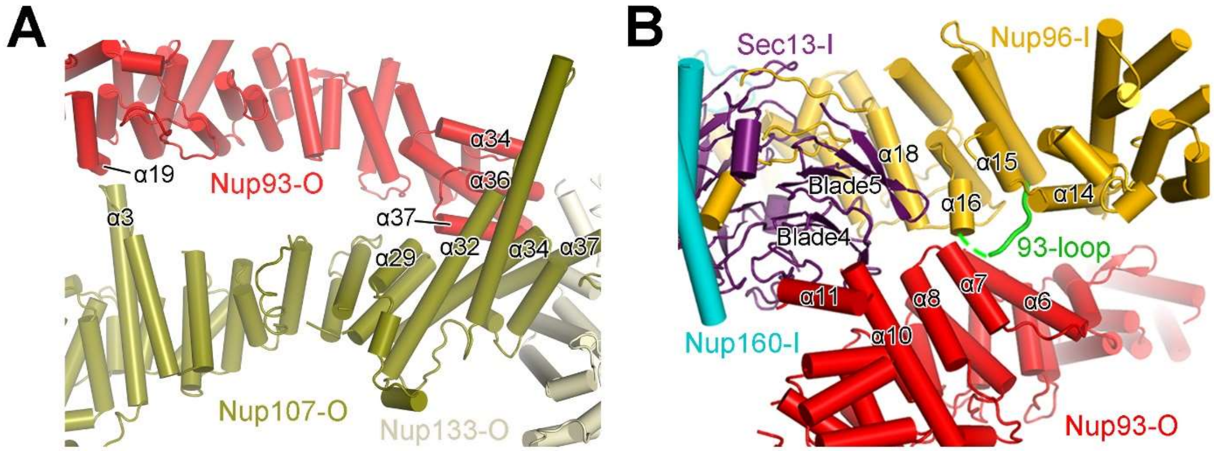


**Fig. S32.**

**Association of Nup93-ACE1-O with other ACE1 proteins in the Y complex.** (**A**) The interface of Nup93-ACE1-O with Nup107-O. On one end, α19 from Nup93-ACE1-O interacts with α3 from Nup107-O. On the other end, helices α34/36/37 from the Tail of Nup93-ACE1-O closely associate with helices α29/32/34/37 of the Tail of Nup107-O. (**B**) The interface between Nup93-O and Nup96-I/Sec13-I. A conserved loop between α15 and α16 of Nup96-I (referred to as 93-loop) interacts with the helical ridge of Nup93-O though binding to α6/7 of Nup93-ACE1-O. The Nup93-Nup96 association appears to be strengthened by Sec13-I, which co-folds with Nup96-I and uses its blades 4/5 to contact helices α11/α10 from the Crown of Nup93-O.

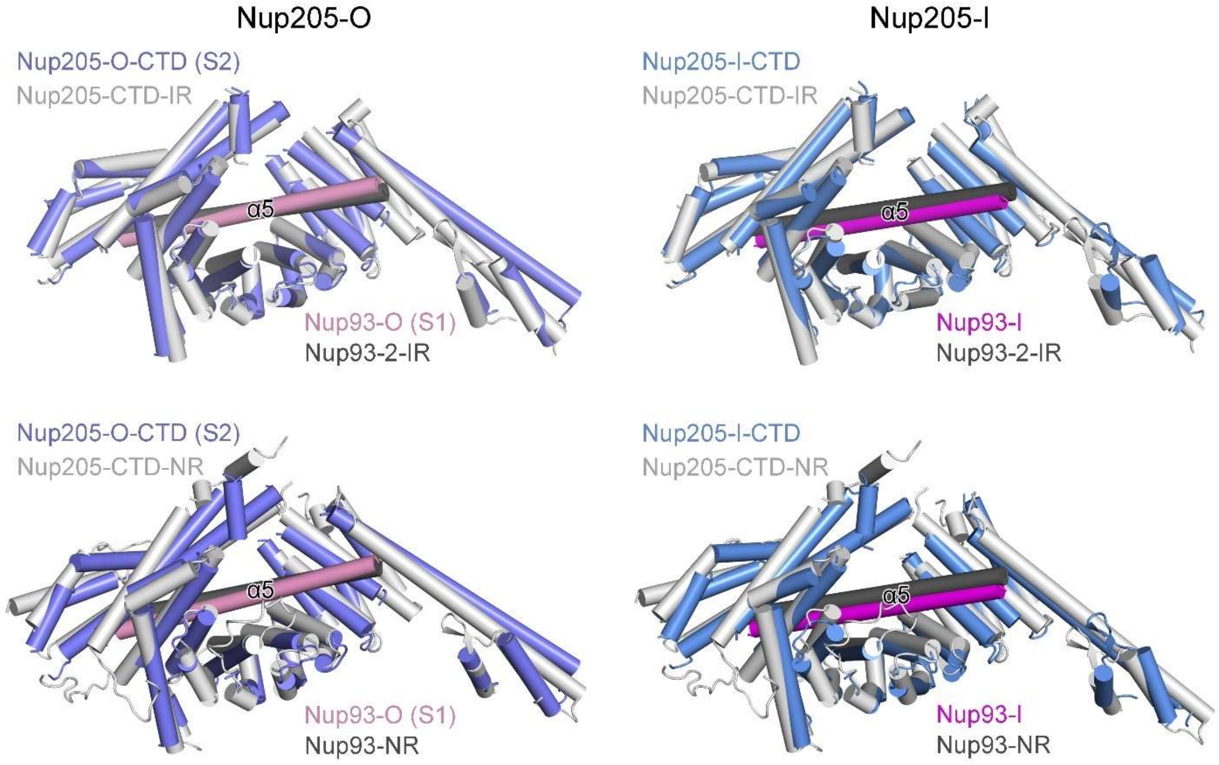


**Fig. S33.**

**A conserved interaction between Nup93 and Nup205.** Shown here are structural comparisons of the *X. laevis* Nup205-CTD (including Tower helix)/Nup93 pairs from the CR subunit with those from the IR and NR subunits. Nup205-O-CTD (S2)/Nup93-O (S1) from the CR can be superimposed with Nup205-CTD/Nup93-2 from the IR and Nup205-CTD/Nup93 from the NR with RMSD of 2.3 Å and 3.4 Å over 571 and 565 aligned Cα atoms, respectively. Nup205-I-CTD/Nup93-I from the CR can be superimposed with Nup205-CTD/Nup93-2 from the IR and Nup205-CTD/Nup93 from the NR with RMSD of 2.0 Å and 3.4 Å over 572 and 565 aligned Cα atoms, respectively.

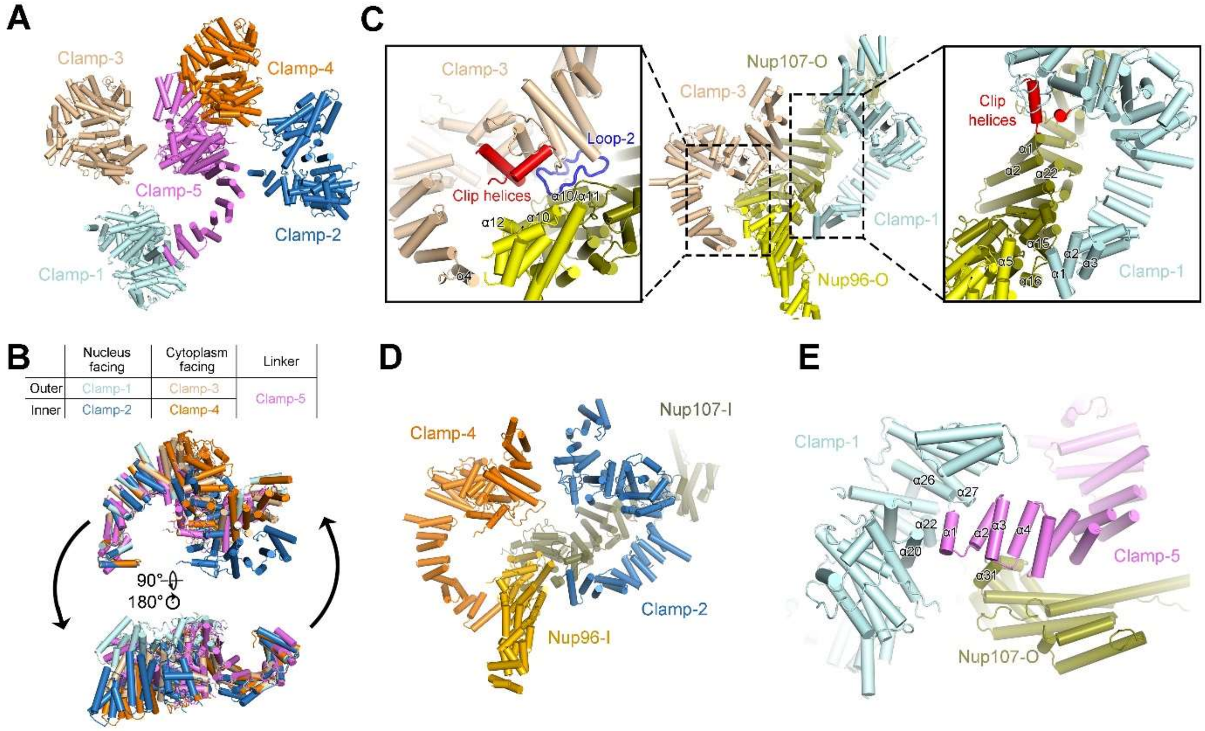


**Fig. S34.**

**Structural analysis of the Nup358 clamps.** (**A**) An overview of the organization of the five Nup358 clamps in the CR subunit. (**B**) Conformational variation of the five Nup358 clamps. With their respective NTD1 aligned, the NTD-2 of Nup358 clamps exhibits apparent conformational changes indicated by the black arrows in the lower panel. (**C**) Clamps-1 and -3 clench the outer Y complex. These interfaces are generally conserved between Clamps-2/-4 and inner Y complex. For Clamp-1, the Clip helices α24/25 are in close contact with the ridge of helices α1/2/22 in the Trunk of Nup107-O, and the N-terminal helices α1/2/3 interact with α15/16 from Nup107-O and α5 from Nup96-O. In contrast, for Clamp-3, the Clip helices α24/25 directly contact helices α10/12 of Nup96-O and Loop-2 interacts with helices α10/α11 of Nup107-O. (**D**) Clamps-2 and -4 clench the inner Y complex. (**E**) Clamp-5 associates with Clamp-1 and grips the outer Y complex through its interface with Nup107-O. At the N-terminus of Clamp-5, helices α1 and α2 contact helices α20/22/26 and α27 from Clamp-1, respectively; helices α3/4 are in close proximity to helix α31 from Nup107-O

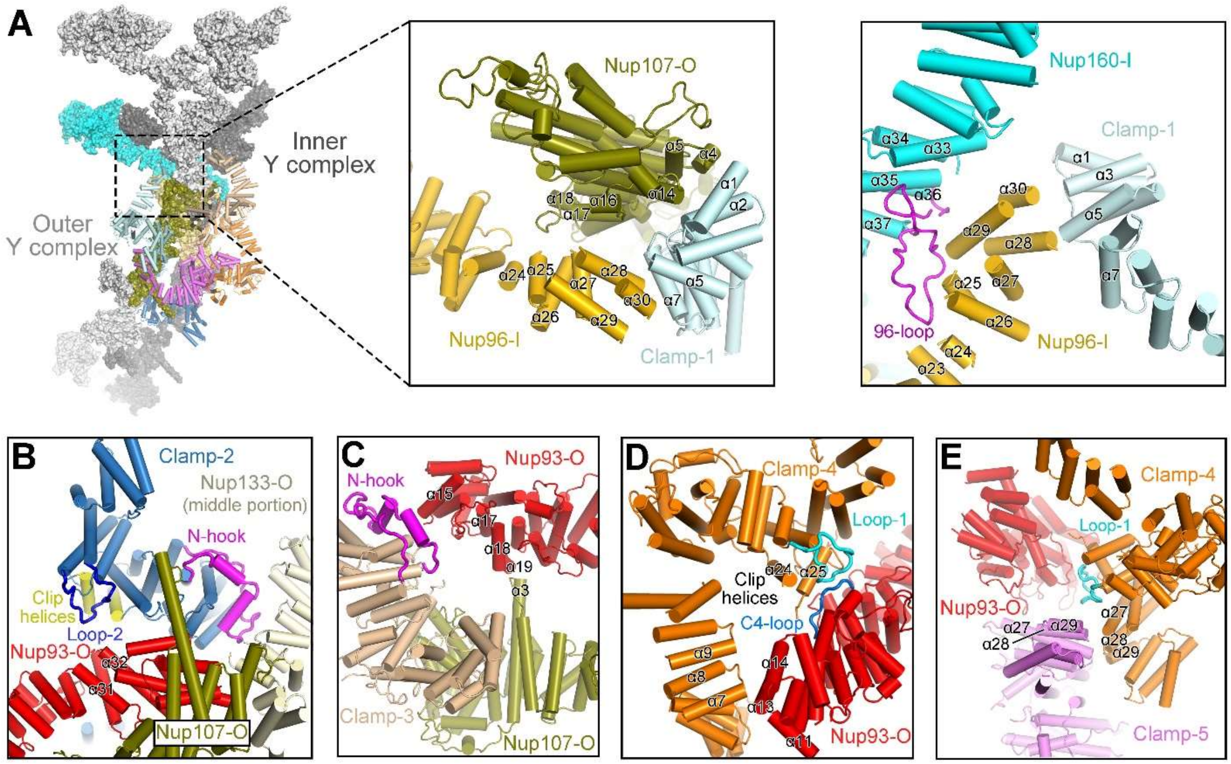


**Fig. S35.**

Additional interfaces between the two Y complexes within the same CR subunit identified in this study. (**A**) Clamp1 and Nup107-O clench the Tail of Nup96-I. Helices α16/17/18 of Nup107-O associates with helices α24/25/27/28 of Nup96-I. Helices α3/α5 of Clamp-1 stack against helices α28/α30 from Nup96-I. (**B**) Interfaces between Clamp-2 and Nup93-ACE1-O, Nup107-O and Nup133-O. Loop-2 and Clip helices of Clamp-2 associate with the ridge of helices α31/α32 from Nup93-O. The N-hook of Clamp-2 interacts with the ridges of helices from the middle portion of Nup133-O. In addition, helices α15/23 of Clamp-2 contact α34 from the Tail of Nup93-ACE1-O. (**C**) Interfaces between Clamp-3 and Nup93-ACE1-O. Helix α15 and the loop between α17 and α18 from the Crown of Nup93-ACE1-O interact with the N-hook of Clamp-3, whereas helix α19 of Nup93-ACE1-O contacts helix α3 of Nup107-O. (**D**) The interfaces between Clamp-4 and the Nup93-ACE1-O. At the Crown of Nup93-ACE1-O, helices α11/13/14 associate with helices α7/8/9 from the NTD1 of Clamp-4. At the Trunk of Nup93-ACE1-O, a flexible loop (named C4-loop, residues 233-241) contacts the Loop-1 and Clip helices of Clamp-4. (**E**) The interface of Clamp-5 with Nup93-O and Clamp-4. At the C-terminus of Clamp-5, helices α27-α29 associate with α27-α29 from Clamp-4 and are in close proximity to the Trunk of Nup93-O. These interfaces bridge clamps-1,3,5 that clench the outer Y complex with clamps-2,4 that clench the inner Y complex.

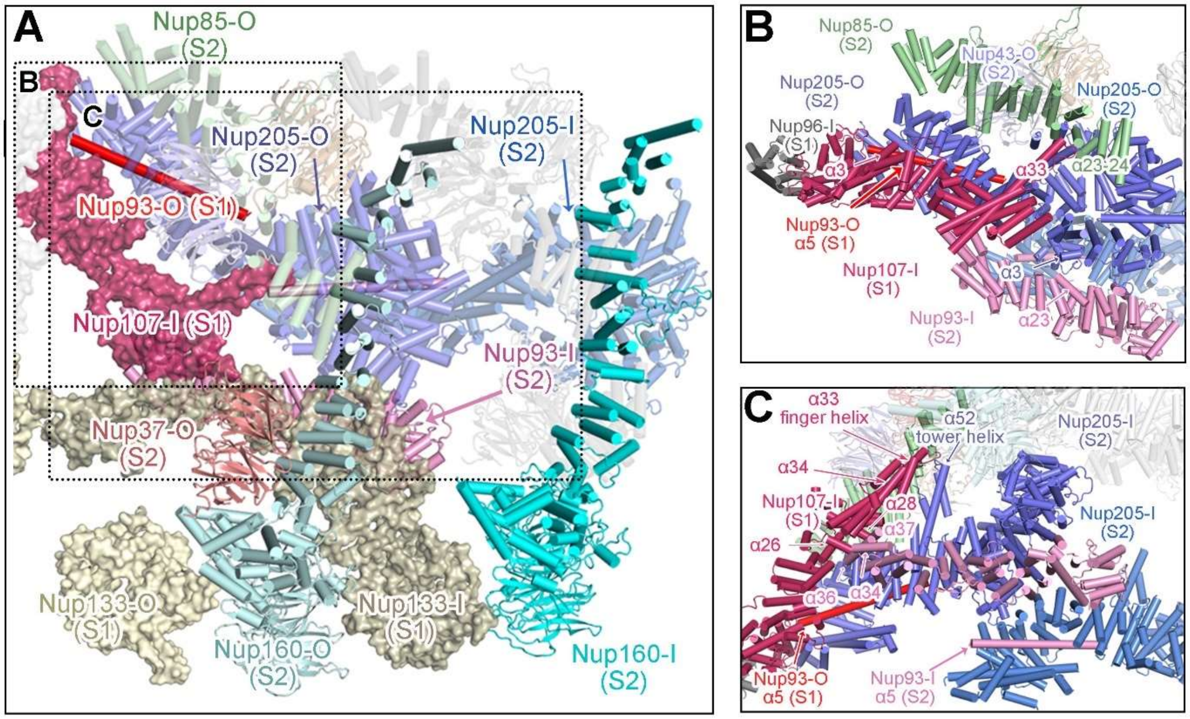


**Fig. S36.**

**Nup205-O plays a central role in the inter-subunit association.** (**A**) Interfaces between adjacent subunits, designated S1 and S2. On one hand, Nup205-O (S2) interacts with the α-solenoid of Nup107-I (S1) and holds the α-solenoid from Nup133-I (S1) with Nup160-O (S2). On the other hand, CTD from Nup205-O (S2) grasps the extended N-terminal helix α5 from Nup93-O (S1). In addition, the N-terminal β-propellers of Nup133 in S1 directly contacts those of Nup160 in S2. (**B**) Interfaces between Nup205-O (S2), Nup93-ACE1-I (S2), and Nup107-I (S1). At the N-terminus of Nup205, helix α23 Nup93-ACE1-I (S2) contact α3 from the NTD of Nup205-O (S2). Around the middle of Nup205-O (S2), the extend finger helix (α33) of Nup107-I (S1) reaches out to contact the long tower helix (α52) of Nup205-O (S2) and the CTD of Nup85-O (S2). The CTD of Nup205-O (S2) interacts with helices α3/α14 in the crown of Nup107-I (S1). (**C**) Interfaces between Nup205 molecules and Nup93 molecules. In addition to the insertion of the Nup93-α5 to the CTD of Nup205, helices α34/36/37 from the Tail of Nup93-ACE1-I associate with the lateral side of helices α26/28 from the Tail of Nup107-I (S1), whereas the Trunk of Nup93-ACE1-I interacts with Nup205-O (S2) NTD.

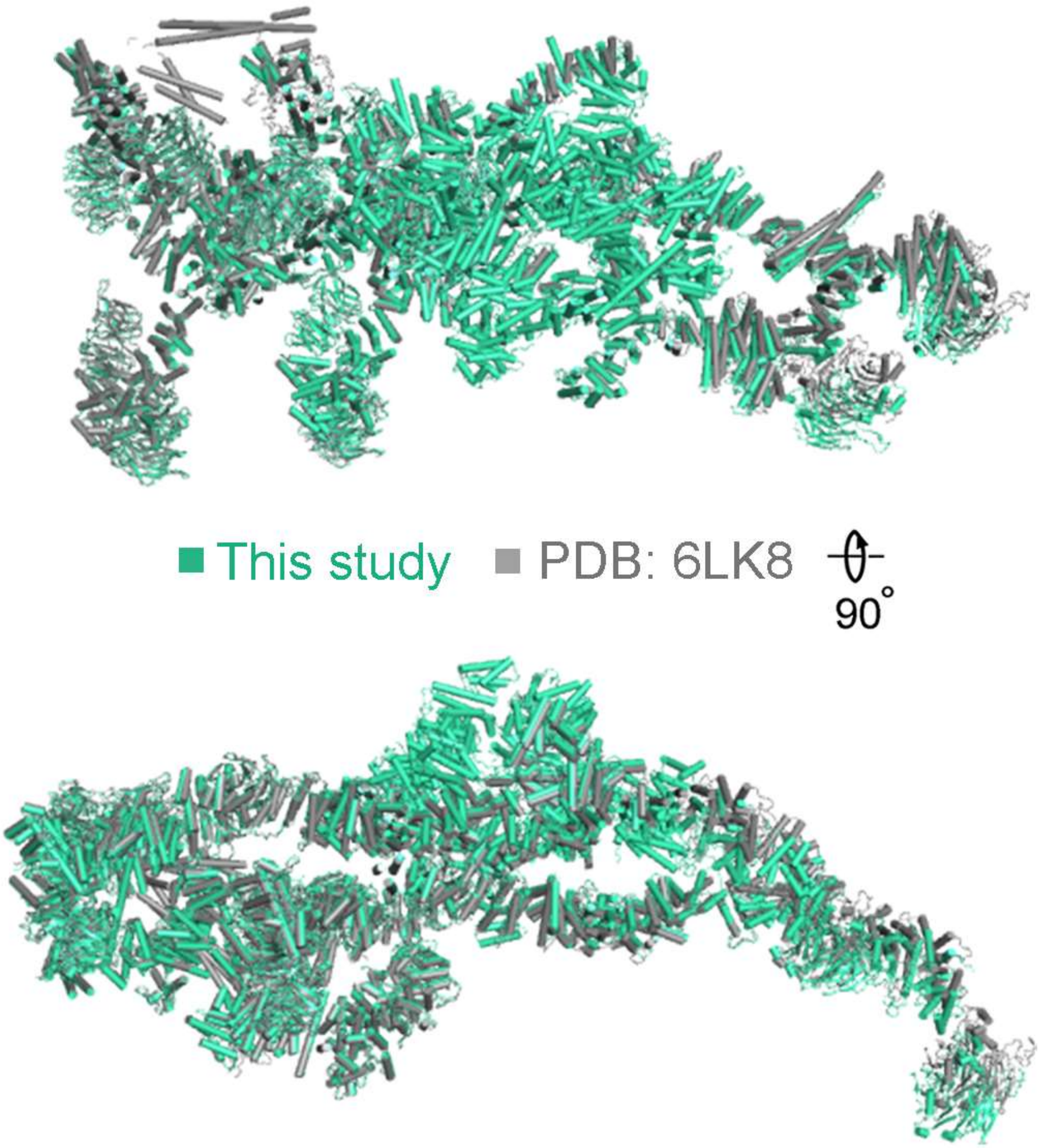


**Fig. S37.**

**Structural comparison of the atomic models in this study with our previous one (*13*).** Superimposition of the atomic models in our previous and present studies are shown in two perpendicular views. The RMSD is approximately 3.7 Å over 10,390 aligned Cα atoms.

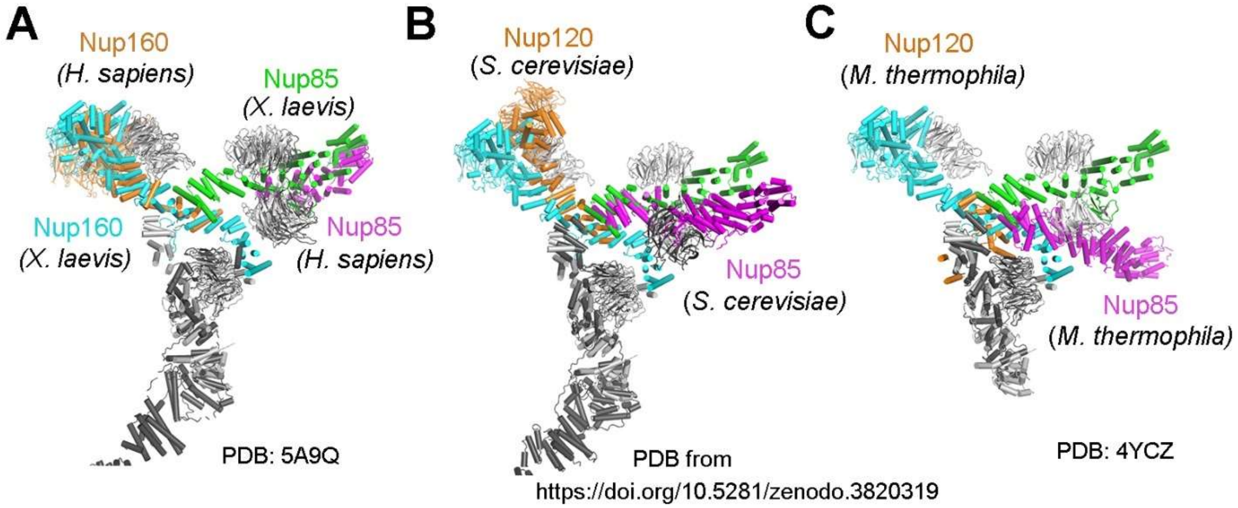


**Fig. S38.**

**Structural comparison of Y complexes from different species.** (*A*) Structural comparison of the Y complexes in this study and that in the *H. sapiens* NPC (*32*). Atomic coordinates of the inner Y complex from *X. laevis* are superimposed with the composite coordinates of the Y complex from *H. sapiens*. The overall conformation and features of the Y complexes remain generally similar between these two structures. The main differences are twofold. First, Nup160-CTF is resolved only in the *X. laevis* EM reconstruction. Second, the top and bottom faces of the Nup43 β-propeller in the *X. laevis* Y complex are opposite of those in the human Y complex. The two β-propellers Nup37 and Nup43 use their top faces to bind to the α-solenoid domains of the short and long arms, respectively. (*B*) Structural comparison of Y complexes from this study and that from *in situ* study of the *S. cerevisiae* NPC. Atomic coordinates of the inner Y complex from *X. laevis* are superimposed with the composite coordinates of the Y complexes from *S. cerevisiae* (*54*). (*C*) Structural comparison of the Y complexes from this study and the core from the *M. thermophila* Y complex (*17*). An apparent rotation of around 40 degrees of the short arm towards the stem-base is observed in the crystal structure of the *M. thermophila* Y complex core. Notably, both Seh1 and Nup160-CTF are absent in *M. thermophila*.

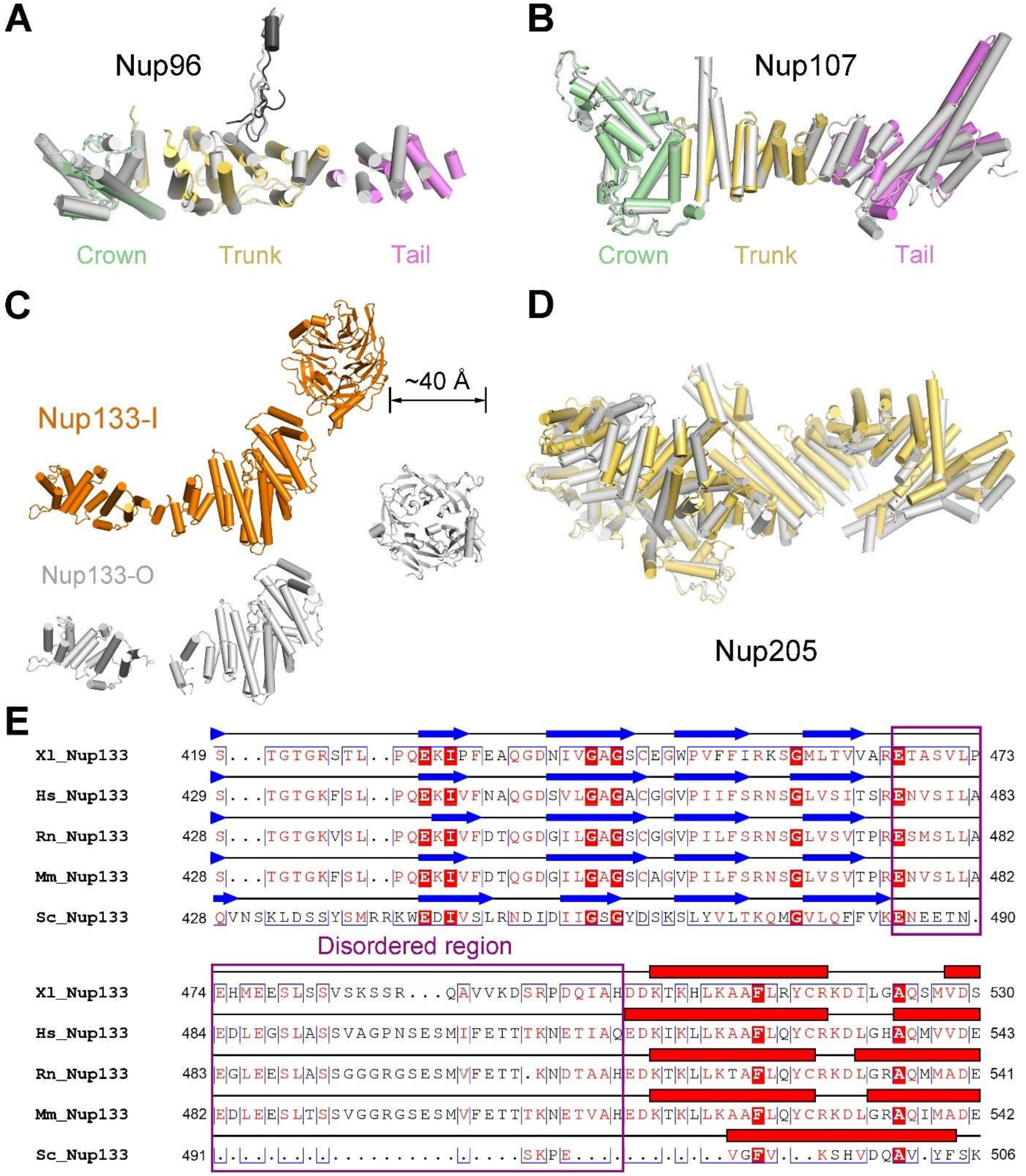


**Fig. S39.**

**Structural comparison of the corresponding components in the two Y complexes.** (**A**) Structural comparison of Nup96-I/O molecules. The Crown, Trunk and Tail of Nup96-I are colored green, yellow and violet, respectively. Nup96-O is shown in gray. The two Nup96 molecules are aligned on their trunk, producing an RMSD of 1.8 Å over 209 aligned Cα atoms. (**B**) Structural comparison of Nup107-I/O molecules. The Crown, Trunk and Tail of Nup107-I complex are colored green, yellow and violet, respectively. Nup107-O shown in gray. The two Nup107 molecules are superimposed relative to their Trunk, producing an RMSD of 0.9 Å over

197 aligned Cα atoms. (**C**) Structural comparison between Nup133-I/O. The N-terminal β-propeller of Nup133-O is shifted by approximately 4 nm against its C-terminal α-solenoid when compared to that of Nup133-I. The two Nup133 molecules are aligned on their C-terminal α-solenoid, producing an RMSD of 2.1 Å over 526 aligned Cα atoms. (**D**) Structural alignment between Nup205-I (yellow) and Nup205-O (gray). The two Nup205 molecules are aligned on their NTD, producing an RMSD of 3.5 Å over 650 aligned Cα atoms. (**E**) Sequence alignment for the disordered region linking the N-terminal β-propeller and α-helical domain in Nup133. The predicted disordered region is absent in *Saccharomyces cerevisiae* Nup133. Xl, *Xenopus laevis*; Hs, *Homo sapiens*; Rn, *Rattus norvegicus*; Mm, *Mus musculus* and Sc, *Saccharomyces cerevisiae*.

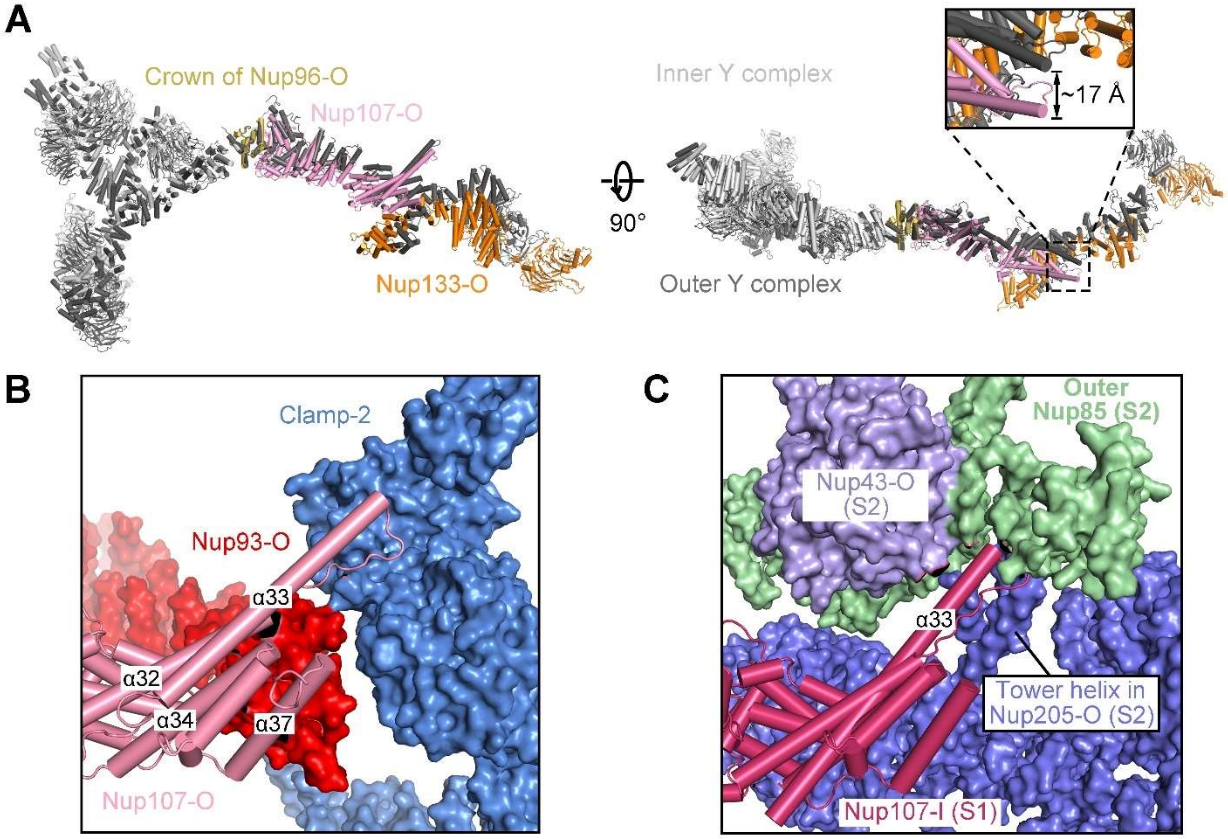


**Fig. S40.**

**Structural comparison between the inner and outer Y complexes.** (**A**) Structural alignment of the inner and outer Y complexes on the Crowns of Nup96-I/O (PDB from this study). The inner and outer Y complexes are shown in light and dark gray, respectively. Components of the stem are color-coded in the outer Y complex. (**B**) Local environment around the finger helix from Nup107-O. (**C**) Local environment around the finger helix from Nup107-I (S1).

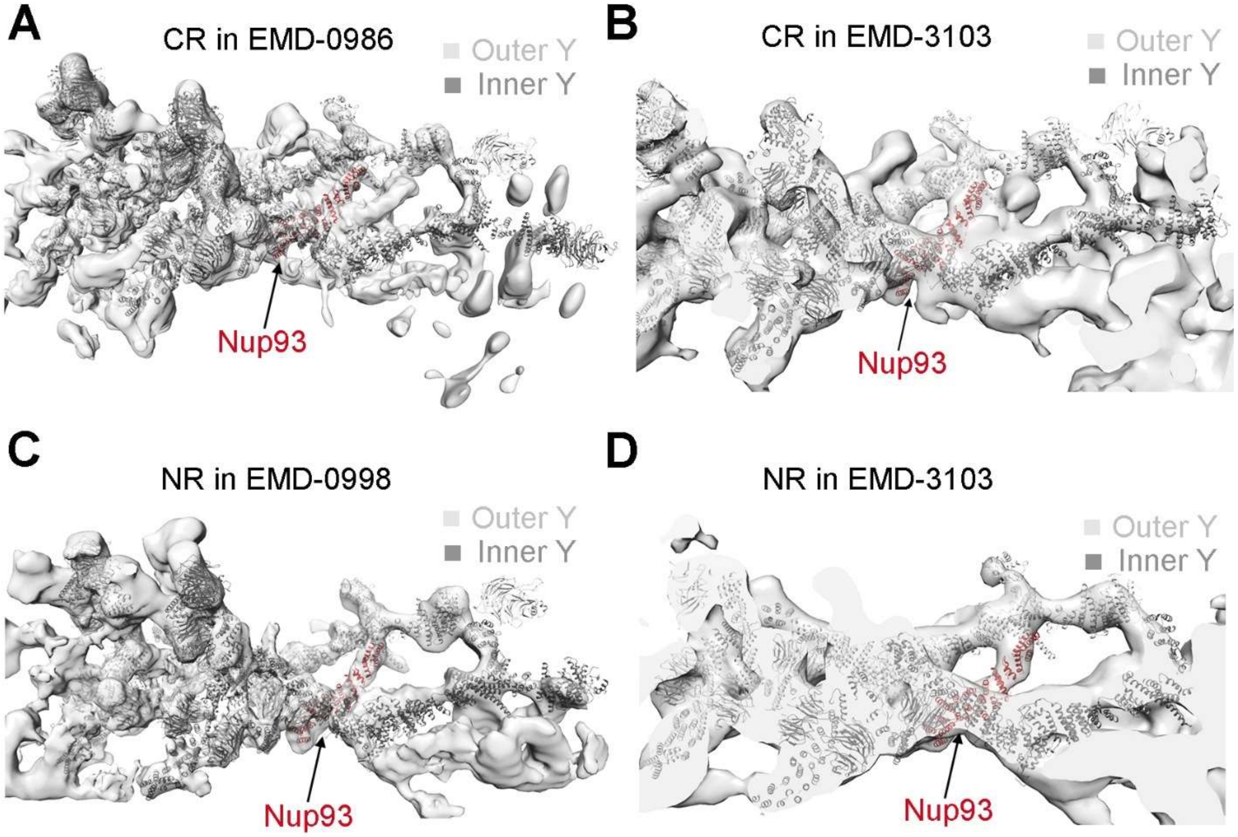


**Fig. S41.**

**Proposed presence of Nup93 in NR.** (**A**) The atomic model of Nup93-O from *X. laevis* CR subunit can be docked into the EM map (EMD-0986) (*9*) for the *X. laevis* CR subunit. (**B**) The atomic model of Nup93-O from *X. laevis* CR subunit can be docked into the EM map for the human CR subunit (EMD-3103) (*32*). The location and pose of the docked Nup93 molecule are almost identical to those of Nup93-O in *X. laevis* CR subunit. (**C**) The atomic model of Nup93-O from *X. laevis* CR subunit can be docked into the EM map (EMD-0998) (*9*) for the *X. laevis* NR subunit. The docked model of Nup93 resides at a location that is almost identical to that of Nup93-O in the CR subunit, linking the two layers of Y complexes. (**D**) The atomic model of Nup93-O from *X. laevis* CR subunit can be docked into the EM map (EMD-3103) (*32*) for the human NR subunit.

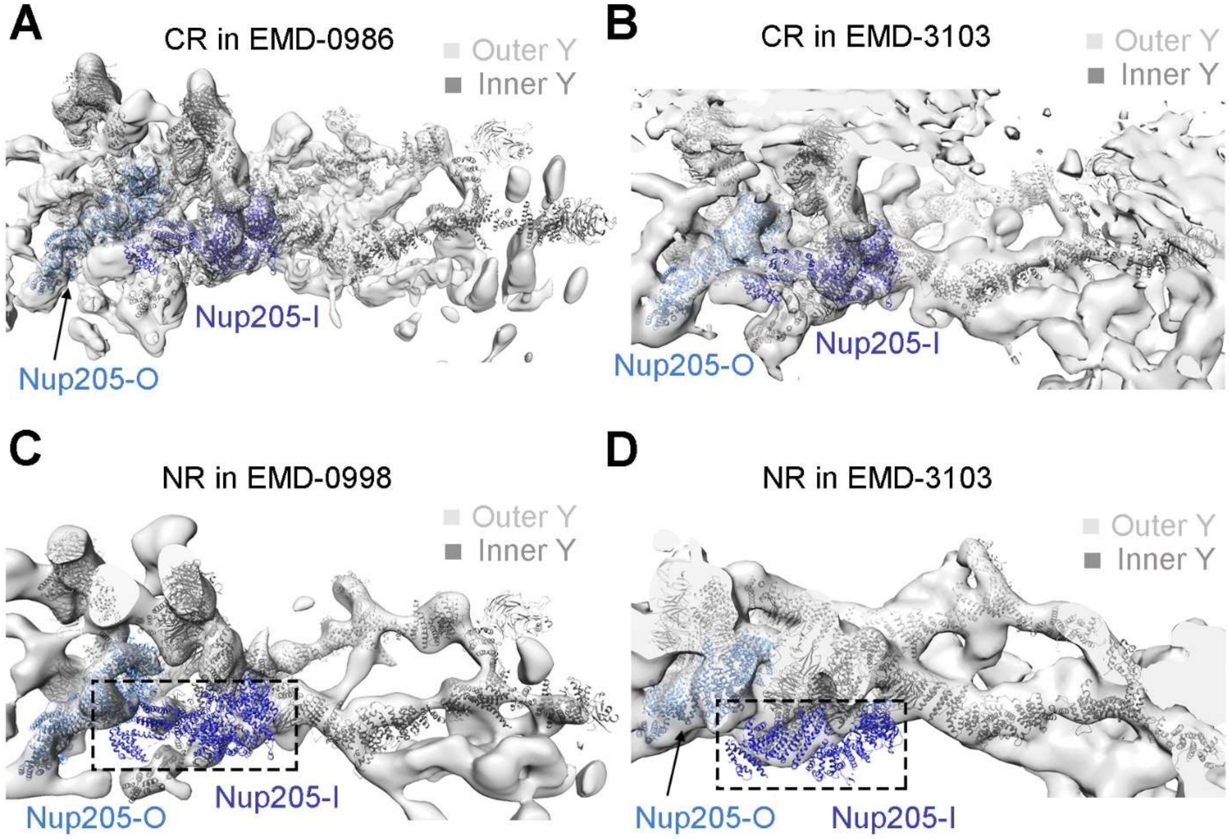


**Fig. S42.**

**Differential placement of Nup205 into the subunits of the CR and the NR.** (**A**) Each CR subunit from the *X. laevis* NPC accommodates two molecules of Nup205. Shown here is the EM map for one CR subunit from the *X. laevis* NPC (EMD-0986) (*9*). (**B**) Each CR subunit from the human NPC can accommodate two molecules of Nup205. Shown here is the EM map for one CR subunit from the human NPC (EMD-3103) (*32*). The docking is reasonable. (**C**) Each NR subunit from the *X. laevis* NPC (EMD-0998) (*9*) only accommodates one molecule of Nup205 (Nup205-O). There is no density corresponding to Nup205-I. (**D**) Each NR subunit from the human NPC (EMD-3103) (*32*) can only accommodate one molecule of Nup205 (Nup205-O). Nup205-I cannot be docked into the map.

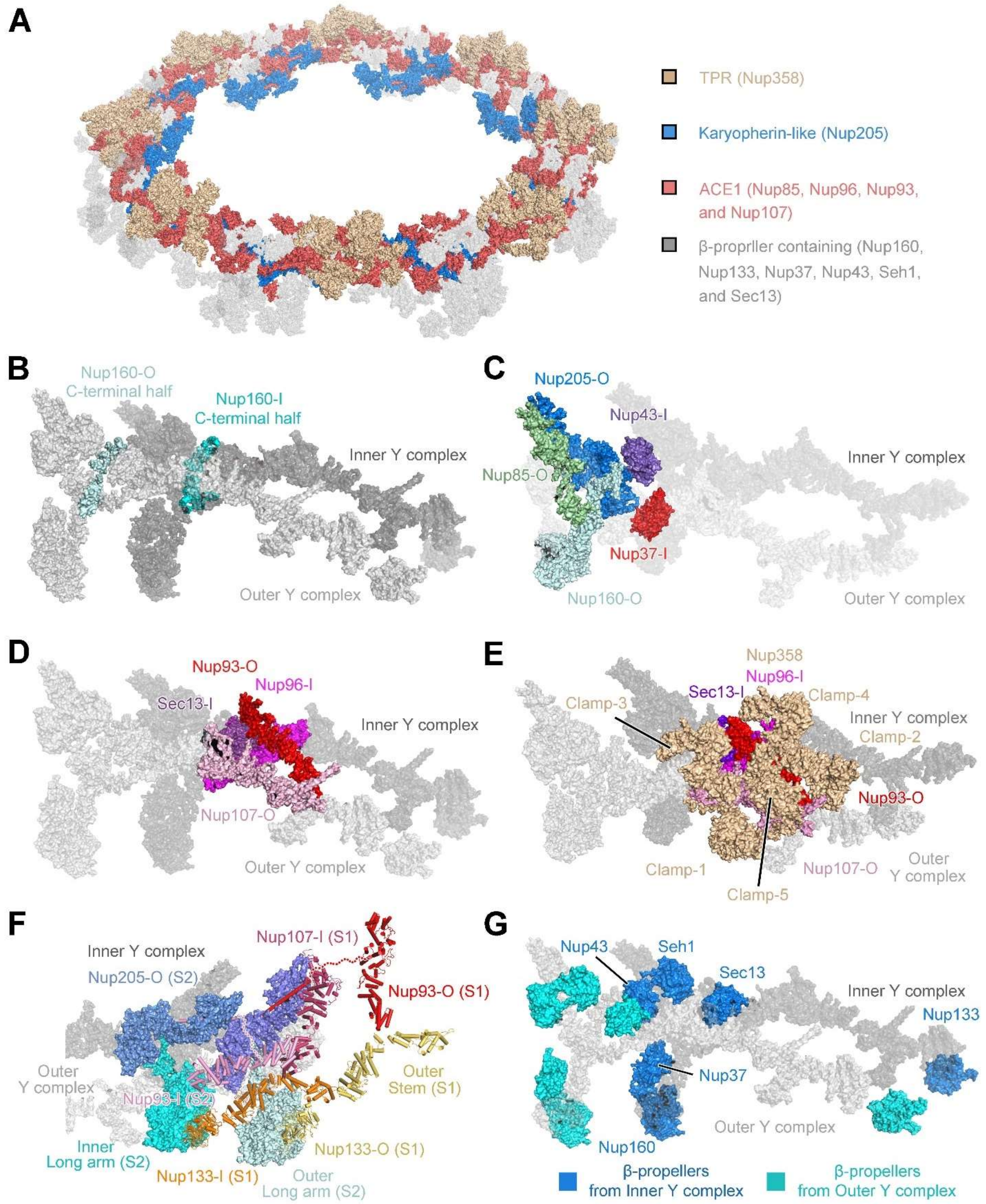


**Fig. S43.**

**A summary of key components and interfaces in the assembly of the CR.** (**A**) Limited number of structural folds in the CR. The ACE1 fold, Karyopherin-like fold and TPR fold α-helical nucleoporins are color coded. The membrane contacting proteins and β-propeller proteins are colored gray. (**B**) The C-terminal half (α32 through α47) of Nup160 at the vertex connects the short arm and the stem base of Y complex by interacting with four nucleoporins (Nup85, Seh1, Nup96, Sec13). (**C**) Nup205-O connects both arms of inner Y complex with the short arm and the vertex of outer Y complex. Nup205-O directly binds Nup85-O, Nup160-O, Nup37-I, and Nup43-I. (**D**) Nup93-O, Nup107-O, Sec13-I, and Nup96-I form six pairwise interfaces in the stems of the Y complexes. (**E**) Nup358 clamps clench onto the stems of the Y complexes and encage the Nup93 ACE1. (**F**) A close-up view on the inter-subunit interfaces. The stem of inner Y complex (S1) associates with the long arms of both inner and outer Y complexes (S2). Within each of the two concentric rings, the β-propeller of Nup133 (S1) interacts with the long arm of the Y complex from the posterior subunit to form the head-to-tail architecture. Nup93-O from S1 uses its N-terminal extended helix α5 to insert into the axial groove of Nup205-O CTD in S2. Nup93-I (S2) connects the Nup107-I (S1) and the NTD of Nup205-O (S2) using its ACE1 and reaches into the axial groove of Nup205-I CTD (S2) with its extended N-terminal helix α5. (**G**) Twelve β-propellers serve as bolts and rivets to fix the elastic α-solenoids of the nucleoporins in two Y complexes.

| | CR subunit (7FIK) (EMD-36000) | Nup358 region (7FIK) (EMD-36001) | Nup358-NTF (7FIL) (EMD-36002) |
|---|---|---|---|
| **Data collection and Processing** | | | |
| Microscope | | Titan Krios (Thermo Fisher Scientific) | |
| Voltage (keV) | | 300 | |
| Camera | | Gatan K3 | |
| Magnification | | 64,000 | |
| Pixel size at detector (Å/pixel) | | 1.387 | |
| Total electron exposure ($e^-/Å^2$) | 50/57/87/100 | | 50 |
| Exposure rate ($e^-$/pixel/sec) | | 19.5 | |
| Number of frames collected during exposure | 32/37/46/56 | | 32 |
| Defocus range (µm) | | -1.0 ~ -4.0 | |
| Automation software | | AutoEMation2 | |
| Tilt angle (°) | 0/30/45/55 | | 0 |
| Energy filter slit width | | 20 eV | |
| Micrographs collected (no.) | 10,040/7,557/14,107/14,439 | | 17,147 |
| Micrographs used (no.) | 8,145/6,171/10,319/9,112 | | 17,147 |
| Total extracted particles (no.) | 5,148,474 | | 2,418,968 |
| **For each reconstruction:** | | | |
| Refined particles (no.) | 1,279,381 | | 452,078 |
| Final particles (no.) | 1,279,381 | | 452,078 |
| Symmetry parameters | | C1 | |
| Estimated error of translations (pixels)/rotations (°) | 1.06/1.42 | 1.39/2.55 | 1.64/3.16 |
| Resolution (global, Å) | | | |
| FSC 0.5 (unmasked/masked) | 4.8/4.3 | 9.6/5.7 | 4.1/3.6 |
| FSC 0.143 (unmasked/masked) | 4.3/3.8 | 7.5/4.6 | 3.3/2.9 |
| Resolution range (local, Å) | 50-3.8 | 50-4.2 | 50-2.8 |
| Resolution range due to anisotropy (Å) | 4.8-4.0 | 7.2-4.4 | 3.1-3.0 |
| Map sharpening *B* factor ($Å^2$) / (B factor Range) | -120 | -125 | -100 |
| Map sharpening methods | | Ad-hoc B-factor | |
| **Model composition** | | | |
| Protein | yes | | yes |
| Ligands | / | | / |
| RNA/DNA | / | | / |
| **Model Refinement (for each model)** | | | |
| Refinement package | | Phenix | |
| - real or reciprocal space | | real | |
| - resolution cutoff | 4.1 | 4.7 | 3.0 |
| Model-Map scores | | | |
| -CC | 0.70 | 0.71 | 0.75 |
| - Average FSC | 4.0 | 5.6 | 4.0 |
| *B* factors ($Å^2$) | 174.49 | | 99.94 |
| Protein residues | 19,037 | | 537 |
| Ligands | / | | / |
| RNA/DNA | / | | / |
| R.m.s. deviations from ideal values | | | |
| Bond lengths (Å) | 0.006 | | 0.004 |
| Bond angles (°) | 0.852 | | 0.687 |
| **Validation** | | | |
| MolProbity score | 2.23 | | 2.77 |
| CaBLAM outliers | 4.3% | | 3.1% |

| | | |
|---|---|---|
| Clashscore | 12.25 | 11.41 |
| Poor rotamers (%) | 1.19 | 7.80 |
| C-beta deviations | 0 | 0 |
| Ramachandran plot | | |
| Favored (%) | 89.06 | 90.93 |
| Outliers (%) | 0.43 | 0 |

**Table S1.**

**Cryo-EM data collection, refinement, and validation statistics.**

| | Molecule **Vertebrates**/ Yeast | Copy No. | | Length *Xenopus* | UniProt No. *Xenopus* | PDB code | Modeling | Model length | Resolution (Å) | Chain ID Outer/Inner |
|---|---|---|---|---|---|---|---|---|---|---|
| Y complex | **Nup85**/Nup85 | 2 | | 653 | Q68FJ0 | 3F3F, 4XMM | HM | ~600 | 4.0~4.5 | B/b |
| | **Nup160**/Nup120 | 2 | | 1435 | A0A1L8GIX3 | 4GQ2, 4XMM | HM | ~1100 | 4.0~4.5 | E/e |
| | | | 160-CTF | - | - | - | DM/AF | ~230 | | |
| | **Nup96**/Nup145C | 2 | | 923 | A0A1L8HBE3 | 3IKO, 4XMM | HM | ~510 | 4.0~4.5 | G/g |
| | **Nup107**/Nup84 | 2 | | 916 | A2RV69 | 3IKO, 3I4R | HM | ~770 | 4.5~6.0 | I/i |
| | **Nup133**/Nup133 | 2 | | 1140 | A0A1L8H1I9 | 1XKS, 3I4R | HM | ~1000 | 4.5~6.0 | J/j |
| | **Sec13**/Sec13 | 2 | | 320 | Q7ZYJ8 | 3BG1 | HM | ~280 | 4.0~4.5 | H/h |
| | **Seh1**/Seh1 | 2 | | 360 | Q4FZW5 | 3F3F | HM | ~310 | 4.0~4.5 | D/d |
| | **Nup43**/- | 2 | | 375 | Q05AW3 | 4I79 | HM | ~370 | 4.0~4.5 | C/c |
| | **Nup37**/Nup37 | 2 | | 326 | Q66IZ6 | 4GQ2 | HM | ~320 | 4.0~6.0 | F/f |
| **Nup358** | **Nup358**/ - | 5 | Clamp-1/2/3/4/5 | 2905 | A0A1L8HGL2 | - | HM/RD | ~740 | 4.5~8.0 | K/L/M/N/O |
| **Nup93** | **Nup93**/Nic96 | 2 | | 820 | Q7ZX96 | - | HM/AF | ~590 | 4.5~6.5 | P/Q |
| | **Nup93 (S2)** | - | | 820 | Q7ZX96 | - | AF | ~50 | 5.0~6.5 | R |
| **Nup205** | **Nup205**/Nup192 | 2 | | 2011 | Q642R6 | 5HB4 | HM/RD | ~1770 | 4.0~5.5 | A/a |
| **Nup88** | **Nup88**/Nup82 | 1 | | 728 | Q4KLQ6 | - | AF/RD | ~400 | 6.0~8.0 | S |
| **Nup155** | **Nup155**/Nup170, Nup157 | 1 | | 1388 | F6UHT0 | - | AF/RD | ~270 | 5.0~7.0 | T |
| **Nup98/X** | **Nup98/X** | 1 | | 819 | A0A1L8HBE3 | - | AF/RD | ~150 | 6.0~8.0 | X |

**Table S2.**

**Summary of model building for the CR subunit of *X. laevis* NPC.** Under the column labeled "Molecule", proteins from vertebrates and yeasts are shown, respectively, with vertebrate components in bold. Under the column labeled "Modeling", HM stands for homology modelling; DM stands for *de novo* modelling; AF stands for manual adjustment based on the AlphaFold model; RD stands for rigid docking.

| Protein | Full length (aa) | Residues modeled | | Number of α-helices | | Number of β-strands | |
|---|---|---|---|---|---|---|---|
| | | Inner | Outer | Inner | Outer | Inner | Outer |
| Nup160 | 1435 | 1259 | 1195 | 47 | 46 | 29 | 29 |
| Nup37 | 326 | 291 | 287 | 0 | 0 | 28 | 28 |
| Nup85 | 653 | 533 | 539 | 27 | 27 | 3 | 3 |
| Nup43 | 375 | 300 | 299 | 0 | 0 | 28 | 28 |
| Seh1 | 360 | 282 | 307 | 0 | 0 | 24 | 24 |
| Nup96 | 923 | 493 | 499 | 30 | 30 | 3 | 3 |
| Sec13 | 320 | 286 | 291 | 1 | 1 | 24 | 24 |
| Nup107 | 916 | 726 | 765 | 37 | 37 | 0 | 0 |
| Nup133 | 1140 | 1028 | 1021 | 33 | 33 | 28 | 28 |
| Nup205 | 2011 | 1604 | 1569 | 78 | 77 | 2 | 0 |
| Nup93 | 820 | 607 | 644 | 33 | 33 | 2 | 2 |

| Protein | Full length (aa) | Residues modeled* | | | | | Number of α-helices/β-strands | | | | |
|---|---|---|---|---|---|---|---|---|---|---|---|
| | | C1 | C2 | C3 | C4 | C5 | C1 | C2 | C3 | C4 | C5 |
| Nup358 | 2905 | 703 | 699 | 654 | 704 | 683 | 32/0 | 32/0 | 29/0 | 32/0 | 32/0 |

| Protein | Full length (aa) | Residues modeled | Number of α-helices/β-strands |
|---|---|---|---|
| Nup155 | 1388 | 269 | 15/0 |
| Nup88 | 728 | 339 | 2/29 |
| Nup98/X | 819 | 161 | 5/11 |

#### Table S3.

**Summary of secondary structural elements in the structurally resolved nucleoporins of the core and Nup358 regions.** *C1~C5: Clamp-1~Clamp-5.

| Inner Y | Outer Y | Full length | Optimal alignment (by PyMol) | |
|---|---|---|---|---|
| | | | RMSD (Å) | No. of aligned Cα |
| Nup37 (291) | Nup37 (287) | 326 | 0.829 | 260 |
| Nup43 (300) | Nup43 (299) | 375 | 0.916 | 279 |
| Sec13 (286) | Sec13 (291) | 320 | 0.729 | 271 |
| Seh1 (282) | Seh1 (307) | 360 | 0.682 | 244 |
| Nup85 (533) | Nup85 (539) | 653 | 3.815 | 379 |
| Nup96 (493) | Nup96 (499) | 923 | 2.431 | 431 |
| Nup160-CTF (152) | Nup160-CTF (166) | 230* | 5.006 | 110 |
| Nup205 (1604) | Nup205 (1569) | 2011 | 3.656 | 1331 |
| Nup107 (726) | Nup107 (765) | 916 | 2.759 | 614 |
| Nup133 β-propeller (398) | Nup133 β-propeller (398) | 1140 | 0.013 | 357 |
| Nup133 α-helical domain (630) | Nup133 α-helical domain (623) | 1140 | 2.108 | 526 |
| Clamp-1 (703) | Clamp-2 (699) | | 2.366 | 543 |
| Clamp-1 (703) | Clamp-3 (654) | | 0.687 | 491 |
| Clamp-1 (703) | Clamp-4 (704) | | 5.567 | 695 |
| Clamp-1 (703) | Clamp-5 (683) | | 4.721 | 614 |
| Clamp-2 (699) | Clamp-3 (654) | | 4.239 | 536 |
| Clamp-2 (699) | Clamp-4 (704) | | 4.087 | 579 |
| Clamp-2 (699) | Clamp-5 (683) | | 7.404 | 600 |
| Clamp-3 (654) | Clamp-4 (704) | | 0.763 | 485 |
| Clamp-3 (654) | Clamp-5 (683) | | 2.705 | 491 |
| Clamp-4 (704) | Clamp-5 (683) | | 5.077 | 562 |

**Table S4.**

**Structural comparison of the CR components.** *Only the C-terminal fragments (consisting of 230 residues) from Nup160-I and Nup160-O were compared.

| No | Chain | Z-score | RMSD (Å) | Aligned residue number | Description |
|---|---|---|---|---|---|
| 1 | 2qx5-A | 14.3 | 8.1 | 400 | Nucleoporin Nic96 |
| 2 | 5ijn-C | 12.1 | 4.3 | 309 | Nuclear pore complex protein Nup93 |
| 3 | 2rfo-A | 10.9 | 4.3 | 314 | Nucleoporin NIC96 |
| 4 | 6x07-A | 10.7 | 4.8 | 306 | Nucleoporin NIC96 |
| 5 | 5hb3-C | 9.9 | 5.5 | 305 | Nucleoporin NIC96 |
| 6 | 4xmm-F | 7.9 | 4.6 | 205 | Nucleoporin NUP84 |
| 7 | 6x02-A | 7.4 | 4.8 | 337 | Nucleoporin NUP84 |
| 8 | 3iko-I | 7.3 | 4.4 | 200 | Nucleoporin NUP84 |
| 9 | 3iko-B | 7.2 | 6.1 | 231 | Nucleoporin NUP145C |
| 10 | 3jro-A | 7.1 | 6.7 | 234 | Fusion Protein of Protein SEC13 and Nucleoporin NUP145 |
| 11 | 6lk8-B | 7.0 | 8.3 | 250 | Nuclear pore complex protein Nup85 |
| 12 | 4xmn-B | 6.7 | 7.3 | 277 | Nucleoporin NUP145C |
| 13 | 4xmn-F | 6.7 | 4.6 | 199 | Nucleoporin NUP84 |
| 14 | 4xmn-B | 6.5 | 8.2 | 294 | Nucleoporin NUP145C |
| 15 | 3bg0-B | 6.2 | 6.8 | 237 | Nucleoporin NUP145C |
| 16 | 1dce-A | 6.0 | 6.7 | 96 | RAB Geranylgeranyltransferase alpha subunit |
| 17 | 4ycz-B | 5.8 | 8.5 | 238 | Fusion protein of SEC13 and NUP145C |
| 18 | 3f3f-D | 5.7 | 8.6 | 218 | Nucleoporin NUP85 |
| 19 | 3ewe-B | 5.2 | 7.7 | 205 | Nucleoporin NUP85 |
| 20 | 6x03-A | 5.2 | 7.7 | 257 | Nucleoporin NUP84 |

**Table S5.**

**Statistics of the Dali-search results of the Bridge domain.** The results are generated from the Dali server (*49*) using the PDB-search option and the list is sorted by Z-score. Shown here are the top 20 neighbors of the initial poly-Ala Bridge domain.

| **Within the same CR subunit** | | | | **Between two CR subunits** | |
|---|---|---|---|---|---|
| **Within the same Y complex** | | **Between two Y complexes** | | Subunit-1 | Subunit-2 |
| Subject-1 | Subject-2 | Subject-1 | Subject-2 | | |
| Nup85 | Seh1 | Nup96-I | Nup107-O | Nup133-O (β-propeller) | Nup160-O-NTD |
| Nup85 | Nup43 | Sec13-I | Nup107-O | Nup133-I (α-helical domain) | Nup160-O-NTD |
| Nup85-CTD | Nup160-CTD | Nup133-I | Nup133-O | Nup133-I (α-helical domain) | Nup37-O |
| Nup160 (96-loop) | Nup96-CTD | **Intra-CR subunit (except inter-Y complex)** | | Nup133-I | Nup205-O |
| Nup160 | Nup37 | Subject-1 | Subject-2 | Nup107-I Finger helix | Nup85-O |
| Nup160-CTF | Sec13 | Nup205-O Tower helix | Nup85-O | Nup107-I Finger helix | Nup205-O Tower helix |
| Nup160-CTF | Nup96-NTD | Nup205-NTD | Seh1 | Nup107-I | Nup43-O |
| Nup160-CTF | Seh1 | Nup205-O-TAIL | Nup85-O | Nup107-I | Nup205-O CTD |
| Nup96 | Sec13 | Nup205-O-NTD | Nup160-O-CTF | Nup93-O | Nup205-O |
| Nup96 | Nup107 | Nup205-O-NTD | Nup43-I (bottom) | | |
| Nup107 | Nup133 | Nup205-O-NTD | Nup37-I (bottom) | | |
| | | Clamp-1 | Nup107-O | | |
| | | Clamp-1 | Nup96-I | | |
| | | Clamp-1 | Clamp-5 | | |
| | | Clamp-3 | Nup96-O | | |
| | | Clamp-3 | Nup107-O | | |
| | | Clamp-4 | Nup96-I | | |
| | | Clamp-4 | Nup93-O | | |
| | | Clamp-4 | Clamp-5 | | |
| | | Clamp-5 | Nup107-O | | |
| | | Clamp-5 | Nup93-O | | |
| | | Nup93-O | Clamp-3 | | |
| | | Nup93-O | Nup107-O | | |
| | | Nup93-O | Nup96-I | | |
| | | Nup93-O | Sec13-I | | |
| | | Nup93-I | Nup205-I | | |

**Table S6.**

**Summary of protein-protein interfaces in the CR.**
